# RORγ bridges cancer-driven lipid dysmetabolism and myeloid immunosuppression

**DOI:** 10.1101/2023.11.19.567414

**Authors:** Augusto Bleve, Francesca Maria Consonni, Martina Incerti, Valentina Garlatti, Chiara Pandolfo, Marta Noemi Monari, Simone Serio, Daniela Pistillo, Marina Sironi, Chiara Alì, Marcello Manfredi, Giovanna Finocchiaro, Cristina Panico, Gianluigi Condorelli, Antonio Sica

**Affiliations:** Department of Pharmaceutical Sciences, University of Piemonte Orientale “A. Avogadro”, Novara, Italy; IRCCS Humanitas Research Hospital, Rozzano, Milan, Italy; Laboratory of Clinical Analysis, IRCCS Humanitas Research Hospital, Rozzano, Milan, Italy; Department of Biomedical Sciences, Humanitas University, Pieve Emanuele, Milan, Italy; Department of Cardiovascular Medicine, IRCCS Humanitas Research Hospital, Rozzano, Milan, Italy; Biobank, Humanitas Cancer Center, IRCCS Humanitas Research Hospital, Rozzano, Milan, Italy; Department of Translational Medicine, University of Piemonte Orientale, Novara, Italy; Center for Translational Research on Autoimmune and Allergic Diseases, University of Piemonte Orientale, Novara, Italy; Division of Thoracic Surgery, Humanitas Cancer Center, IRCCS Humanitas Research Hospital, Rozzano, Milan, Italy

## Abstract

Despite well-documented metabolic and hematopoietic alterations during tumor development^1^, the mechanisms underlying this crucial immunometabolic intersection have remained elusive. Of particular interest is the ligand-activated transcription factor retinoic acid-related orphan receptor 1 (RORC1/RORγ), whose activity is boosted by cholesterol metabolites^2^, acting as a modulator of cancer-related emergency myelopoiesis^3^, while hypercholesterolemia itself is associated with dysregulated myelopoiesis^4,5^. Here we show that both cancer growth and hypercholesterolemic diet can independently or cooperatively activate RORγ-dependent expansion of myeloid-derived suppressor cells (MDSCs) and M2 polarization of tumor-associated macrophages (TAMs), thereby supporting cancer spread. Moreover, we report that tumor development enhances the hepatic production of IL-1β and IL-6, which in turn promote upregulation of hepatic proprotein convertase subtilisin/kexin type 9 (PCSK9) gene, as we confirmed in models of fibrosarcoma, melanoma, colorectal (CRC), and lung cancer, as well as in CRC, non-small-cell lung cancer (NSCLC), breast (BRC), pancreatic ductal adenocarcinoma (PDAC), biliary tract carcinoma (BTC) and pancreatic neuroendocrine tumor (PNET) patients. Importantly, lowering cholesterol levels prevents MDSC expansion and M2 TAM accumulation in a RORγ-dependent manner, unleashing specific anti-tumor immunity and improving the efficacy of anti-PD-1 immunotherapy. Overall, we identify RORγ as a novel sensor of lipid disorders in cancer bearers, bridging hypercholesterolemia and pro-tumor myelopoiesis.

---

Cancers generate complex immunological stressors that can alter the myelopoietic output, leading to the formation of heterogeneous myeloid cell populations (*i.e.*, TAMs and MDSCs) endowed with immunosuppressive and tumor-promoting activities^6-9^. The concerted action of selected transcription factors, such as c/EBPβ, RORγ, STAT3, IRF8, and p50 NF-κB^3,10,11^, translates tumor-derived signals into altered myelopoiesis, causing changes in the differentiation, metabolism, and activation state of myeloid cells^1,12^. Adipose tissue signaling has also been shown to influence the composition of myeloid cells in various tissues^13-15^ and contribute to cancer-related immune dysfunctions^16^. While the correlation between cholesterol levels and cancer risk or progression remains controversial^17^, compelling evidence links dysregulated lipid and cholesterol pathways to cancer invasion and metastasis through transcriptional and epigenetic changes that reprogram tumor metabolism, favoring inflammation in the tumor microenvironment (TME) while evading immune destruction^16,18-25^. Indeed, recent studies have brought attention to the association between cholesterol dysmetabolism and tumor progression^26^. Furthermore, hepatic proprotein convertase subtilisin/kexin type 9 (PCSK9), which interferes with cholesterol absorption and clearance by promoting endo-lysosomal degradation of hepatic LDLR^27^, has been found to reduce the recycling of MHC I on the surface of tumor cells regardless of its cholesterol-regulating functions, potentially affecting immune checkpoint therapy outcomes^28^. A further correlation between hypercholesterolemia and myeloid cell expansion, mobilization, and inflammatory activities comes from cardiovascular studies^4,5^, suggesting that the identification of molecular sensors of lipid dysregulation may also be relevant for diseases associated with chronic metabolic inflammation, such as atherosclerosis. Given the critical role of retinoic acid-related orphan receptor (RORC1/RORγ) in controlling emergency myelopoiesis in cancer patients^3^ and its activation by cholesterol and its metabolites^2^, here we sought to determine the potential involvement of this cholesterol sensor in linking lipid dysmetabolism to pro-tumoral myelopoiesis.

## Cancers alter cholesterol metabolism

We initially conducted a comprehensive evaluation of blood cholesterol levels in mice bearing various tumor types (*i.e.*, MN/MCA1 fibrosarcoma, B16-F10 melanoma, MC38 colon adenocarcinoma, and LLC Lewis lung carcinoma) at both early disease (ED) and advanced disease (AD) stages^3^ (Supplementary Fig. 1a). Our findings revealed a consistent increase in total and LDL cholesterol levels, particularly in AD mice (Fig. 1a and Extended Data Fig. 1a, b). We also observed alterations in total (Fig. 1b), LDL, and HDL (Extended Data Fig. 1c) cholesterol levels in stage I-II (early) and stage III/IV (advanced) cancer patients, including non-small-cell lung cancer (NSCLC, n=146), colorectal (CRC, n=65), breast cancer (BRC, n=36), pancreatic ductal adenocarcinoma (PDAC, n=58), biliary tract cancer (BTC, n=35), and pancreatic neuroendocrine tumor (PNET, n=36). We then evaluated the expression of cholesterol regulatory enzymes in the liver and small intestine, as these organs are the main regulators of cholesterol uptake, synthesis, and excretion^29^. mRNA expression of *Hmgcr,* a rate-limiting enzyme in cholesterol biosynthesis, and *Ldlr*, the receptor of low-density lipoprotein^27^, showed no consistent changes in livers from MN/MCA1, B16- F10, LLC, and MC38 mice (Extended Data Fig. 1d; Supplementary Table 1). In contrast, the expression levels of genes encoding key regulators of blood cholesterol clearance (*i.e.*, *Vldlr*, *Apoe*, *Apob*)^27^ were downmodulated in tumor-bearing (TB) mice. Likewise, the expression of several key enzymes mediating the early steps of oxysterol and bile acid formation (*i.e.*, *Cyp7a1*, *Cyp27a1*, *Cyp7b1*)^30^ and their efflux and biliary excretion (*i.e.*, *Abcg5*, *Abcg8*, *Abcb11*)^27,31^ (Extended Data Fig. 1d; Supplementary Table 1) was decreased. In the small intestine, we observed decreased expression of cholesterol efflux transporter genes *Abcg5*/*Abcg8*, while the expression of *Npcll1*, involved in the absorption of cholesterol, remained unchanged (Extended Data Fig. 1e). Consistently, MN/MCA1 mice showed reduced total bile acid (TBA) levels in their feces (Extended Data Fig. 1f). In agreement with the increased LDL cholesterol levels, hepatic PCSK9 mRNA (Extended Data Fig. 1d) and protein levels (Extended Data Fig. 1g) were upregulated in livers from AD TB mice, which correlated with downregulation of hepatic LDLR protein (Extended Data Fig. 1g). Modulation of hepatic cholesterol metabolism gene expression was also confirmed in healthy liver parenchyma from CRC patients (n=23) compared to distant healthy liver parenchyma of benign adenomas or angiomas (CTRL, n=6) from non-dyslipidemic patients (Extended Data Fig. 1h). Moreover, treatment of primary hepatocytes with MN/MCA1 tumor-conditioned medium (TCM) increased *Pcsk9* gene expression, while it reduced both *Vldlr* and *Abcg8* gene expression (Extended Data Fig. 1i; Supplementary Table 1).

**Figure 1.**
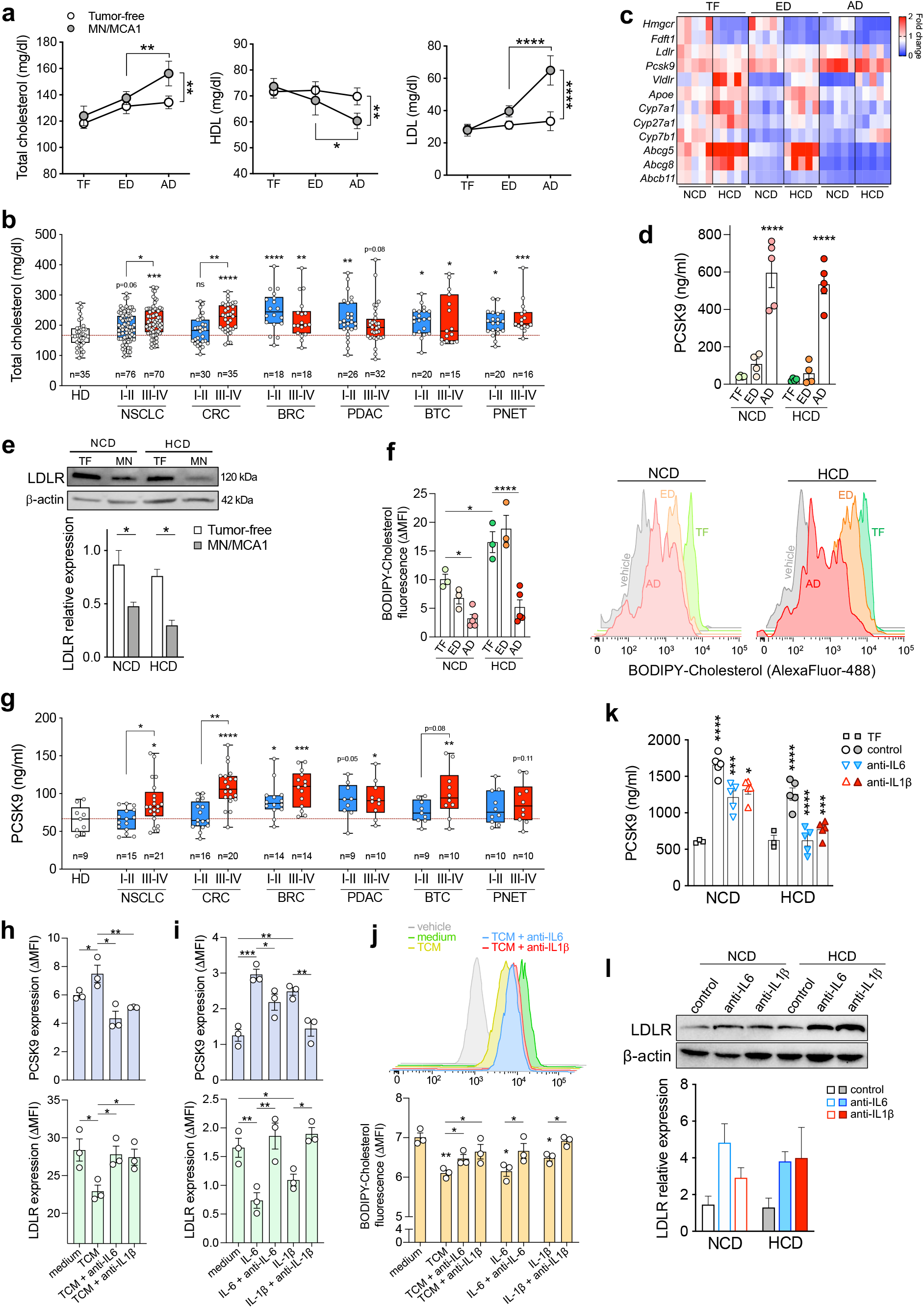
Tumor progression alters cholesterol metabolism. **a,** Total HDL and LDL blood cholesterol levels in MN/MCA1 mice, at early (ED) or advanced disease (AD) stage compared to healthy, age- and sex-matched tumor-free (TF) mice (n=4). **b,** Total blood cholesterol levels in patients with early (I-II) *vs.* advanced (III-IV) NSCLC (n=76 *vs.* n=70), CRC (n=30 *vs.* n=35), BRC (n=18 *vs.* n=18), PDAC (n=26 *vs.* n=32), BTC (n=20 *vs.* n=15), or PNET (n=20 *vs.* n=16) compared to healthy donors (HD, n=35). **c,** Heatmap showing differential mRNA expression of genes involved in hepatic cholesterol metabolism (n=5) and (**d**) circulating PCSK9 levels (n=5) in ED or AD MN/MCA1 *vs.* TF mice fed with either normal chow diet (NCD) or high-cholesterol diet (HCD). **e**, Immunoblot analysis of LDLR protein in the livers from NCD- or HCD-fed MN/MCA1 mice at ED or AD *vs.*TF similarly fed (n=2). **f**, FACS quantification of BODIPY-Cholesterol mean fluorescence intensity (ΔMFI) of hepatic CD45^-^CD31^-^ cells from MN/MCA1 mice at ED (n=3) or AD (n=5) on NCD or HCD compared to TF mice (n=3) similarly fed. **g**, Circulating PCSK9 levels in patients with early (I-II) *vs.* advanced (III-IV) NSCLC (n=15 *vs.* n=21), CRC (n=16 *vs.* n=20), BRC (n=14 *vs.* n=14), PDAC (n=9 *vs.* n=10), BTC (n=9 *vs.* n=10), or PNET (n=10 *vs.* n=10) in comparison with healthy donors (HD, n=9). **h-j,** FACS analysis of PCSK9 and LDLR protein expression (**h, i**) or BODIPY-Cholesterol (**j**) in Hepa1-6 cells treated with MN/MCA1 tumor-conditioned medium (TCM), IL-6, or IL-1β alone or in combination with anti-IL6 or anti-IL1β agent (n=3). **k,** Circulating PCSK9 levels in TF (n=3) or MN/MCA1 bearing mice in AD (n=5) on NCD or HCD in the absence (control) or presence of anti-IL-6 or anti-IL-1β agent. **l**, Immunoblot (upper panel) and relative densitometry quantification (lower panel, n=2) of hepatic LDLR protein using protein extracts from mice treated as described above. Data are representative of four (**a**), two (**e, h, l**), or three (**i**) independent experiments; **c, d, f, j, k,** one experiment was performed. **a, d-f, h-l,** Data are presented as mean ± SEM; **b, g,** Box-and-whisker min-to-max plots. \**P* < 0.05, \*\**P* < 0.01, \*\*\**P* < 0.001, \*\*\*\**P* < 0.0001 between selected relevant comparisons. **a,** Two-way ANOVA with Tukey’s multiple comparisons test. **b-l,** One-way ANOVA with Tukey’s multiple comparisons test.

We next assessed the effects of diet on plasma cholesterol levels by feeding mice bearing transplantable fibrosarcoma MN/MCA1 or conditional *K-ras^LSL-G12D/+^;p53*^fl/fl^ (KP) mice developing NSCLC^32^ on either normal chow diet (NCD) or hypercholesterolemic diet (HCD) (Extended Data Fig. 2a, d). We observed that cancer development further enhanced HCD-induced hypercholesterolemia (Extended Data Fig. 2b, c, e). Furthermore, while *Vldlr*, *Cyp7a1*, *Cyp27a1*, *Abcg5*, *Abcg8* gene expression was upregulated in HCD-fed tumor-free mice, MN/MCA1 tumor development prevented this upregulation, as well as that of the *Apoe* and *Abcb11* genes (Fig. 1c; Supplementary Table 1). Of note, AD marked the most pronounced changes in gene expression, including *Pcsk9* whose mRNA expression (Fig. 1c; Supplementary Table 1), and blood protein levels (Fig. 1d) increased regardless of diet type, correlating with reduced levels of LDLR protein expression (Fig. 1e). Fittingly, tumor development in KP mice increased the circulating levels of PCSK9 (Extended Data Fig. 2f). We next evaluated whether the increase in blood cholesterol was also associated with changes in the hepatic levels of mevalonate and squalene, crucial intermediates of the cholesterol biosynthesis chain^26^. MN/MCA1 progression resulted in reduced levels of both metabolites (Extended Data Fig. 2g), suggesting that hepatic cholesterol biosynthesis does not significantly contribute to the observed increase in blood cholesterol in cancer bearers. Furthermore, regardless of the type of diet, oral administration of fluorescence-labeled cholesterol (BODIPY-Ch) clearly showed a decrease in hepatic cholesterol during tumor progression (Fig. 1f), indicating its accumulation in extrahepatic tissues, such as tumor cells (CD45^-^CD31^-^), TAMs, myeloid progenitors (both CMPs and GMPs), and circulating Cd11b^+^Ly6C^hi^ monocytic cells (Extended Data Fig. 2h, i). This was further supported by reduced excretion of total bile acids in both diet regimens (Extended Data Fig. 2j). Finally, serum from TB mice increased PCSK9 expression in primary hepatocytes (Extended Data Fig. 2k), indicating that tumors secrete factors that modulate hepatic cholesterol metabolism, particularly upregulating PCSK9. In line with these findings, circulating PCSK9 was significantly elevated in patients with advanced cancers (Fig. 1g).

**Figure 2.**
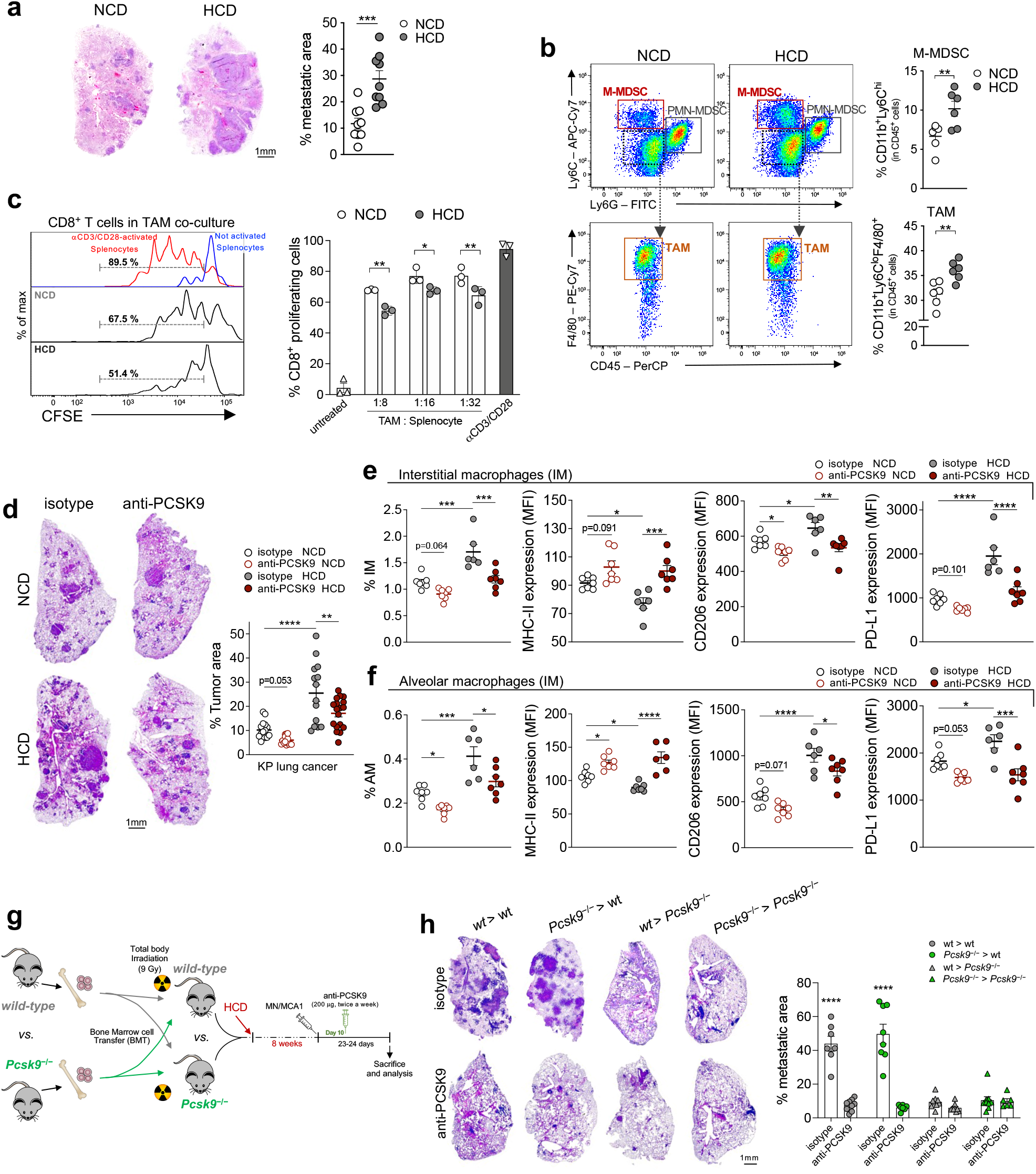
Cholesterol levels modulate cancer-driven myeloid immunosuppression. **a,** Lung metastatic area in NCD- or HCD-fed MN/MCA1 mice (n=3). Representative images are shown. Scale bar, 1 mm. **b,** Representative FACS plot (left) and frequencies (right) of intratumoral CD11b^+^Ly6G^-^ Ly6C^hi^ monocytic (M)-MDSCs and CD11b^+^Ly6G^-^Ly6C^lo^F4/80^+^ TAMs from NCD- or HCD-fed MN/MCA1 mice (n=6). **c,** Effects of TAMs, FACS-sorted from NCD- or HCD-fed MN/MCA1 mice, on the proliferation of CFSE-labeled CD8^+^ T cells activated with anti-CD3/anti-CD28 (n=3). **d-f,** NCD- or HCD-fed mice bearing the genetically induced *Kras*^LSL-G12D/+^; *p53*^fl/fl^ (KP) lung cancer and treated with anti-PCSK9 or isotype control (n=5). **d,** Lung metastatic area. Representative images are shown. Scale bar, 1 mm. FACS quantification of interstitial (IM) (**e**) and alveolar macrophages (AM) (**f**) and their relative mean fluorescence intensity (MFI) of MHC-II, CD206, and PD-L1. **g, h,** Experimental design (**g**): wild-type (wt) or PCSK9-deficient (*Pcsk9*^-/-^) mice were lethally irradiated (9 gy) and respectively transplanted with either wt (wt>wt; wt>*Pcsk9*^-/-^) or *Pcsk9*^-/-^ (*Pcsk9*^-/-^>wt; *Pcsk9*^-/-^>*Pcsk9*^-/-^) bone marrow (BM) cells. After 4 weeks, needed for complete hematopoietic reconstitution, mice were conditioned for an additional 8 weeks with HCD and then engrafted with MN/MCA1 cells. After 25 days of tumor growth, mice were sacrificed for sample collection and analysis, including the evaluation of lung metastatic areas (n=5) (**h**); representative images are shown. Scale bar, 1 mm. Data are representative of at least five (**a, b**) or two (**c-f**) independent experiments; **h,** one experiment was performed. Data are presented as mean ± SEM. \**P* < 0.05, \*\**P* < 0.01, \*\*\**P* < 0.001, \*\*\*\**P* < 0.0001 for selected relevant comparisons. **a, b,** Unpaired two-tailed *t*-test. **c**, One-way ANOVA with Šidák’s multiple comparisons test. **d-f,** One-way ANOVA with Tukey’s multiple comparisons test. **h,** Two-way ANOVA with Tukey’s multiple comparisons test.

Emerging evidence has linked inflammatory conditions (*e.g.,* infections, rheumatoid arthritis, and acute coronary syndrome) and their mediators to altered lipid metabolism^33-35^ and PCSK9 upregulation^36^. In particular, IL-6 and IL-1β, which play critical roles in both cancer development^37^ and metabolic alterations^14^, have been found to influence the activity of PCSK9/LDLR axis as well as cholesterol levels^38-40^. As shown in Extended Data Fig. 3a, TCM contained significant levels of IL-6 and IL-1β. Fittingly, stimulation of hepatocytic cells (Hepa1-6) with TCM, IL-6 or IL-1β enhanced PCSK9 expression and decreased LDLR protein expression, while treatment with an anti-IL-6 or anti-IL-1β agent reversed this effect (Fig. 1h, i), restoring the ability of the hepatocytes to uptake extracellular cholesterol (Fig. 1j). In line with these findings, both liver (Extended Data Fig. 3b-c, e-f) and adipose tissue (Extended Data Fig. 3d) from TB mice and the healthy liver parenchyma from CRC patients (Extended Data Fig. 3g) showed enriched levels of both IL-6 and IL-1β. Of note, no changes were observed for *Pcsk9* expression in adipose tissue, while *Ldlr*, *Vldlr* and *Acat2* gene expression was decreased, the latter responsible for the esterification of cholesterol (not shown). Additionally, the administration of anti-IL6 or anti-IL1β agent to MN/MCA1 mice led to the suppression of PCSK9 upregulation (Fig. 1k and Extended Data Fig. 3h) and restoration of LDLR protein levels (Fig. 1l). Interestingly, these effects were consistent with the inhibition of tumor growth and reduction in blood cholesterol levels observed in NCD- or HCD-fed mice (Extended Data Fig. 3i, j).

**Figure 3.**
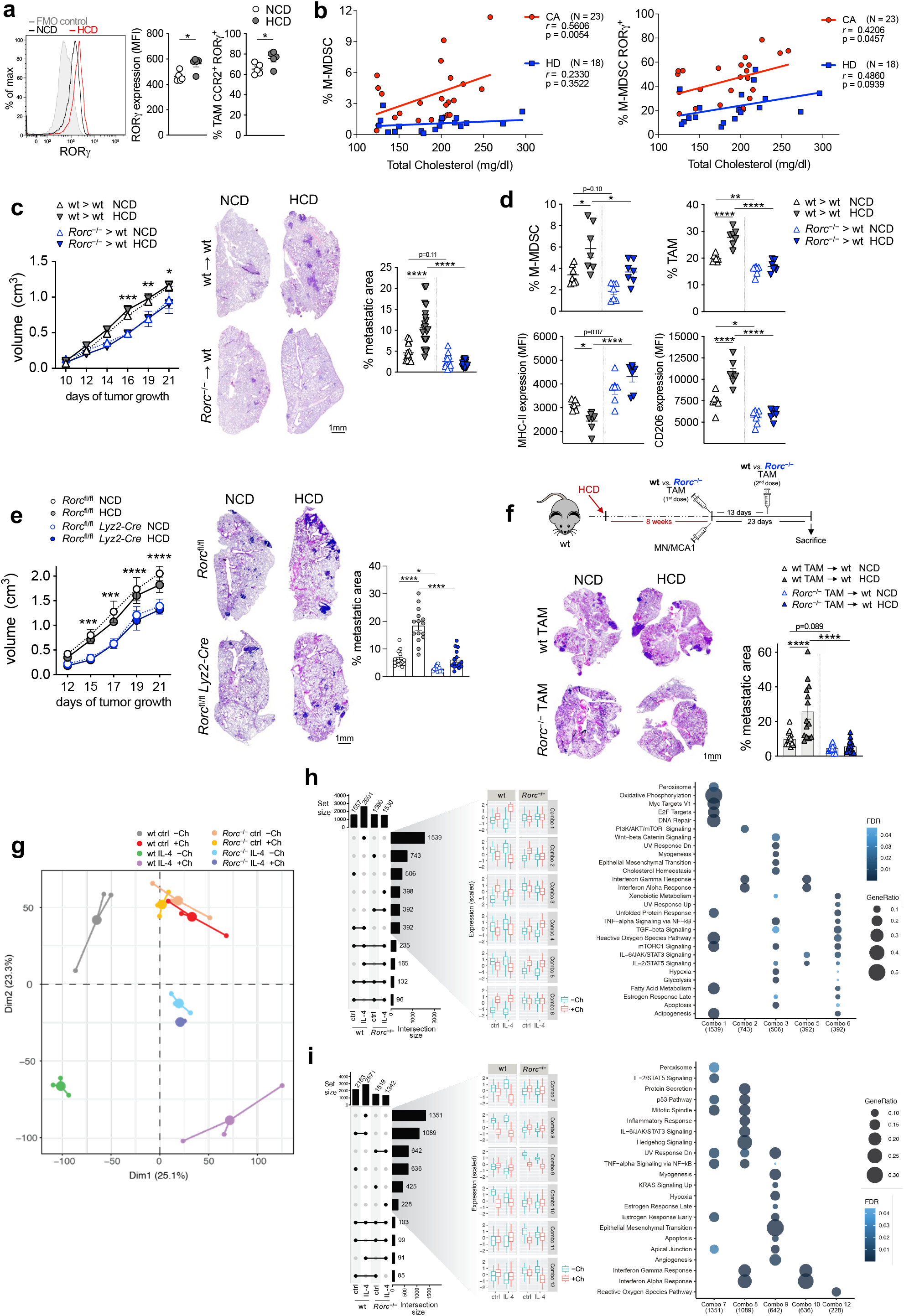
RORγ bridges hypercholesterolemia and protumoral myelopoiesis. **a,** RORγ expression and RORγ^+^ cells in CCR2^+^ TAMs from NCD- or HCD-fed MN/MCA1-bearing mice (n=5). **b,** Linear Pearson correlation between total cholesterol levels and M-MDSC (HLA-DR^lo/–^CD14^+^) (left) or M- MDSC RORγ^+^ cells (right) in PBMCs from healthy donors (HD) or NSCLC patients (CA) (stage III- IV). Pearson correlation coefficient (*r*) and corresponding *p*-value (p) are indicated. **c, d,** NCD- or HCD-fed MN/MCA1 wt mice transplanted with either wt (wt>wt) or *Rorc*^-/-^ (*Rorc*^-/-^>wt) BM cells (n=7). **c**, Primary tumor growth (left) and lung metastatic areas (right). Representative images are shown. Scale bar, 1 mm. **d,** Intratumoral M-MDSC and TAM frequency (upper panels) and MHC-II and CD206 expression in TAMs (lower panels). **e,** Primary tumor growth (left) and lung metastatic area (right) in *Rorc*^fl/fl^ and *Rorc*^fl/fl^ *Lyz2-Cre* MN/MCA1 mice on NCD or HCD (n=5). Representative images are shown. Scale bar, 1 mm. **f** (top), Experimental design: wt mice on NCD or HCD were co-injected with MN/MCA1 and FACS-sorted wt or *Rorc*^-/-^ TAMs (dose 1). A second adoptive transfer of FACS-sorted TAMs were intratumorally administered after 13 days of tumor growth (dose 2) (n=5). **f** (bottom), Lung metastatic areas. Representative images are shown. Scale bar, 1 mm. **g-i,** RNA-seq analysis on wt or *Rorc*^-/-^ BM-derived macrophages (BMDMs) differentiated with M-CSF, with or without cholesterol (Ch) supplementation, and then treated or not (ctrl) with IL-4 (n=3). **g,** Principal components analysis (PCA) based on transcriptional profiles. **h, i,** (left panels), Upset plots of significant (FDR < 0.05 & |log2FC| > 0.5) up- (**h**) or down-regulated (**i**) genes found in each group supplemented with Ch *vs.* non-Ch-supplemented control across the four different biological conditions: wt untreated ctrl; wt IL-4; *Rorc*^-/-^ untreated ctrl; and *Rorc*^-/-^ IL-4. An intersection between two or more gene sets is displayed as black dots connected by a solid black line. Boxplots of the scaled expression distribution for the first six combination gene sets obtained from the up- (**h**) or down-regulated (**i**) genes are depicted. **h, i**, (right panels) Over-representation analysis of MSigDB Hallmark 2020 terms for the first six combination sets obtained from the up- (Combo 1-6) (**h**) or down-regulated (Combo 7-12) (**i**) genes. Combo 4 and 11 did not obtain significant (FDR < 0.05) terms. **a,** Data are representative of five independent experiments; **b-i,** one experiment was performed. **a, c-f,** Data are presented as mean ± SEM. \**P* < 0.05, \*\**P* < 0.01, \*\*\**P* < 0.001, \*\*\*\**P* < 0.0001 between selected relevant comparisons. **a,** Unpaired two-tailed *t*-test. **b,** Pearson’s correlation with two-tailed *P* value. **c** (left), **e** (left), two-way ANOVA with Tukey’s multiple comparisons test. **c** (right)**, d, e** (right)**, f**, One-way ANOVA with Tukey’s multiple comparisons test. **g-i,** Detailed description of statistical analysis is described in Methods section.

## Hypercholesterolemia boosts protumoral myelopoiesis

Interestingly, while HCD did not affect primary MN/MCA1 tumor growth (Extended Data Fig. 4a), it did lead to an increased number of circulating (mCherry^+^) tumor cells (CTCs) (Extended Data Fig. 4b) and a higher frequency of CD31^+^ endothelial cells in both primary and metastatic lungs (Extended Data Fig. 4c), ultimately resulting in a pronounced exacerbation of lung metastasis formation (Fig. 2a). Similarly, hypercholesterolemia also promoted the formation of lung metastasis in the K1735- M2 melanoma model (Extended Data Fig. 4d) and contributed to the growth of tumor lesions in the genetic KP lung cancer model (Extended Data Fig. 4e).

**Figure 4.**
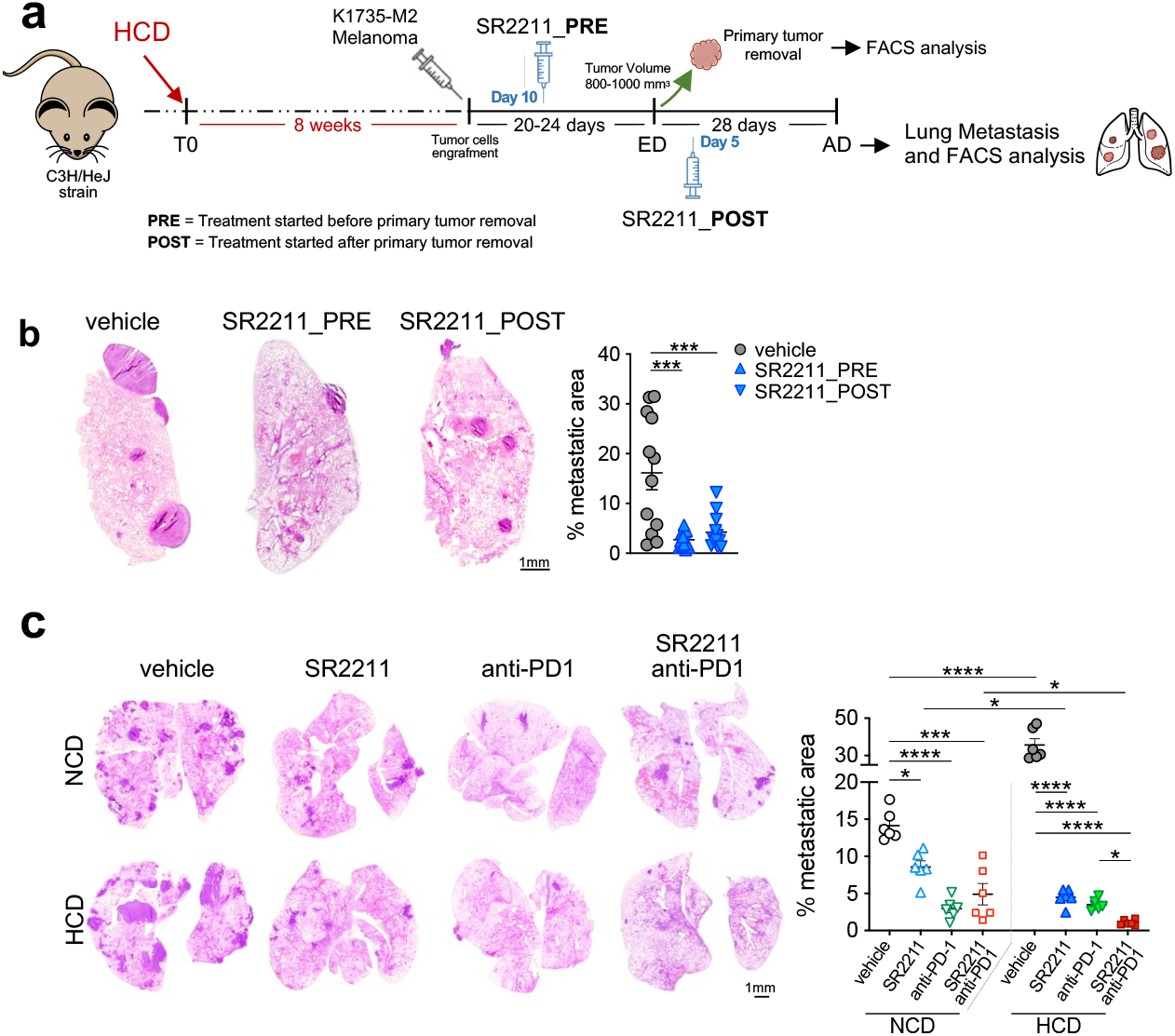
RORγ inhibition improves the response to immunotherapy. **a,** Experimental design: adult wt C3H/HeJ mice were fed with HCD for 8 weeks before being injected with K1735-M2 tumor cells. When the tumor volume reached ∼0.80-1.00 cm^3^, typically 20-24 days after tumor cell injection, primary tumors were surgically removed. Twenty-eight days after tumor removal, mice were sacrificed for sample collection and analysis. SR2211 was administered to a first group of mice 10 days before tumor removal (SR2211_PRE). A second group of mice received SR2211 treatment 5 days after tumor removal (SR2211_POST). **b,** Lung metastatic areas (n=5) in HCD-fed K1735-M2 mice treated with either SR2211 or vehicle control. Representative images are shown. Scale bar, 1mm. **c,** Lung metastatic areas in NCD- or HCD-fed MN/MCA1 wt mice treated with vehicle, SR2211, anti-PD1, or SR2211 plus anti-PD1 (n=5). Representative images are shown. Scale bar, 1mm. One experiment was performed. Data are expressed as mean ± SEM. \**P* < 0.05, \*\**P* < 0.01, \*\*\**P* < 0.001, \*\*\*\**P* < 0.0001. One-way ANOVA with Tukey’s multiple comparisons test.

Since obesity^12,14,41^ and high cholesterol^4,5^ are associated with preferential differentiation of hematopoietic stem cells (HSCs) into the monocyte-macrophage cell lineage, and given that myeloid cells exhibit tumor promoting activities^7-9^, we further investigated the myeloid compartment of hypercholesterolemic MN/MCA1 mice. HCD increased the frequency of blood and splenic CD11b^+^Ly6C^hi^ monocytic cells, under both tumor-free (TF) and TB conditions (Extended Data Fig. 4f), whereas, in contrast with previous observations^15^, the frequency of Cd11b^+^Ly6G^+^ granulocytic cells remained unaffected (Extended Data Fig. 4f). Flow cytometry analysis of tumor-infiltrating myeloid cells (Supplementary Fig. 1b) confirmed that HCD specifically increased the presence of TAMs and M-MDSCs (Fig. 2b), which showed higher lipid-load levels as evidenced by LipidTOX neutral lipid (*i.e.*, cholesterol, cholesterol esters, triglycerides) staining (Extended Data Fig. 4g) and by BODIPY-Ch uptake (Extended Data Fig. 2h). In-depth lipidomic analysis of TAMs revealed significant imbalances in numerous lipid species induced by HCD (Extended Data Fig. 4h-j). Remarkably, the lipidomic analysis revealed 52 significantly enriched lipids and 148 down-regulated ones (Extended Data Fig. 4i). Importantly, this lipidomic profiling showed an overall enrichment of triglyceride (TG) and sterol (ST) lipid classes, with a notable increase in cholesterol (CH) levels at the expense of its esterified forms (CE) (Extended Data Fig. 4k). Noteworthy, the lipid changes observed in TAMs from HCD-fed mice were accompanied by a prominent shift toward M2 polarization^42^, as judged by the downregulation of M1 markers (*i.e.*, TNFα, MHC-II, iNOS, and CCR7) (Extended Data Fig. 4l, upper panels) and simultaneous increase in M2 indicators (*i.e.*, CD206, PD-L1, CD204, and IDO1) (Extended Data Fig. 4l, lower panels).

As macrophages and MDSCs are known to contribute to the formation of pre-metastatic niches^8,9,43^, and there is evidence that cholesterol metabolites and, more in general, obesity may promote cancer metastasis^15,26,44-46^, we analyzed the frequency of lung myeloid cells in mice fed NCD or HCD (Extended Data Fig. 5a). HCD-fed TB mice showed an increased frequency of CD11b^+^Ly6C^hi^ monocytic cells (Extended Data Fig. 5b), as well as of BM-derived interstitial macrophages (IM) and resident alveolar macrophages (AM) (Extended Data Fig. 5c and Supplementary Fig. 1c)^47,48^. In contrast, conventional CD103^+^ (cDC1) and CD11b^+^ (cDC2) dendritic cell subsets (Supplementary Fig. 1c)^47^ remained unaffected (Extended Data Fig. 5d). Furthermore, HCD induced a shift toward M2 polarization (TNFα^lo^MHC-II^lo^CD206^hi^PD-L1^hi^) of IM and AM cells (Extended Data Fig. 5e). These HCD-induced alterations in myeloid cell frequency and phenotype were similarly observed in the K1735-M2 metastatic melanoma model (Extended Data Fig. 5f-i). Additionally, while HCD improved the frequency of CD4^+^ and CD8^+^ T cells in both primary tumors and metastatic lungs, their phenotype displayed signs of dysfunction and exhaustion^49^, with higher expression of program cell death protein 1 (PD-1) and cytotoxic T-lymphocyte antigen 4 (CTLA4) and lower expression of interferon-γ (IFNγ) (Extended Data Fig. 5j, k). We further assessed the proliferation rate of T cells co-cultured with TAMs (Fig. 2c) or M-MDSCs (Extended Data Fig. 5l), FACS-sorted from HCD or NCD-fed mice, and found that myeloid cells from HCD-fed mice significantly reduced CD8^+^ T-cell proliferation compared to their NCD-fed counterparts, indicating that hypercholesterolemia induces the expansion of myeloid cells endowed with suppressive activity. Given the observed hepatic elevation of PCSK9 promoted by tumor growth (Fig. 1c, d, g and Extended Data Fig. 2f) and its association with immunosuppressive myeloid cells (Fig. 2b and Extended Data Fig. 4f, 5a-e), we explored the potential impact of administration of the cholesterol-lowering anti-PCSK9 monoclonal antibody, mAb1^50^, on protumoral myelopoiesis and tumor development in both MN/MCA1 (Extended Data Fig. 6a) and KP cancer models (Extended Data Fig. 6j). Strikingly, anti-PCSK9 antibody treatment lowered blood cholesterol levels in MN/MCA1 (Extended Data Fig. 6b) and KP mice (Extended Data Fig. 6k), and decreased primary MN/MCA1 (Extended Data Fig. 6c) and KP tumors (Fig. 2d), as well as MN/MCA1 lung metastasis (Extended Data Fig. 6d), with a prominent effect in the context of HCD. These beneficial effects were accompanied by a reduced frequency of M-MDSCs, TAMs, and lung metastasis-associated IMs and AMs in MN/MCA1 (Extended Data Fig. 6e-g), as well as decreased M-MDSCs (Extended Data Fig. 6l), IMs and, AMs in primary KP lung tumors (Fig. 2e, f). Consistent with the reduction in circulating cholesterol levels (Extended Data Fig. 6b), anti-PCSK9 treatment in MN/MCA1-bearing mice decreased neutral lipid content in TAMs (Extended Data Fig. 6f). This restoration of lipid levels corresponded with a shift in TAMs toward an M1 phenotype (MHC-II^hi^CD206^lo^) (Extended Data Fig. 6f), as well as in IMs and AMs from the metastatic lungs (Extended Data Fig. 6g). Likewise, anti-PCSK9 blockade in KP lung cancer bearers reduced blood cholesterol (Extended Data Fig. 6k), whereas it increased expression of the M1 markers TNFα (Extended Data Fig. 6m) and MHC-II (Fig. 2e, f) and decreased M2 markers (CD206, PD-L1) in IMs and AMs (Fig. 2e, f). These anti-PCSK9- mediated events were associated with reduced expression of the exhaustion markers PD-1 and CTLA4, accompanied by increased production of IFNγ by CD8^+^ T cells in both primary MN/MCA1 tumors (Extended Data Fig. 6h) and metastatic lungs (Extended Data Fig. 6i), as well as in the KP lung cancer model (Extended Data Fig. 6n).

Given that anti-PCSK9 treatment had no effect on tumor *cell* viability and proliferation *in vitro* (Extended Data Fig. 6o), and to rule out possible tumor- or immune cell-intrinsic roles of PCSK9, we transplanted wild-type (wt) or *Pcsk9*^-/-^ BM cells into lethally irradiated wt or *Pcsk9*^-/-^ recipient mice, and vice versa, and evaluated the development of MN/MCA1 under HCD conditions (Fig. 2g). Extramedullary deficiency of PCSK9 markedly lowered circulating cholesterol levels (Extended Data Fig. 6p) and inhibited tumor growth (Fig. 2h and Extended Data Fig. 6q), along with a decrease in the frequency and M2 polarization of TAMs (Extended Data Fig. 6r). As expected, PCSK9 deletion in the hematopoietic compartment did not produce these effects, while anti-PCSK9 treatment elicited cholesterol-lowering and anti-tumoral activities only in extramedullary PCSK9-proficient recipient mice (Fig. 2h and Extended Data Fig. 6q). Altogether, these results indicate hepatic PCSK9 as the main driver of cholesterol metabolism alterations in cancer bearers.

We then proceeded to characterize the roles of the two main pathways of macrophage accumulation, CSF-1/CSF-1R^51^ and CCL2/CCR2^8^, in hypercholesterolemic MN/MCA1 mice. Even though HCD did not affect CSF-1R (CD115) expression in TAMs, IMs, and AMs (Extended Data Fig. 7a), in agreement with previous reports^52,53^ it did increase the frequency of CCR2^+^ myeloid cells by augmenting their CCR2 surface expression levels (Extended Data Fig. 7b). Furthermore, we detected higher CSF-1 (M-CSF) and CCL2 mRNA expression levels in the TME of HCD-fed mice (Extended Data Fig. 7c). Coherently, treatment with anti-CSF1R or anti-CCR2 agent reduced primary MN/MCA1 growth and lung metastatic burden (Extended Data Fig. 7d, e), paralleled by a marked reduction in M-MDSCs (Extended Data Fig. 7f), as well as TAMs (Extended Data Fig. 7g), IMs, and AMs (Extended Data Fig. 7h), which, however, displayed higher levels of TNFα (Extended Data Fig. 7g, h). Fittingly, anti-CSF1R or anti-CCR2 treatment increased the frequency of CD8^+^ T cells and their IFNγ expression levels (Extended Data Fig. 7i). Given the significance of the CSF-1/CSF-1R and CCL2/CCR2 axes in the differentiation and mobilization of myeloid progenitors^1^, we next evaluated the lineage commitment of Lin^−^c-Kit^+^Sca-1^−^ (LK) bone marrow (BM) hematopoietic progenitors in HCD-fed TB mice. HCD increased the lipid and cholesterol load of myeloid progenitors (Extended Data Fig. 2i and 8b), leading to enhanced commitment of common myeloid progenitors (CMPs) to granulocyte-macrophage progenitors (GMPs) in both MN/MCA1 (Extended Data Fig. 8a, b) and KP mice (Extended Data Fig. 8c). Both anti-PCSK9 treatment (Extended Data Fig. 8b, c) and extramedullary PCSK9 deletion (Extended Data Fig. 8d) prevented these effects, as result of a drastic reduction of systemic cholesterol levels (Extended Data Fig. 6b, p). Moreover, anti-CSF1R and anti-CCR2 treatments also hampered the HCD-induced commitment of GMPs (Extended Data Fig. 8e).

Taken together, these results suggest that PCSK9 plays a pivotal role in the cholesterol-induced modulation of myeloid progenitor differentiation, ultimately affecting tumor-associated macrophages and myeloid cell populations within the TME.

## RORγ tunes myelopoiesis to cholesterol metabolism

As we have previously reported RORC1/RORγ to be a marker of advanced cancer inflammation and a promoter of both MDSC and TAM differentiation^3^, and given that cholesterol precursors and oxysterol derivatives not just enhance RORγ intrinsic activity^2^ but are also aberrantly expressed in both cancer^54^ and obesity^55,56^, we asked whether increased cholesterol levels would influence cancer myelopoiesis in a RORγ-dependent manner. Analysis of tumors, lungs, BM, and spleens from HCD-fed MN/MCA1 mice showed increased RORγ expression in BM-derived CCR2^+^ subsets of TAMs (Fig. 3a), M-MDSCs, IMs and AMs (Extended Data Fig. 9a-d) but not in their CCR2^-^ counterparts (Extended Data Fig. 9e). Congruently, HCD led to the upregulation of RORγ in both CMPs and GMPs (Extended Data Fig. 9f).

To validate our findings in cancer patients, we assessed monocytic cell subsets in PBMCs (Supplementary Fig. 1d) from healthy donors (HDs, n=18) and NSCLC patients (CA, stage III/IV, n=23). In line with our observation in the mouse model, the frequency of M-MDSCs and their RORγ^+^ subset correlated with total or LDL cholesterols in lung cancer patients (Fig. 3b and Supplementary Table 2). To test whether circulating cholesterol would modulate RORγ transcriptional activity, we co-transfected HEK293 cells with a vector constitutively expressing the *Rorc* gene and a plasmid expressing the luciferase reporter gene under the control of the minimal promoter of the *il17a* gene (minIL17Aprom-Luc)^57^, transcriptional target of RORγ (Extended Data Fig. 9g). Transfected cells were treated with free cholesterol, LDL, the RORγ agonist SR0987, or desmosterol, an immediate cholesterol precursor acting as endogenous RORγ agonist^2^. All four conditions significantly induced luciferase activity (Extended Data Fig. 9h), indicating that circulating cholesterol is crucial for the activation of RORγ-dependent cellular programs. Next, we treated transfected HEK293 cells with serum from NCD- or HCD-fed TB mice, or with serum from TF mice, and measured luciferase activity. While exposure to serum from NCD-fed TB mice (TB-NCD) led to increased RORγ transcriptional activity, this response was consistently higher when cells were stimulated with serum from HCD-fed TB mice (TB-HCD) (Extended Data Fig. 9i). Notably, TCM alone poorly affected RORγ activity and its increase by cholesterol supplementation was inhibited by RORγ inhibitor SR2211^3,58^ (Extended Data Fig. 9j).

Considering that RORγ transcriptional activation by cholesterol-rich serum correlated with the expansion of monocytic cells seen in hypercholesterolemic tumor bearers (Fig. 2b and Extended Data Fig. 4f), we aimed to evaluate the extent of the interplay between RORγ and hypercholesterolemia in NCD- or HCD-fed MN/MCA1 RORγ-deficient mice (*Rorc*^-/-^). While RORγ deficiency did not affect cholesterol levels (Extended Data Fig. 10a, b), it reduced primary tumor growth (Extended Data Fig. 10c) and inhibited the formation of metastasis and circulating mCherry^+^ MN/MCA1 tumor cells (Extended Data Fig. 10d, e), regardless of the diet type. Consistent with our previous findings, a predominant M2-polarized immune profile was observed in the TME of HCD- *vs.* NCD-fed mice, as judged by increased mRNA expression of *Il4*, *Tgfb*, *Ccl22*, *Mrc1* (CD206), *Ccl2*, and *Csf1* (M-CSF) (Extended Data Fig. 10f). This M2 polarization was further supported by an increased frequency of TAMs, lung IMs, AMs, and tumor infiltrating M-MDSCs (Extended Data Fig. 10g). Supporting a key role of RORγ in bridging hypercholesterolemia and protumoral myelopoiesis, all the differences were lost in both NCD- or HCD-fed RORγ null mice (Extended Data Fig. 10c-g), concomitant with a predominant expression of M1 cytokines (*i.e.*, *Tnfa, Cxcl9, Cxcl10, Cxcl11*) (Extended Data Fig. 10f). In contrast, no variation was observed in the PMN-MDSC subset (Extended Data Fig. 10g). Regardless of RORγ expression, TAMs (Extended Data Fig. 10h), IMs, and AMs (Extended Data Fig. 10i) from HCD-fed mice exhibited an accumulation of neutral lipids. Cholesterol-specific Filipin-III staining confirmed the cholesterol overload in TAMs from hypercholesterolemic mice (Extended Data Fig. 10j). Consistent with the role of RORγ as a mediator of cholesterol-induced M2 polarization, the M2-like activation state of TAMs, IMs, and AMs induced by hypercholesterolemia was lost in RORγ null mice, as evidenced by increased expression of TNFα and MHC-II and downmodulation of CD206 and PD-L1 (Extended Data Fig. 10k, l), as well as by a shift of TAMs toward an IL-12p40^high^/IL-10^low^ phenotype (Extended Data Fig. 10m).

To rule out the possibility of any anti-tumoral effect resulting from total RORγ deficiency, we transplanted lethally irradiated wt recipient mice with either *Rorc*^-/-^ or wt BM cells. These mice were then subjected to HCD or NCD conditioning before engrafting them with MN/MCA1 cells (Extended Data Fig. 11a). Hematopoietic RORγ deficiency (*Rorc*^-/-^>wt) curtailed HCD-induced tumor progression (Fig. 3c), which was accompanied by a reduced frequency of M-MDSCs, TAMs (Fig. 3d), IMs, and AMs (Extended Data Fig. 11b), all of which displayed a polarization shift toward an M1 phenotype (Fig. 3d and Extended Data Fig. 11b).

In light of cholesterol-induced expression of RORγ in both CMPs and GMPs (Extended Data Fig. 9f), which suggested a direct link between lipid metabolism and RORγ-dependent emergency myelopoiesis^3^, we assessed the frequency of BM myeloid progenitors in hypercholesterolemic tumor-bearing and lethally irradiated wt mice transplanted with *Rorc*^-/-^ BM cells. In agreement with a previous report^3^, we observed a reduction in Lin^-^c-kit^+^Sca-1^+^ (LSK) hematopoietic cells in the BM of TB mice reconstituted with wt BM cells (wt>wt) under HCD conditions. By contrast, RORγ deficiency in hematopoietic cells (*Rorc*^-/-^>wt) increased the frequency of LSK cells in a diet-independent manner (Extended Data Fig. 11c). Similar results were observed in total RORγ-deficient mice (data not shown). In addition, while CMPs and GMPs displayed elevated intracellular neutral lipids and cholesterol content under HCD conditions, regardless of RORγ expression (Extended Data Fig. 11d, e), the increased transition of CMPs to GMPs observed in hypercholesterolemic mice (Extended Data Fig. 8a) was significantly inhibited by RORγ deficiency (Extended Data Fig. 11c), indicating that RORγ acts as a key transcriptional effector in monocyte/macrophage differentiation induced by hypercholesterolemia.

During our experiments, we also noticed that both whole RORγ-deficient mice (*Rorc^-/-^*) (not shown) and *Rorc^-/-^* BM cell-transplanted mice (*Rorc*^-/-^>wt) displayed an increased number of tumor-infiltrating CD4^+^ and CD8^+^ T cells expressing high IFNγ levels compared to their wt counterparts (Extended Data Fig. 11f). To ascertain whether RORγ directly influenced T cell differentiation, we used myeloid-specific RORγ−deficient mice (*Rorc*^fl/fl^ *Lyz2-Cre*). These mice displayed reduced MN/MCA1 primary tumor growth and metastasis formation (Fig. 3e), particularly under HCD conditions. Correspondingly, specific ablation of RORγ in myeloid cells (*Rorc*^fl/fl^ *Lyz2-Cre* mice) reprogrammed TAMs toward an M1 phenotype, as evidenced by increased MHC-II and decreased CD206 and PD-L1 expression levels (Extended Data Fig. 11g). This phenotypic change was associated with enhanced effector functions of infiltrating CD8^+^ T cells, as judged by the higher expression levels of IFNγ and granzyme-B (Extended Data Fig. 11h). Moreover, in contrast with wt TAMs, adoptive transfer of *Rorc^-/-^* TAMs into either NCD- or HCD-fed MN/MCA1 wt mice (Fig. 3f) led to increased IFNγ and granzyme-B expression in infiltrating CD8^+^ T cells (Extended Data Fig. 11i), effectively hindering tumor progression (Fig. 3f).

To investigate the role of cholesterol-induced RORγ activity in monocyte-to-macrophage differentiation driven by M-CSF and alternative polarization, we performed RNA-sequencing on RNA purified from wt or *Rorc*^-/-^ BM-derived macrophages (BMDMs) treated with IL-4 *vs.* untreated control, to induce M2 polarization in the presence or absence of cholesterol supplementation. Principal components analysis and unsupervised hierarchical clustering revealed distinct responses to cholesterol between wt and *Rorc*^-/-^ cells (Fig. 3g and Extended Data Fig. 12a). RORγ deficient cells had a similar transcriptional profile regardless of cholesterol supplementation. In contrast, cholesterol addition profoundly changed the transcriptional profile in wt macrophages, with a more pronounced effect in IL-4-treated cells (Fig. 3g). Specifically, IL-4-treated wt macrophages supplemented with cholesterol displayed the highest number of differentially expressed genes (DEGs) (1539 up-regulated and 1351 down-regulated) (Fig. 3h, i). Consistent with our findings *in vivo* (Fig. 3d, Extended Data Fig. 10f, k-m and 11b, f), the intersection analysis of gene sets identified in wt and *Rorc*^-/-^ macrophages revealed opposite modulation of gene expression upon cholesterol supplementation (Fig. 3h, i). Specifically, there was an inverse regulation of genes involved in the “interferon alpha/gamma response” pathway in wt *vs*. *Rorc*^-/-^ macrophages (Fig. 3h, i; Combo 2 and 8). Furthermore, while the “inflammatory response” and “TNF-signaling” pathways were only associated with down-regulated genes in wt cells (Fig. 3i; Combo 7 and/or 8), genes of the “TGF- beta signaling” pathway were up-regulated exclusively in wt macrophages in response to cholesterol treatment (Fig. 3h; Combo 3 and 6). In addition, we cultured peritoneal macrophages (PECs) from wt or *Rorc^-/-^* mice with TCM, supplemented or not with cholesterol. As shown in Extended Data Fig. 12b, RORγ-null PECs displayed decreased expression of M2 genes (*i.e.*, *Il4, Tgfb, Ccl22, Mrc1, Ccl2, Csf1*), while upregulating M1 genes (*i.e.*, *Tnfa, Cxcl9, Cxcl10, Cxcl11*).

In agreement with Extended Data Fig. 7a, the expression of the M-CSF receptor CD115 (*Csf1r*) was not affected by either cholesterol or RORγ deficiency (Extended Data Fig. 12c), whereas cholesterol-dependent CCR2 upregulation was reduced in the absence of RORγ (Extended Data Fig. 12d). TAM analysis corroborated RORγ-dependent regulation of CCR2 (Extended Data Fig. 12e). Since CCR2 and CSF1R/CD115 are chemotactic receptors, we assessed the migration of wt *vs. Rorc^-/-^* PECs, pre-conditioned with cholesterol for 24 h, in response to M-CSF, CCL2, or TCM. In line with the unchanged expression of CSF1R/CD115 (Extended Data Fig. 12c, e), the chemotaxis ability of cholesterol-conditioned *Rorc^-/-^* PECs was severely impaired in response to CCL2 and TCM, but not M-CSF (Extended Data Fig. 12f). Consistent with the M1 skewing observed in *Rorc^-/-^* PECs (Extended Data Fig. 12b), RORγ deficiency in TAMs or M-MDSCs from hypercholesterolemic mice enhanced the proliferation rate of co-cultured CD4^+^ and CD8^+^ wt T cells (Extended Data Fig. 12g). Likewise, BM-derived macrophages (BMDMs) from *Rorc*^-/-^ mice, conditioned with TCM plus cholesterol, displayed increased CD4^+^ and CD8^+^ T cell proliferation (Extended Data Fig. 12h). Collectively, these findings couple the cholesterol sensing function of RORγ to its role in driving the differentiation and tumor infiltration of M2-like immunosuppressive myeloid cells.

To dissect the relative contribution of RORγ and cholesterol to tumor development, we treated hypercholesterolemic wt or *Rorc*^-/-^ mice with an anti-PCSK9 antibody. Tumor progression was reduced to a similar extent in both anti-PCSK9-treated wt mice and untreated *Rorc*^-/-^ mice (Extended Data Fig. 13a, b). Moreover, the antitumor activity driven by RORγ deficiency was not further improved by anti-PCSK9 treatment, indicating that cholesterol lowering (Extended Data Fig. 13c) did not enhance the impact of RORγ-deficiency on tumor inhibition (Extended Data Fig. 13a, b) and the frequency of tumor-infiltrating M-MDSCs, TAMs, IMs, and AMs (Extended Data Fig. 13d-f). In this regard, the unchanged transition of CMPs to GMPs observed in the context of RORγ deficiency (Extended Data Fig. 11c and 13g) was consistent with the unaltered effects of cholesterol lowering under these conditions. Additionally, anti-PCSK9 treatment reduced the lipid content in TAMs without affecting CCR2, TNFα, MHC-II, and CD206 expression levels in RORγ−null animals (Extended Data Fig. 13h). Similar results were observed for IM and AM cell subsets (Extended Data Fig. 13f). In keeping with these findings, while anti-PCSK9 treatment of HCD-fed mice enhanced IFNγ and reduced both CTLA4 and PD-1 expression by CD8^+^ T cells, these effects were hindered by RORγ deficiency (Extended Data Fig. 13i). Furthermore, we confirmed the importance of myeloid cell-dependent suppression of specific antitumor immunity by depleting CD4^+^ and CD8^+^ T lymphocytes in HCD-fed TB mice, which curbed the antitumor activity of RORγ deficiency (Extended Data Fig. 13j, k).

Overall, these results demonstrate that the cholesterol-RORγ axis plays a critical role in myeloid cell-mediated suppression of specific antitumor immunity.

## Employing cholesterol-RORγ blockade as therapeutic approach

To validate the potential of targeting RORγ as a pharmacological intervention in hypercholesterolemia-induced protumoral myelopoiesis, we treated NCD- or HCD-fed mice with the RORγ inhibitor SR2211^3,58^ and evaluated MN/MCA1 tumor development. SR2211 treatment led to a substantial reduction in lung metastasis formation (Extended Data Fig. 14a) and only had a mild effect on primary tumor growth (Extended Data Fig. 14b), without altering cholesterol levels (Extended Data Fig. 14c). Furthermore, while SR2211 treatment of HCD-fed TB mice reduced both TAM accumulation and their CCR2 expression levels (Extended Data Fig. 14d), it increased the expression of TNFα and MHC-II, leaving CD206 expression unchanged (Extended Data Fig. 14e). Similar results were obtained in both IMs and AMs (Extended Data Fig. 14f). In good agreement with the observed decrease in TAMs in HCD-fed mice (Extended Data Fig. 14d), SR2211 also effectively blocked the transition of CMPs to GMPs, particularly under hypercholesterolemic conditions (Extended Data Fig. 14g). SR2211-induced reprogramming of TAMs, IMs, and AMs was paralleled by increased IFNγ and reduced PD-1 and CTLA4 expression levels in tumor-infiltrating CD8^+^ T cells (Extended Data Fig. 14h). The antitumor activity of the RORγ inhibitor was further demonstrated in K1735-M2 metastatic melanoma-bearing mice (Fig. 4a, b and Extended Data Fig. 14i), where SR2211 treatment led to a reduction in the frequency of TAMs, IMs, and AMs (Extended Data Fig. 14j, k), restored the TNFα^high^ M1 phenotype of pulmonary macrophages (Extended Data Fig. 14k), and reactivated the immune functions of CD8^+^ T cells (CD8^+^IFNγ^hi^PD-1^lo^) (Extended Data Fig. 14l). In line with the more pronounced protumoral phenotype of RORγ-proficient TAMs from hypercholesterolemic mice (Fig. 3d, f; Extended Data Fig. 10k and 11g), SR2211 treatment significantly improved the antitumor efficacy of anti-PD-1 immunotherapy in MN/MCA1 mice, particularly under HCD conditions (Fig. 4c and Extended Data Fig. 14m). Thus, these findings underscore the significant therapeutic potential of targeting the cholesterol-RORγ axis as an effective strategy to counteract protumoral myelopoiesis.

## Conclusions

Our work identifies RORC1/RORγ as a key immunometabolic sensor, linking tumor-induced dyslipidemias to myelopoietic alterations that fuel disease progression. Tumor-dependent induction of PCSK9 leads to hepatic LDL receptor (LDLR) downregulation, elevating circulating cholesterol levels and activating suppressive RORγ-dependent myeloid populations. Cholesterol-induced RORγ activation triggers the transition of CMPs to GMPs, resulting in increased intratumoral accumulation of TAMs and M-MDSCs, as well as of lung M-MDSCs, IMs, and AMs, through a CSF1R- and CCR2- dependent mechanism. Inhibition of these pathways restores specific antitumor immunity. Thus, while cholesterol is a fundamental component of phospholipid bilayers, which enables membrane biogenesis and proliferation of cancer cells^59^, the activation of RORγ, consequent to lipid dysmetabolism, triggers a protumoral myelopoiesis that establishes immunosuppressive conditions in the host, facilitating both metastatic spread and seeding. Indeed, hypercholesterolemia induces the expression of PD-L1 in TAMs, IMs, and AMs (Fig.2e, f, Extended Data Fig. 4l and Extended Data Fig. 5e), as well as of PD-1 in CD4^+^ and CD8^+^ lymphocytes (Extended Data Fig. 5j,k), and these effects are driven by RORγ (Extended Data Fig. 10k-m and Extended Data Fig. 14h, l).

The scientific literature on the connection between cholesterol and cancer is full of contradictory observations^60^ and the lipid alterations that we observed during tumor development can in principle be further affected by more advanced stages of the disease, since it is known that cachexia promotes an increase in lipolysis^61^.

Our work establishes that while tumor progression is fueled by cholesterol-driven protumor myelopoiesis, hypercholesterolemia may favor the efficacy of immune checkpoint inhibitors (ICIs) due to increased levels of PD-L1 (Fig. 2e, f and Extended Data Fig. 4l, 5e). While our observations should be reasonably extended and validated in other types of tumors as well, our conclusions are in agreement with recent evidence indicating a positive correlation between overweight and the efficacy of ICIs^49^.

As efforts intensify to develop strategies targeting suppressor myeloid cells, and recognizing the significance of PD-L1 as a biomarker for responsiveness to ICB treatment^62^, especially in obese/hypercholesterolemic cancer patients^49,63^, our findings provide crucial insights into the potential adjuvant role of RORC1/RORγ inhibitors and cholesterol-lowering drugs such as anti-PCSK9 agents.

## Methods

### Animals

All mice used in this study were 7–18 weeks of age, with both males and females included. Wild-type C57BL/6J and C3H/HeJ mice were obtained from Charles River Laboratories. *Pcsk9*- deficient mice (B6;129S6-*Pcsk9^tm1Jdh/^*J) and B6.129P2-*Kras^tm4Tyj^/J* (*Kras^LSL-G12D^*); *Trp53^tm1Brn^*/J (*p53^flox/flox^*) (KP) mice were purchased from The Jackson Laboratory. *Rorc* mutant mice (B6.129P2(Cg)-*Rorc^tm1Litt^*/J) were kindly provided by Dr. Dan Littman (New York University). *Rorc*^flox/flox^ (*Rorc*^fl/fl^, B6(Cg)-*Rorc^tm3Litt^*/J; The Jackson Laboratory) mice were crossed with *Lyz2-Cre* (B6.129P2-*Lyz2^tm1(cre)Ifo^*/J; The Jackson Laboratory) mice to generate *Rorc*^fl/fl^ *Lyz2-Cre* mice. All colonies were housed and bred in a specific pathogen-free animal facility at the Humanitas Research Hospital (Rozzano, Milan, Italy) in individually ventilated cages. Mice were housed under a 12/12-h light/dark cycle at an ambient room temperature (RT) of 22 ± 1 °C with 52–55% humidity. Mice were randomized based on sex, age, and weight. All procedures involving handling and care of C57BL/6J mice followed approved protocols by the Humanitas Research Hospital and were in compliance with national (Legislative Decree No. 116, G.U., suppl. 40, 02-18-1992 and No. 26, G.U., 03-04-2014) and international law and policies (EEC, Council Directive Nos. 2010/63/EU, OJ L 276/33 and 09- 22-2010; National Institutes of Health Guide for the Care and Use of Laboratory Animals, US National Research Council, 2011). The study received approval from the Italian Ministry of Health (Nos. 97/2014-PR and 25/2018-PR). Every effort was made to minimize the number of animals used and their suffering.

### Cell lines

The following cell lines were used: 3-MCA-derived sarcoma MN/MCA1; B16-F10 and K1735-M2 melanoma; Lewis lung carcinoma (LLC); and MC38 colon adenocarcinoma. B16-F10, LLC, Hepa1-6, and HEK293T cells were all purchased from ATCC. K1735-M2 cells were kindly provided by L. Carminati (Laboratory of Tumor Microenvironment, Istituto di Ricerche Farmacologiche Mario Negri IRCCS, Bergamo, Italy). The 3-MCA-derived sarcoma cell line MN/MCA1 was derived from stock periodically renewed through primary cells isolated from tumors implanted in wt mice. All cell lines were verified to be free from mycoplasma contamination.

### Cancer models

Mice were injected intramuscularly in their left hind limb with MN/MCA1 cells (1×10^5^ per mouse in 100 μL of PBS) or subcutaneously with 100 μL of PBS containing 5×10^5^ murine melanoma B16-F10 cells, 1×10^6^ MC38 cells, or 1×10^6^ LLC cells. When tumors became palpable (∼9-12 days after tumor cell injection), tumor growth was monitored three times per week with a caliper. ED and AD were established based on the metastatic rate in MN/MCA1 fibrosarcoma model, as described in previous experiments^3^ (see also Supplementary Fig. 1a). The primary tumor growth of the MN/MCA1 model was then used as reference to define ED or AD in the B16-F10, MC38, and LLC models. For spontaneous lung metastasis assay, C3H/HeJ mice were subcutaneously injected in the flank with 100 μL of PBS containing 1×10^5^ murine melanoma K1735-M2 cells. The growth of the primary tumors was monitored with a caliper, and when the tumors reached 800-1,000 mm^3^, mice were anesthetized, and the tumors surgically removed. For the analysis of spontaneous lung metastasis, all mice were autopsied 28 days after surgery^43^. The *Kras*/*p53*-driven lung cancer model was generated as previously described^33^. Briefly, *K-ras^LSL-G12D/+^;p53*^fl/fl^ (KP) mice were intranasally inoculated with 2.5×10^7^ infectious particles of an adenoviral vector constitutively expressing the Cre recombinase (Ad5-CMV-Cre) to induce sporadic mutations and lung tumor development. MN/MCA1 mCherry^+^ (1×10^5^ cells per mouse in 100 μL of PBS) were injected intramuscularly in the left hind limb of wt or *Rorc*^-/-^ mice. Blood was collected 20 days after tumor cell injection and analyzed by RT–PCR for mCherry expression and by FACS to quantify circulating mCherry^+^ MN/MCA1 tumor cells.

### Patients

To measure cholesterol and PCSK9, human peripheral blood sera were isolated from non-small cell lung cancer (NSCLC), colorectal cancer (CRC), breast cancer (BRC), pancreatic ductal adenocarcinoma (PDAC), biliary tract cancer (BTC), and pancreatic neuroendocrine tumor (PNET) patients at both early (I/II) and advanced (III/IV) stages according to the AJCC cancer staging systems, eighth edition, at the Humanitas Research Hospital. For the CRC patients cohort (n=65), HDL and LDL measurements were performed according to volume sample availability (n=31). The patients for PCSK9 measurement were randomly selected based on the residual sample volume available. Frozen biopsies of non-neoplastic liver tissues from metastatic CRC patients (CRC) and healthy liver tissues from patients with benign lesions (CTRL) (i.e., adenomas, angiomas, and hyperplasia) were provided by Biobanca Humanitas. The samples were selected from non-obese (BMI < 30) and non-dyslipidemic subjects (Supplementary Table 3). Human peripheral blood leukocytes were isolated from NSCLC patients at stages III/IV, healthy normocholesterolemic subjects or hypercholesterolemic patients with dyslipidemia and stable coronary artery disease. Samples from the cohorts described in this study were obtained with informed consent from the patients. The study was conducted according to the Declaration of Helsinki Principles and following approval by the Ethics Committee of Humanitas Research Hospital, Milan, Italy. More detailed information about the patients, including tumor staging and therapy, can be found in Supplementary Table 3.

### Blood cholesterol measurement

Mouse blood samples were collected at the described time points by puncturing the facial vein into microcentrifuge tubes and kept on ice for 20 min. They were then centrifuged for 20 min at 13,000 rpm at 4°C to separate the serum. Human blood samples were obtained by patients and healthy donors and collected in BD Vacutainer^®^ SST^TM^ tubes. The samples were allowed to clot for 20 min at RT and then centrifuged for 15 min at 2000 rpm at RT to separate the serum from the cells. Serum samples were measured for total, LDL, and HDL cholesterol content at the Clinical Analysis Laboratory of Humanitas Hospital. The test assay kits used were purchased from Abbott (Cholesterol 7D62; Direct LDL 1E31-20; Ultra HDL 3K33-21).

### ELISA measurement

Murine and human IL-6 and IL-1β were quantified using R&D DuoSet® ELISA Development Systems. Human and murine PCSK9 levels were measured with Abcam kits (ab209884; ab215538). Murine VLDL was quantified using a Biorbyt kit (orb692924).

### Lung histology

Lungs were collected following intracardiac perfusion with cold PBS, formalin fixed for 24 h, dehydrated, and paraffin embedded for histological analysis. Histology was performed on two or three longitudinal serial sections spaced 100-150 μm apart, with a width of 7 μm, from each lung. The sections were stained with hematoxylin and eosin (H&E) and scanned using a VS120 Dot-Slide BX61 virtual slide microscope (Olympus Optical). The areas of lung lesions were identified by manually tracing the perimeter of lesions using Image Pro-Premiere software 9.2 (Media Cybernetics). In graphs showing percentages of lung metastatic areas, each dot value represents the mean area of two contiguous sections obtained from each longitudinal level of a single lung.

### Bone marrow transplantation

Bone marrow transplantation (BMT) was performed by intravenously injecting 5×10^6^ BM cells into lethally irradiated (two doses of 4.50 Gy each) 8-week-old mice. For *Rorc*-deficient BM cell transfer, CD45.1 recipient wt mice were transplanted with BM cells from either wt or *Rorc*^-/-^ CD45.2 donors. BM engraftment was verified 4 weeks later through FACS analysis (Supplementary Figure 1e) of blood cells stained with CD45.1 and CD45.2 antibodies (Supplementary Table 4).

### TAM adoptive transfer

FACS-sorted TAMs (2×10^5^) from wt or *Rorc*^-/-^ MN/MCA1-bearing mice on NCD were intramuscularly injected together with 1×10^5^ MN/MCA1 tumor cells into the hind limb of wt mice fed NCD or HCD (Dose 1). After 13 days of growth, when tumors were palpable, a second injection of FACS-sorted wt or *Rorc*^-/-^ TAMs (1×10^5^) was administered intratumorally. At day 23, mice were sacrificed and further analyses were performed.

### *In vivo* treatments

To model diet-induced obesity and hypercholesterolemia, 7 to 8-week-old mice were assigned to either a high-fat, high-cholesterol (HCD; ssniff ® EF D12079: 21% w/w butterfat, 0.2 % w/w cholesterol) or a standard low-fat, low-cholesterol (normal chow diet, NCD; Special Diet Service VRF1(P): 5% w/w fat, 0.006% w/w cholesterol) irradiated diet. After 8 weeks, the animals were injected with tumor cells and maintained on their respective dietary regimen throughout the experiments. BODIPY-Cholesterol (TopFluor® Cholesterol, Avanti Polar Lipids) was dissolved in a mixture of DMSO and Corn oil (1:1). Mice were fasted 24 h before sacrifice and concomitantly administered BODIPY-Ch (250 μg/mouse) using an intragastric syringe. Starting from day 10 after tumor cell injection, MN/MCA1-bearing mice received intraperitoneal treatment with the following reagents: anti-IL6 monoclonal antibody (BioXcell, Clone MP5-20F3) at a dose of 200 μg twice a week; anti-IL1 (anakinra, Sobi, Stockholm, Sweden) at a dose of 200 μg twice a week; anti-PCSK9 monoclonal antibody mAb1 (kindly provided by Amgen Inc, Thousand Oaks, CA, USA) at a dose of 200 μg per mouse twice a week (MN/MCA1 tumor) or once a week (KP mice); anti-CSF1R monoclonal antibody (BioXcell, Clone AFS98) at an initial dose of 400 μg per mouse, followed by twice-weekly doses of 200 μg for the duration of the experiment; anti-CCR2 inhibitor (Tocris) at a dose of 75 μg per mouse twice a week; and the RORγ inverse agonist SR2211 (Tocris) at a dose of 50 μg per mouse twice a week. When specified, starting from the day before tumor injection, mice received an intraperitoneal injection of 300 μg of anti-mouse CD4 (Rat Anti-Mouse CD4 Monoclonal Antibody, Unconjugated, Clone GK1.5, BioXcell) and 300 μg of anti-mouse CD8 (Rat Anti-Mouse CD8α Monoclonal Antibody, Unconjugated, Clone 2.43, BioXcell) once a week. FACS analysis of peripheral blood samples confirmed the depletion of CD4^+^ and CD8^+^ cells over a period of 7 days. When specified, MN/MCA1-bearing mice were intraperitoneally injected with 100 μg of anti-PD-1 antibody (BioXcell, Clone RPM1-14), twice a week, starting 10 days after tumor cell injection, either alone or in combination with SR2211 treatment. For the spontaneous metastasis assay, SR2211 (50 μg, intraperitoneally, twice a week) was administered either before or after surgical removal of K1735-M2 primary melanomas or only after surgical removal of primary melanoma^43^.

### Quantitative RT-PCR

Liver tissues from mice and patients, small intestine and adipose tissues from mice, and MN/MCA1 tumor tissues were disrupted by TissueLyser II (QIAGEN) with stainless steel beads following the manufacturer’s instructions. Total RNA from tissues, primary hepatocytes, and PECs was extracted using TRIzol reagent (Invitrogen) following the manufacturer’s instructions. Total RNA from FACS-sorted TAMs was extracted using an RNA/DNA isolation kit (Zymo). Complementary DNA was synthesized by reverse transcription using a High-Capacity cDNA Archive Kit (Applied Biosystems), and quantitative real-time PCR was performed using SybrGreen PCR Master Mix (Applied Biosystems) through ViiA-7 Fast Real-Time System (Applied Biosystems). Data were processed using ViiA™ 7 Software (Applied Biosystems) and analyzed with the 2^(−ΔCT)^ method. Data were normalized to *β-actin* or *18S* expression and represented as fold change over control. Primer sequences used in the manuscript are available upon request.

### Immunoblotting

Liver tissues were disrupted by TissueLyser II (QIAGEN) and lysed in radioimmunoprecipitation assay (RIPA) buffer supplemented with protease and phosphatase inhibitors (Sigma) for 2 min at 25 Hz at 4°C, repeated twice. The lysates were centrifuged at 13,000 rpm at 4 °C for 15 min, and the supernatants were run on SDS–PAGE (30 μg of total proteins per lane) and transferred to polyvinyldifluoride (PVDF) membrane. Immunoblotting was performed using the following antibodies: (primary) rabbit anti-mouse LDLR (Invitrogen), mouse anti-mouse PCSK9 (Invitrogen), mouse anti-mouse vinculin (Santa Cruz), goat anti-mouse actin (Santa Cruz); (secondary) goat anti-mouse and anti-rabbit IgG horseradish peroxidase (HRP)-conjugated (Amersham). ChemiDoc™ Touch Imaging System and Image J software (National Institutes of Health, NIH) were employed for image acquisition and densitometric analysis of blots.

### Lipid extraction from feces and TBA assay

Fecal lipids were extracted using a modified Folch method, as previously described^64^. Briefly, feces were collected from MN/MCA1-bearing mice housed for 4 days (from day 18 to day 22 of tumor growth) or from age-matched tumor-free mice. Three replicates of 1 g of feces per experimental group were homogenized in 5 mL of saline in a 15- ml tube. To this, 5 ml of a chloroform/methanol mixture (2:1) was added, and the mixture was thoroughly mixed. After centrifugation at 400×g for 10 min at RT, the separation into three layers occurred, and the bottom layer, containing the organic phase, was recovered and dried. Lipid extracts were resuspended in 300 μl of ethanol. The solution was analyzed by the commercially available total bile acid (TBA) assay kit purchased from Abcam (ab239702), following the manufacturer’s instructions.

### Murine primary hepatocytes isolation

Primary hepatocytes were isolated from 12-week-old mice anesthetized and sacrificed by bleeding as previously described^65^. Briefly, liver was first perfused with Hank’s balanced salt solution (HBSS) without magnesium or calcium supplemented with 0.5 mM EGTA, and then with digestion medium (DMEM low-glucose with 100 U/ml penicillin/streptomycin, 15 mM HEPES, 0.8 mg/mL of collagenase-IV). The digestion was performed at 37°C for about 7-8 min, after which the liver was excised, finely minced using forceps into a Petri dish containing digestion medium under sterile conditions, and then filtered through a 70- μm cell strainer. The cells were centrifuged at 1,000 rpm for 5 min, washed twice with DMEM:HAM’S F-12 (1:1) with 100 U/ml penicillin/streptomycin, 2 mM glutamine, without FBS. Cell viability was assessed by trypan blue exclusion. Viable cells were resuspended in DMEM:HAM’S F-12 (1:1) supplemented with 10% FBS and then seeded in 6-well plates (5×10^5^ cells/well) that had been pre-coated the day before with type I collagen (Sigma) dissolved in 0.02 N acetic acid. Cells were incubated for 2 h at 37°C. After the cells had attached, they were washed and incubated in fresh medium for 1 h before treating them with the following compounds: TCM (30%), cholesterol (50 μg/ml), or mouse serum (20%). Cells were then incubated for 24 h.

### Metabolomic analyses of liver tissues

Mevalonate quantification was assessed as previously described^66^. Briefly, liver samples (15 mg) were homogenized in water, and HCl 6 M was added. The samples were then incubated for 30 min in the dark at RT to enable complete lactonization of mevalonate, eluted with methanol, and centrifuged at 4°C at 14,500×g for 15 min. The supernatants were collected and dried using a speed vacuum concentrator. Dried samples were then resuspended in methanol and quantified. For squalene quantification, livers were homogenized in water, and hexane was added. After incubating for 30 min at 15 rpm, the supernatants were collected and then dried, concentrated with a speed vacuum concentrator, and finally resuspended in hexane and analyzed. Each sample was spiked with internal standards for data normalization and instrument stability monitoring. Quality control (QC) samples containing spiked standards were also acquired. Mevalonate and squalene were quantified using a GCXGC/ TOFMS (Leco Corp., St. Josef, MI, USA). The first-dimension column was a 30 m Rxi-5Sil MS capillary column (Restek Corp., Bellefonte, PA, USA), with an internal diameter of 0.25 mm and a stationary phase film thickness of 0.25 mm. The second-dimension chromatographic column was a 2 m Rxi-17Sil MS (Restek Corp., Bellefonte, PA, USA) with a diameter of 0.25 mm and a film thickness of 0.25 mm. High-purity helium (99.9999%) was used as the carrier gas, with a flow rate of 1.4 mL/min. For both molecules, 1 mL of sample was injected in splitless mode at 250°C. Sample analysis was performed using two different temperature programs. For mevalonate, the initial temperature program started at 70°C and increased gradually at a rate of 4°C/min until reaching 280°C. For squalene, the initial temperature program began at 150°C and was raised to 250°C at a faster rate of 40°C/min, where it was maintained for 2 min. Then, it was further ramped to 285°C at 5°C/min and finally heated to 300°C at a rate of 15°C/min. Throughout both analyses, the secondary column was consistently maintained at a temperature 5°C higher than the GC oven temperature of the first column to ensure proper functioning and accurate results. Electron impact ionization (70 eV) was applied, with the ion source temperature set at 250°C. The mass range was 25 to 550 m/z, with an extraction frequency of 32 kHz and acquisition rates of 200 spectra/s. The modulation period for the entire run was 4 s. The modulator temperature offset was set at +15°C relative to the secondary oven temperature, while the transfer line was set at 280°C. The chromatograms were acquired in total ion current mode, and m/z 69 and m/z 58 at 616 and 358 s were selected to identify squalene and mevalonate, respectively. The raw data were processed with ChromaTOF version 5.31. Mass spectral assignment was performed by matching data with NIST MS Search 2.3 libraries and FiehnLib. m/z and retention times were also confirmed with standards. External calibration curves and internal standards were used for quantification of mevalonate and squalene.

### Lipidomic analysis

Lipids were extracted from 5×10^5^ FACS-sorted TAMs using a 1 mL solution of 75:25 IPA/H2O, after the addition of deuterated lipid standard (Splash Lipidomix®, Merck, Berlin). The samples were vortexed and sonicated for 2 min and then incubated for 30 min at 4°C under gentle and constant shaking. To remove debris and other impurities, the samples were centrifuged for 10 min at 3,500g at 4°C. Subsequently, 1 mL of supernatant was dried using a speed vacuum concentrator, and then reconstituted in 100 μL of MeOH containing the internal standard CUDA (12.5 ng/mL). The reconstituted lipids were analyzed by UHPLC Vanquish system coupled with Orbitrap Q-Exactive Plus (Thermo Scientific). A reverse phase column was used for lipid separation (Hypersil Gold™ 150 × 2.1 mm, particle size 1.9 μm), with the column maintained at 45°C at a flow rate of 0.260 mL/min. Mobile phases and mass spectrometry parameters were set as previously reported^67^. The acquired raw data from the untargeted analysis were processed using MSDIAL software version 4.24 (Yokohama City, Kanagawa, Japan). This involved the detection of peaks, MS2 data deconvolution, compound identification, and the alignment of peaks across all samples. To obtain an estimated concentration expressed in μg/mL, the normalized areas were multiplied by the concentration of the internal standard. An in-house library of standards was also used for lipid identification. Statistical analysis was performed using MetaboAnalyst 5.0 software (www.metaboanalyst.org) and GraphPad Prism v. 8.

### FACS analysis of murine samples

Primary tumors and explanted lungs were cut into small pieces, disaggregated with 0.5 mg/ml collagenase-IV and 150 IU/ml DNase-I in RPMI 1640 for 30 min at 37°C, and filtered through a 70-μm strainer. Splenocytes were collected from spleens after disaggregation and filtration through a 70-μm strainer. BM cells were isolated from the tibias and femurs of both TF and TB mice. Whole blood samples were collected from the facial vein or heart in EDTA-coated collection tubes and directly processed using a standardized red blood cell lysis protocol. The resulting cells were resuspended in HBSS supplemented with 0.5% FBS. Staining was performed at 4 °C for 20 min using the antibodies listed in Supplementary Table 4. In addition, LipidTOX™ Green Neutral Lipid Stain (1:200, Invitrogen) was used after fixing the cells with PFA 1% for 30 min. LipidTOX staining was performed in PBS for 30 min up to 2 h at RT. Cell viability was determined using either Zombie Aqua™ Fixable Viability Kit (1:800, Biolegend) or LIVE/DEAD™ Fixable Violet Dead Cell Stain Kit (1:1,000); negative cells were considered viable. For intracellular staining of TNFα, iNOS, CD206, IDO1, IFNγ, RORγ, PCSK9, and LDLR, a Foxp3/Transcription Factor Staining Buffer Set (eBioscience) was used. Expression of TNFα and IFNγ was analyzed by flow cytometry following 4 h of treatment with brefeldin A (5 μg/ml), PMA (50 ng/ml), and ionomycin (1 μg/ml). PCSK9 mouse and LDLR rabbit monoclonal antibodies were followed by incubation with secondary goat anti-mouse Alexa Fluor 647-conjugated and goat anti-rabbit Alexa Fluor 488-conjugated antibodies (ThermoFisher), respectively. Cell detection was performed using either BD FACSCanto II, BD LSRFortessa or BD FACSymphony, and data analyzed with FlowJo 9.9.6 software. Results are reported as mean fluorescence intensity (MFI) or MFI normalized to isotype control or fluorescence minus one (ΔMFI). To assess BODIPY-Cholesterol fluorescence in tissues or cells, flow cytometry (AF488 channel) was performed, and relative MFI was normalized to the biological vehicle control (ΔMFI). For t-distributed stochastic neighbor embedding (tSNE) analysis^68^, a unique computational barcode was assigned to single samples. Events gated on live CD45^+^ cells were subsequently concatenated and visualized with tSNE visualization (Barnes-Hut implementation; iterations, 1000; perplexity, 20; initialization, deterministic; theta, 0.5; eta: 200) was carried out. The expression of the following markers was analyzed: CD11b, CD11c, CD103, CD64, Ly6C, Ly6G, and F4/80.

### FACS analysis of human PBMC samples

Human PBMCs were obtained through Ficoll density gradient centrifugation (Ficoll-Paque PLUS, GE Healthcare). Multicolor staining for cell populations was performed on cryopreserved samples. Cells were thawed in RPMI medium supplemented with 10% FBS, incubated at 37°C for 2 h, and then washed and resuspended in staining buffer (HBSS with 0.5% FBS). Staining was performed at 4°C for 20 min. using the antibodies listed in Supplementary Table 5. Cell viability was determined by LIVE/DEAD™ Fixable Near-IR Dead Cell Stain Kit (1:1000, ThermoFisher), where negative cells were considered viable. A Foxp3/Transcription Factor Staining Buffer Set (eBioscience) was used for intracellular staining of RORγ. Cells were detected using BD LSR Fortessa and analyzed with FlowJo 9.9.6 software.

### Purification of mouse leukocytes

PECs were harvested by peritoneal lavage from mice injected with 1 mL of 3% (weight/vol) thioglycollate medium (Difco) 5 days before isolation as previously described^40^. PECs were cultured in RPMI 1640 medium containing 10% FBS, 2 mM glutamine, and 100 U/ml penicillin/streptomycin, with the addition of TCM (30%) with or without cholesterol (50μg/ml) for 48 h. Isolation of TAMs was performed as previously described^40^. Briefly, 18-21 days after tumor cell injection, primary tumors were cut into small pieces, disaggregated with 0.5 mg/ml COL IV and 150 U/ml DNase I in RPMI 1640 for 30 min at 37 °C, and filtered through a strainer. The cell suspension was enriched in CD11b^+^ cells through positive selection using CD11b microbeads (MACS, Miltenyi Biotec). The purity of CD11b^+^ cells was ∼90% as determined by FACS. To obtain high-purity TAM populations, CD11b^+^ cells were further stained (Live/Dead-Pacific Blue, CD45-FITC, CD11b-PerCP-Cy5.5, Ly6C-PE-Cy7, Ly6G-APC, F4/80-PE) and sorted using a FACSAria cell sorter (BD Bioscience). Myeloid progenitor cells (CMPs and GMPs) were isolated from the BM of MN/MCA1-bearing mice after 23 days of tumor growth. Briefly, BM cells were flushed from femurs and tibias of different mice and then pooled for each experimental group. To obtain high-purity CMP and GMP populations, after red blood cell lysis, the cells were stained (Live/Dead-APC-Cy7, cKit-PE-Cy7, Lin-eFluor450, Sca1-BV711, CD34-FITC, CD16/32-APC) and sorted on a FACSAria cell sorter. The purity of each sorted population was ≥ 95%. The resulting sorted cells were processed for adoptive transfer, lipid immunofluorescence staining, or mRNA extraction. BM cells were collected from femurs and tibias of healthy tumor-free C57BL6/J mice and differentiated for 6 days into BMDMs in RPMI 1640 medium (10% FBS, 2 mM glutamine, 100 U/ml penicillin/streptomycin) supplemented with 30 ng/ml of M-CSF (plus cholesterol at 50 μg/ml, when described). After 3 days of culturing in the M-CSF-supplemented medium, BMDMs underwent medium refreshment. Following a total culturing period of 6 days, cells were stimulated with IL-4 (20 ng/ml) for 20 h, or with TCM (30%) for 48 h after which they were analyzed as described above.

### Neutral lipid and cholesterol immunofluorescence

FACS-sorted TAMs, CMPs, or GMPs were seeded on Poly-L-Lysine (Sigma-Aldrich) coated sterile rounded glasses at a density of 2×105 cells/ml in RPMI medium and incubated for 2 h at 37°C. Cells were then fixed with 4% PFA for 10 min at RT. Cells were washed twice with PBS and stained with fluorescent lipid staining reagents: LipidTOX™ Green (1:500, Invitrogen), for total neutral lipids, or Filipin-III solution (1:20, Sigma-Aldrich), for cellular cholesterol. The staining was performed for 1 h at RT. For LipidTOX-stained cells, nuclei were counterstained with DAPI (Invitrogen), whereas for Filipin-III-stained cells, nuclei were visualized with SYTO Green (Invitrogen). Coverslips were mounted using the antifade medium FluorPreserve Reagent (EMD Millipore) and analyzed with an SP8 Laser Confocal Microscope (Leica) equipped with fine-focus oil immersion lens (×4/1.3 numerical aperture (NA)) and operated with lasers with 405 and 488 excitations. The intensity of fluorescence was analyzed using LAS X software (Leica).

### Cell treatments

To obtain MN/MCA1 TCM, digested MN/MCA1 tumors from wt mice were seeded at a density of 4×10^6^ cell/ml in RMPI 1640 supplemented with 10% FBS, 2 mM glutamine, and 100 U/ml penicillin/streptomycin. After 24 h of incubation, the supernatant was collected and filtered through a 0.2-μm filter. For the preparation of mouse serum for cell treatment, mice were anesthetized and sacrificed by bleeding. Blood was collected with a syringe via heart puncture to maintain sterility. The collected blood was centrifuged for 20 min at 13000 rpm, 4°C to separate the serum, which was then heat-inactivated for 15 min at 55°C and filtered as above. For cell stimulation, the following compounds and murine recombinant cytokines were used: cholesterol, desmosterol, LDL cholesterol, and the RORγ agonist SR0987 (all purchased from Sigma-Aldrich); IL-6, IL-1β, IL-4, M-CSF, and CCL2 (all purchased from Peprotech). Hepa1-6 cells were treated for 24 h with TCM (30%), IL-6 (10 ng/ml), or IL-1β (10 pg/ml), along with anti-IL6 (300 ng/ml) and anti-IL1 (300 ng/ml) where applicable. *In vitro* administration of BODIPY-Cholesterol (0.5 μg/ml) was performed 3 h before cell collection. Subsequently, the cells were fixed with PFA 1% and subjected to FACS analysis. MN/MCA1 cells were treated *in vitro* with cholesterol (50 μg/ml) and/or anti-PCSK9 mAb (300 ng/ml). The cell cycle was then assessed by DAPI staining. Briefly, cells were washed with PBS, fixed with 80% ethanol, and permeabilized for 10 min with 0.1% Triton X-100 (PBS). Nuclei were stained with 1 μg/ml DAPI (PBS) for 3 min. Alternatively, cell viability was measured by Annexin V-FITC/PI-staining. Samples were analyzed by flow cytometry.

### RNA sequencing and analysis

Total RNA was purified from BMDMs treated as described above using the Zymo Research kit (No. R2050) according to manufacturer’s instruction. The quality of the RNA was assessed using a High Sensitivity RNA ScreenTape Assay with a 4200 TapeStation System (Agilent). mRNA-seq library preparation was performed with SMART-Seq® v4 PLUS Kit (R4000752, Takara Bio). Single-end multiplexed libraries were sequenced using the NextSeq 2000 instrument (Illumina, San Diego, US), resulting in approximately 76±14 million reads/sample. The 75-bp single-end reads were aligned to the GENCODE *Mus musculus* reference genome (build GRCm38/mm10) using STAR v2.7.2b^69^. Raw read counts were normalized with TMM implemented in edgeR^70^, and low-expressed genes were filtered out with the filterByExpr function(min. count=19). Differential expression analysis of read counts was performed using voom, lmFit, and eBayes (robust=T) function of limma v.3.46^71^ package in R. Significant differential genes were chosen based on an FDR < 0.05 and |log2FC| > 0.5. Hierarchical clustering of significantly modulated genes was performed using the hclust and dist R functions on log2 CPM. Clustering was performed with the ward.D method and Pearson’s correlation distance to generate a heatmap using pheatmap^72^, with row scaling applied. Principal components analysis was performed using PCA function of FactoMineR v2.7^73^ package in R. The Upset plots were obtained using the R package UpSetR^74^. Over-representation analysis (ORA) was performed using enrichR^75^ with the “MSigDB Hallmark 2020” gene-set library, and the results were visualized using the “ggplot2” package.

### T cell suppression assay

Splenocytes from wt mice were labeled with 3 μg/ml of CellTrace CFSE (Life technology) for 10 min at 37 °C protected from light. Subsequently, cells were washed, resuspended in RPMI 1640 complete medium, and seeded on wells coated with anti-CD3 (2 μg/ml) and anti-CD28 antibodies (3 μg/ml). Myeloid cells (*i.e.*, TAMs and M-MDSCs FACS-sorted, and BMDMs) were co-cultured with CFSE-labeled splenocytes at different ratios of splenocyte:myeloid cells. After 72 h of co-culture, cells were collected, stained with anti-CD4 and anti-CD8 antibodies, and analyzed by flow cytometry. CellTrace signal from anti-CD3/CD28-activated lymphocytes, which were not co-cultured with myeloid cells, was used to evaluate cell proliferation.

### Plasmid transfection and luciferase assay

A plasmid vector constitutively expressing the *Rorc* gene under the control of the CMV promoter (pCMV6-*Rorc*) (Origene, No. MR222309) was co-transfected with a plasmid expressing the luciferase gene (*Luc*) under the control of the minimal promoter of the *Il17a* gene (minIL17prom-Luc) (Addgene, No. 20124). Transfections were performed using calcium phosphate method. At 18-h post transfection, HEK293T cells were washed, fresh medium was added, and the cells were allowed to rest for 1 h before being treated with SR0987 (20 μM), desmosterol (20 μM), cholesterol (20 μM), LDL (50 μg/ml), mouse serum (20%), TCM (30%), or SR2211 (20 μM). After 24 h, the cells were collected and subjected to luciferase assay following the manufacturer’s instructions (Promega, E1500). Luminescence was measured through Synergy 2 (BioTek).

### PEC migration

PECs were seeded for 18 h in low-adherence plates in RPMI 1640 with 10% FBS, 2 mM glutamine, and 100 U/ml penicillin/streptomycin, with or without the supplementation of cholesterol (50 μg/ml). Cells were then collected, washed, resuspended in RPMI 1640 medium with 1% FBS, and seeded (2×10^5^) on the upper chamber of a 5-µm-pore transwell insert (Corning, NY) in 24-well plates, while the bottom chamber was filled with RPMI 1640 with 10% FBS in the presence or absence of M-CSF (50 ng/ml), CCL2 (50 ng/ml), TCM (30%), with or without cholesterol (50μg/ml). After 5 h of incubation, the chambers were fixed and stained with Diff-Quik (Baxter).

### Statistical information

Statistical analysis was performed using GraphPad Prism v.8.0 software. P- values below 0.05 were considered statistically significant. The results are represented as mean ± SEM, unless otherwise stated. Statistical significance is indicated in each graph except for the heatmap graphs, for which the statistical analysis is provided in Supplementary Table 1. RNA- seq analysis is described in the corresponding method section above. For the experiments with more than two groups and two factors, two-way analysis of variance (ANOVA) was applied, and Šidák’s or Tukey’s multiple-comparisons tests were performed based on the nature of the comparison. Data with more than two independent groups were analyzed using one-way ANOVA with Šidák’s or Tukey’s multiple-comparisons correction, depending on the specific experimental design. When comparing only two groups, an unpaired two-tailed *t*-test was utilized to assess statistical significance. The number of biological replicates and the statistical methods used are specified in all figure legends.

### Data availabity

RNA-seq data supporting the findings of this study are available from the NCBI BioProject database (https://www.ncbi.nlm.nih.gov/bioproject/) under accession code PRJNA1000461. A reviewer link has been created for BioProject PRJNA1000461 at the link https://dataview.ncbi.nlm.nih.gov/object/PRJNA1000461?reviewer=s1e8t1fc1un7tfrkcihqgg9ck8.

## Acknowledgements

This work was supported by Associazione Italiana per la Ricerca sul Cancro (AIRC) IG (Nos. 19885 to A.S.); AIRC 5×1000 no. 22757; Fondazione Cariplo and Ministero Università Ricerca (project No. 2017BA9LM5_001); Associazione “Augusto per la Vita” Novellara (RE); and Associazione “Medicine Rocks”, Milano. We thank doctors Arianna Felicetta, Barbara Bottazzi, Annalisa Del Prete, Tiziana Schioppa, Sergio Marchini and the Genomics Facility (HuGE) at Humanitas Research Hospital for the technological and intellectual support provided in various phases of the work. We thank the Mario Negri Institute for Pharmacological Research, Milan, for the kind hospitality offered to us in some experimental phases.

## Author contributions

A.B. and F.M.C. were actively involved in designing and conducting most of the experiments. M.I. provided in vivo evidence on the protumor role of myeloid-specific RORC1 in response to cholesterol. C. Pandolfo and V.G. provided technical support with murine samples and collection and management of blood samples from NSCLC patients. M.N.M. performed the biochemical analysis of blood cholesterols. S.S. performed the bioinformatics analysis of genomic profiles. D.P. contributed plasma samples and liver biopsies from colorectal cancer patients, as well as blood samples from cancer patients. M.S. and C.A. managed the animal experimentation and performed the determination of inflammatory cytokines in mice and patient tissues. M.M. performed the metabolomic and lipidomic analyses. G.F. played a significant role in selecting NSCLC patients and organized the relative blood collection, under stringent clinical criteria. C. Panico and G.C. contributed by providing blood samples from hypercholesterolemic non-cancer patients. A.B. and A.S. drafted the manuscript. A.S. provided the guiding hypothesis of the work and contributed to the experimental design and overall supervision of the research.

## Figures and figure legends

**Extended Data Fig. 1.**
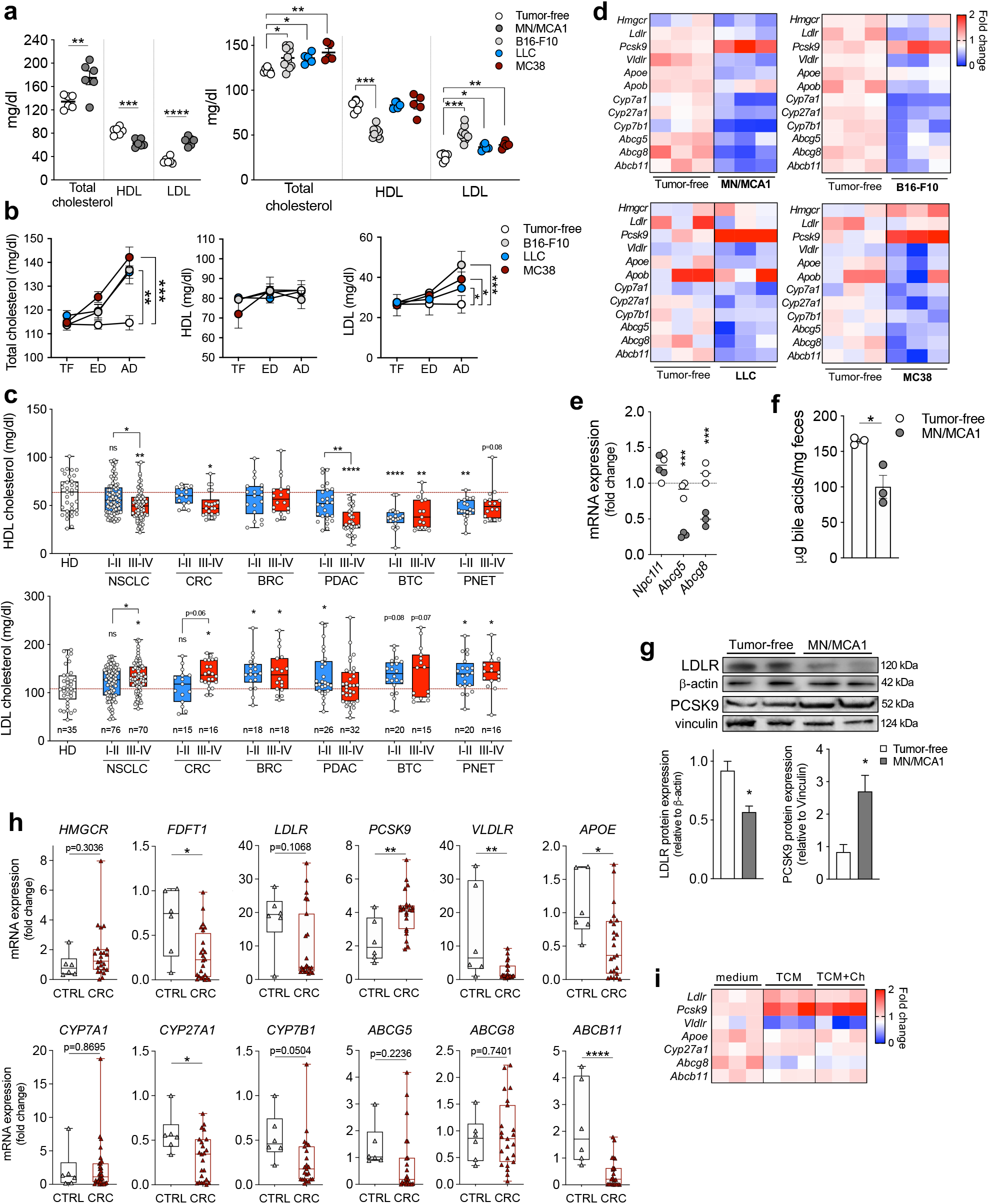
Tumor progression alters cholesterol homeostasis. **a,** Total, HDL, and LDL blood cholesterol levels in mice bearing MN/MCA1 fibrosarcomas (left) (n=6), B16-F10 melanomas (n=11), or LLC lung (n=5) or MC38 colon (n=5) cancers (right), at advanced disease (AD) stage, compared to tumor-free, age- and sex-matched control mice (n=6). **b,** Total, HDL, and LDL blood cholesterol levels in B16-F10, LLC, and MC38 mice at early (ED) and AD stages *vs.* tumor-free (TF) mice (n=5). **c,** HDL (top) and LDL (bottom) cholesterol levels in patients with early (I-II) *vs.* advanced (III-IV) NSCLC (n=76 *vs.* n=70), CRC (n=15 *vs.* n=16), BRC (n=18 *vs.* n=18), PDAC (n=26 *vs.* n=32), BTC (n=20 *vs.* n=15), and PNET (n=20 *vs.* n=16) in comparison with healthy donors (HD, n=35). **d,** Heatmap representing the expression levels of cholesterol metabolism genes in the livers of MN/MCA1, B16-F10, LLC, and MC38 mice at AD *vs.* TF mice (n=3). **e,** mRNA expression levels of *Npc1l1*, *Abcg5*, and *Abcg8* genes in the small intestine of MN/MCA1 mice at AD *vs.* TF mice (n=3). **f,** Total bile acid (TBA) quantification into lipids extracted from the feces of MN/MCA1 *vs.* TF mice (n=3). **g,** Immunoblot analysis of LDLR and PCSK9 protein in livers from AD MN/MCA1 *vs.* TF mice (n=2). **h,** mRNA expression levels of cholesterol metabolism genes in healthy liver parenchyma from CRC patients (n=23) compared to healthy liver parenchyma from patients with benign lesions (n=6). **i,** Heatmap representing the relative mRNA expression levels of cholesterol metabolism genes in primary unstimulated hepatocytes or stimulated with MN/MCA1 tumor-conditioned medium (TCM) with or without cholesterol (Ch) supplementation (n=3). Data are representative of at least four (**a**) or two (**d,** top left; **g**) independent experiments; as for **b, d, e, f, i,** one experiment was performed. **a, b, e-g,** Data are presented as mean ± SEM; **c, h,** Box-and-whisker min-to-max plots. \**P* < 0.05, \*\**P* < 0.01, \*\*\**P* < 0.001, \*\*\*\**P* < 0.0001 between selected relevant comparisons. **a, e-h,** Unpaired two-tailed *t*-test. **b, i,** Two-way ANOVA with Tukey’s multiple comparisons test. **c,** One-way ANOVA with Tukey’s multiple comparisons test.

**Extended Data Fig. 2.**
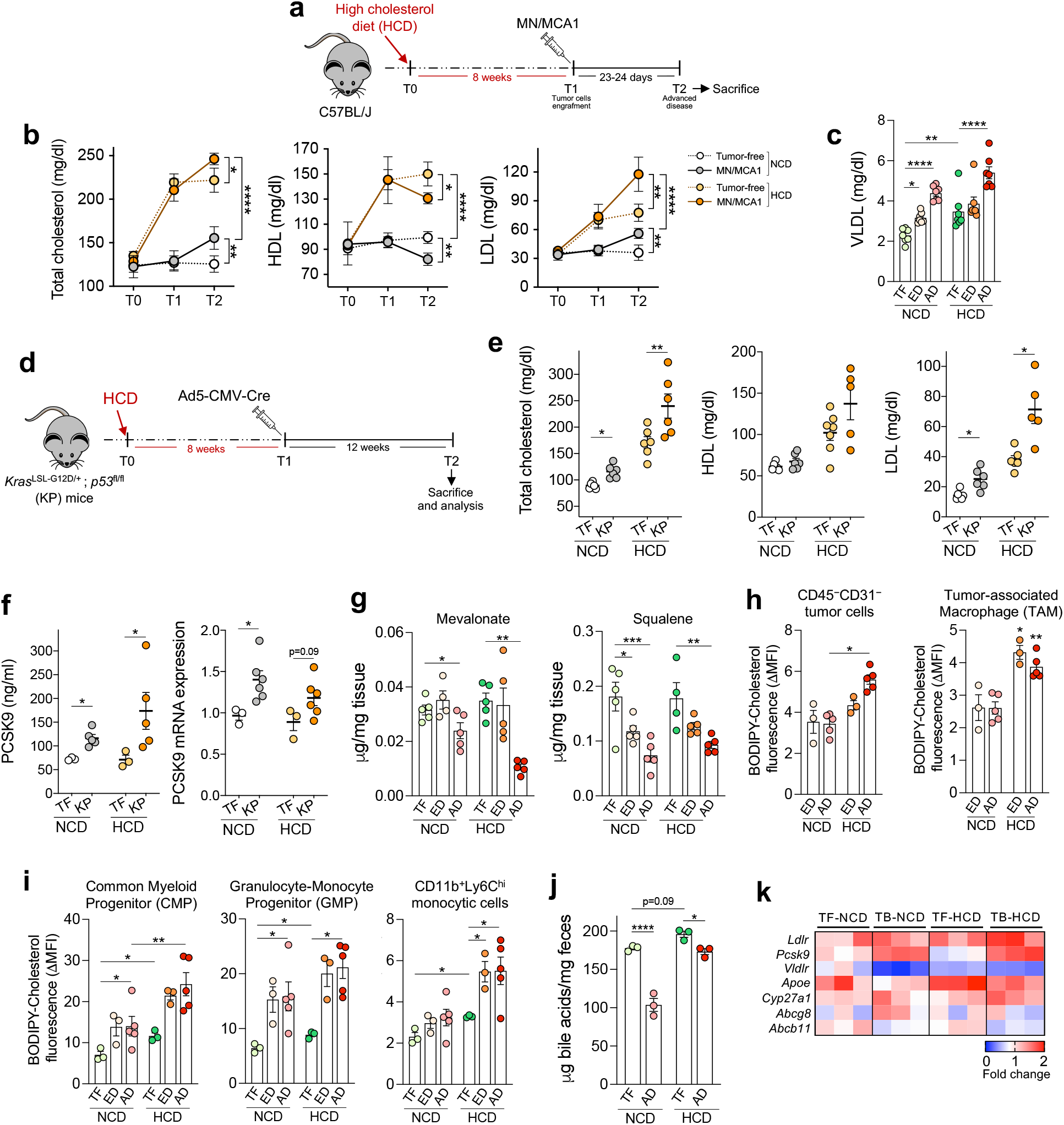
Hypercholesterolemia exacerbates alterations in cholesterol metabolism. **a,** The experimental design involved adult mice being fed with HCD for 8 weeks before being injected with MN/MCA1 tumor cells. These mice were maintained on HCD for an additional 23-24 days, after which they were sacrificed for sample collection and analysis. **b,** Blood levels of total, HDL, and LDL cholesterol in NCD- or HCD-fed MN/MCA1 mice *vs.* TF mice. Starting from time 0 (T0), mice were maintained on their respective dietary regimens for 8 weeks, injected (T1) with MN/MCA1 tumor cells, and then sacrificed at AD stage (T2) (n=9). **c**, Blood VLDL levels in NCD- or HCD-fed MN/MCA1 mice at ED or AD compared to those observed in TF mice (n=8). **d**, The experimental design for *Kras*-mutated (*Kras*^LSL-G12D/+^) and *p53*-floxed (*p53*^fl/fl^) (KP) mice at 6 weeks of age involved conditioning of these animals with HCD or NCD for 8 weeks before intranasal inoculation of an adenoviral vector expressing the Cre recombinase (Ad5-CMV-Cre) to cause sporadic mutations and promote lung tumor development. After 12 weeks, mice were sacrificed for sample collection and analysis. **e**, Blood levels of total, HDL, and LDL cholesterol in KP mice on NDC or HCD *vs.*TF (not inoculated with Ad5-CMV-Cre) mice (n=6). **f,** Circulating PCSK9 levels (n=5) and hepatic *Pcsk9* mRNA expression (n=6) in KP mice on NDC or HCD *vs.* TF mice (n=3). **g,** Mass spectrometry determination of mevalonate and squalene in livers from NCD- or HCD-fed MN/MCA1 mice at ED or AD *vs.* TF mice (n=5). **h, i**, FACS quantification of BODIPY-Cholesterol mean fluorescence intensity (ΔMFI) of intratumoral CD45^-^CD31^-^ cells and CD11b^+^Ly6C^lo/–^F4/80^+^ tumor-associated macrophages (TAMs) (**h**), and in bone marrow (BM) myeloid progenitors (both CMPs and GMPs) or blood CD11b^+^Ly6C^hi^ monocytic cells (**i**) from TF (n=3), ED (n=3), or AD (n=5) MN/MCA1 mice on NCD or HCD. **j,** Quantification of fecal total bile acids (TBA) in NCD- or HCD-fed MN/MCA1 mice (n=3). **k,** Heatmap representing differential mRNA expression levels of cholesterol metabolism genes in primary hepatocytes from healthy wt mice stimulated *in vitro* with serum from NCD- or HCD-fed MN/MCA1 tumor-bearing (TB) or tumor-free (TF) mice (n=3). Data are representative of three (**b**) or two (**e, f**) independent experiments; as for **c, g-k**, one experiment was performed. Data are presented as mean ± SEM. \**P* < 0.05, \*\**P* < 0.01, \*\*\**P* < 0.001, \*\*\*\**P* < 0.0001 between selected relevant comparisons. **b, k,** Two-way ANOVA with Tukey’s multiple comparisons test. **c, e-j,** One-way ANOVA with Tukey’s multiple comparisons test.

**Extended Data Figure 3.**
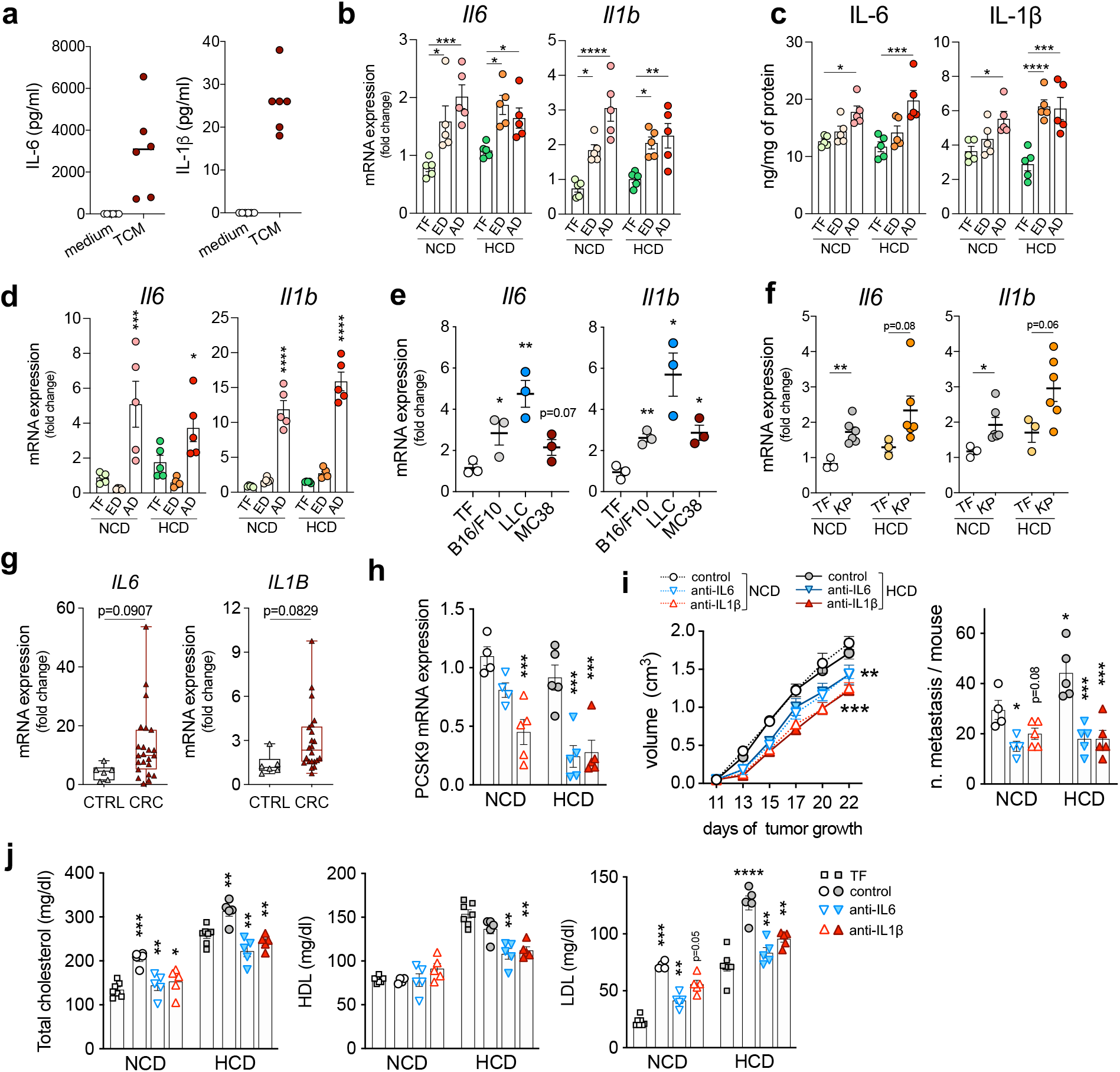
IL-6 and IL-1β modulate PCSK9 expression and cholesterolemia in MN/MCA1-bearing mice. **a,** IL-6 and IL-β protein quantification in MN/MCA1 tumor-conditioned medium (TCM) *vs.* unconditioned medium (n=6). **b-d**, IL-6 and IL-1β quantification in NDC- or HCD-fed MN/MCA1 mice at ED or AD *vs.* TF mice (n=5): hepatic mRNA expression (**b**); hepatic protein quantification (**c**); adipose mRNA expression (**d**). **e, f**, IL-6 and IL-1β mRNA expression in livers from B16-F10, LLC, and MC38 mice on NCD (n=3) (**e**) or KP mice on NCD or HCD (n=6) (**f**) compared to their respective TF controls (n=3). **g,** IL-6 and IL-1β mRNA expression in healthy liver parenchyma of CRC patients (n=23) compared to healthy liver parenchyma from patients bearing benign lesions (n=6). **h, i**, Hepatic PCSK9 mRNA expression (**h**), primary tumor growth (left panel) and lung metastasis count (right panel) (**i**) in NCD- or HCD-fed MN/MCA1 mice treated with anti-IL-6 or anti-IL-1β agent compared to untreated control mice similarly fed. **j**, Total, HDL, and LDL blood cholesterol levels in NCD- or HCD-fed MN/MCA1 mice treated with anti-IL-6 or anti-IL-1β agent compared to untreated control or TF mice similarly fed. **a-h, j,** One experiment was performed; **i**, Data are representative of two independent experiments. **a-f, h-j,** Data are presented as mean ± SEM; **g**, Box-and-whisker min-to-max plots. \**P* < 0.05, \*\**P* < 0.01, \*\*\**P* < 0.001, \*\*\*\**P* < 0.0001 between selected relevant comparisons. **b-f, h, i** (right)**, j,** One-way ANOVA with Tukey’s multiple comparisons test. **g**, Unpaired two-tailed *t*-test. **i** (left), Two-way ANOVA with Tukey’s multiple comparisons test.

**Extended Data Figure 4.**
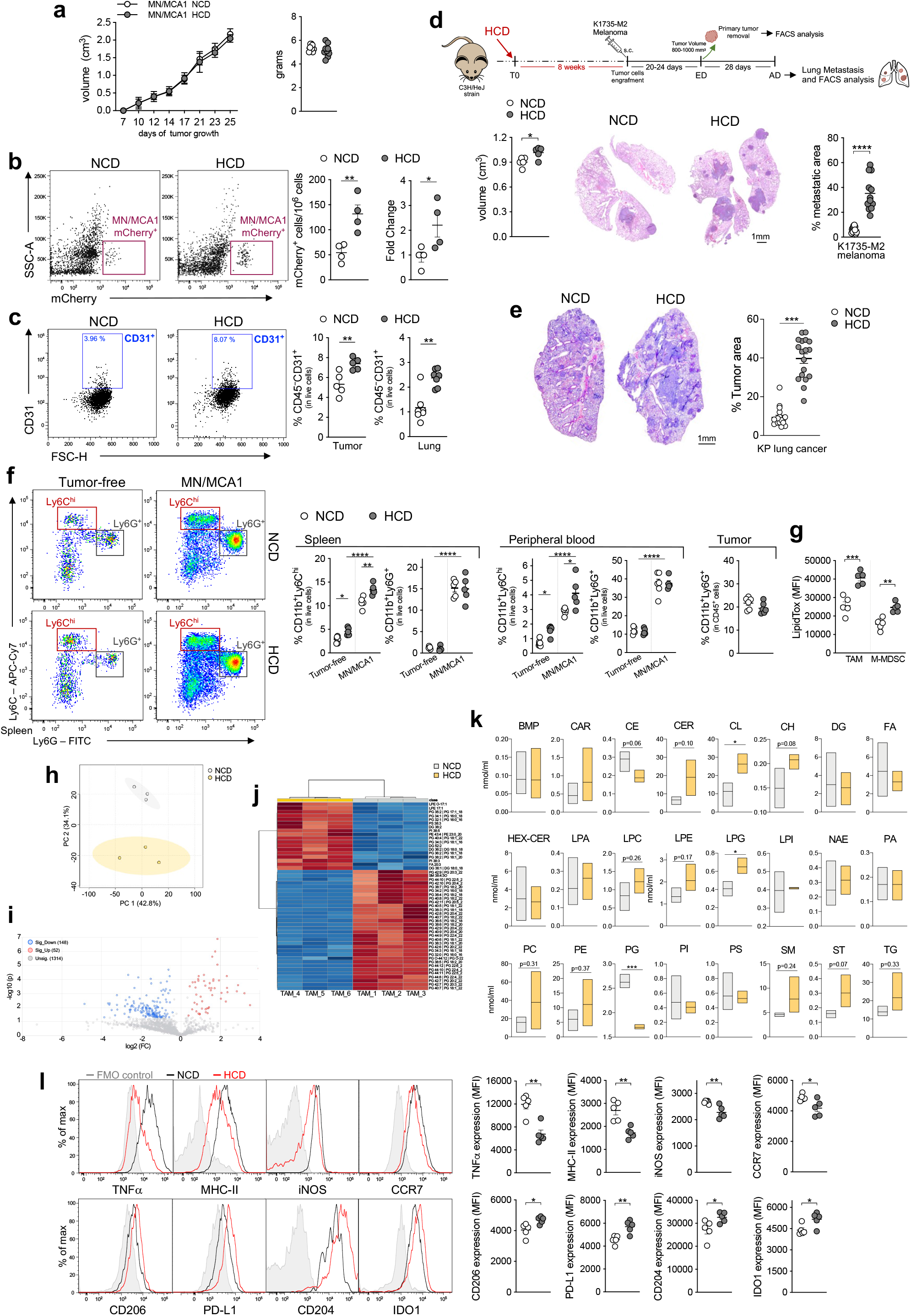
Hypercholesterolemia exacerbates cancer progression and shapes the immunosuppressive myeloid compartment. **a-c,**MN/MCA1 mice fed with NCD or HCD. **a,** Primary tumor growth, volume, and weight (n=9). **b**, Representative FACS plots and quantification of blood circulating mCherry^+^ MN/MCA1 tumor cells and mCherry mRNA expression (RT-PCR) (n=4). **c**, Quantification of CD45^-^CD31^+^ endothelial cells (FACS) in tumors and lungs (n=5). **d** (top), Experimental design: C3H/HeJ adult mice were kept on HCD for 8 weeks before being injected with K1735-M2 tumor cells. The mice were maintained on their respective dietary regimens for an additional 23-24 days before tumor explantation (volume ∼ 0.80-1.00 cm^3^). **d** (bottom), Primary tumor volume (left) and lung metastatic area (right) (n=4). Representative images are shown. Scale bar, 1 mm. **e**, Primary tumor area in KP mice under NCD or HCD (n=6). Representative images are shown. Scale bar, 1 mm. **f, g, k,** MN/MCA1 or tumor-free mice fed with NCD or HCD. **f** (left), Representative FACS plots of monocytic and granulocytic cells in the spleens of both tumor-free and MN/MCA1-bearing mice. **f** (right), Frequencies of monocytic (CD11b^+^Ly6G^-^Ly6C^hi^) and granulocytic (CD11b^+^Ly6G^+^Ly6C^lo^) cells in the spleens (n=5), peripheral blood (n=5) and tumors (n=6). **g,** Mean fluorescence intensity (MFI) of LipidTOX in TAMs and M-MDSCs (n=5). **h-k,** Lipidomic analysis of TAMs from NCD- or HCD-fed mice (n=3). **h,** Principal component (PC) analysis of lipidomic data; **i,** Volcano plot depicting the modulation of 200 lipids (FC > 1.3 and p-value < 0.05); **j,** Hierarchical heat maps of the abundances of quantified lipids; **k,** Graphs representing the main lipid species or classes analyzed: bis(monoacyl)glycerophosphate (BMP), acylcarnitine (CAR), cholesteryl esters (CE), ceramides (CER), cholesterol (CH), cardiolipin (CL), diacylglycerol (DG), free fatty acids (FA), hexosylceramides (HEX-CER), lysophosphatidic acids (LA), lysophophatidylcholines (LC), lysophosphatidylethanolamines (LPE), lysophosphatidylglycerols (LPG), lysophosphatidylinositols (LPI), N-acyl ethanolamines (NAE), phosphatidic acids (PA), phophatidylcholines (PC), phosphatidylethanolamines (PE), phosphatidylglycerol (PG), phosphatidylinositols (PI), phosphatidylserines (PS), sphingomyelins (SM), sterols (ST), and triacylglycerols (TG). **l,** FACS analysis of M1 (top: TNFα, MHC-II, iNOS, and CCR7) and M2 (bottom: CD206, PD-L1, CD204, and IDO1) polarization markers of TAMs from NCD- or HCD-fed MN/MCA1 mice (n=5). Data are representative of at least five (**a**), three (**f, g**), or two (**e, l**) independent experiments. **b-d, h-k,** One experiment was performed. **a-g, l,** Data are presented as mean ± SEM. **k**, Box-and-whisker min-to-max plots. \**P* < 0.05, \*\**P* < 0.01, \*\*\**P* < 0.001, \*\*\*\**P* < 0.0001 between selected relevant comparisons. **a,** Two-way ANOVA with Tukey’s multiple comparisons test; **b-e, g, k, l**, Unpaired two-tailed *t*-test. **f**, One-way ANOVA with Tukey’s multiple comparisons test.

**Extended Data Figure 5.**
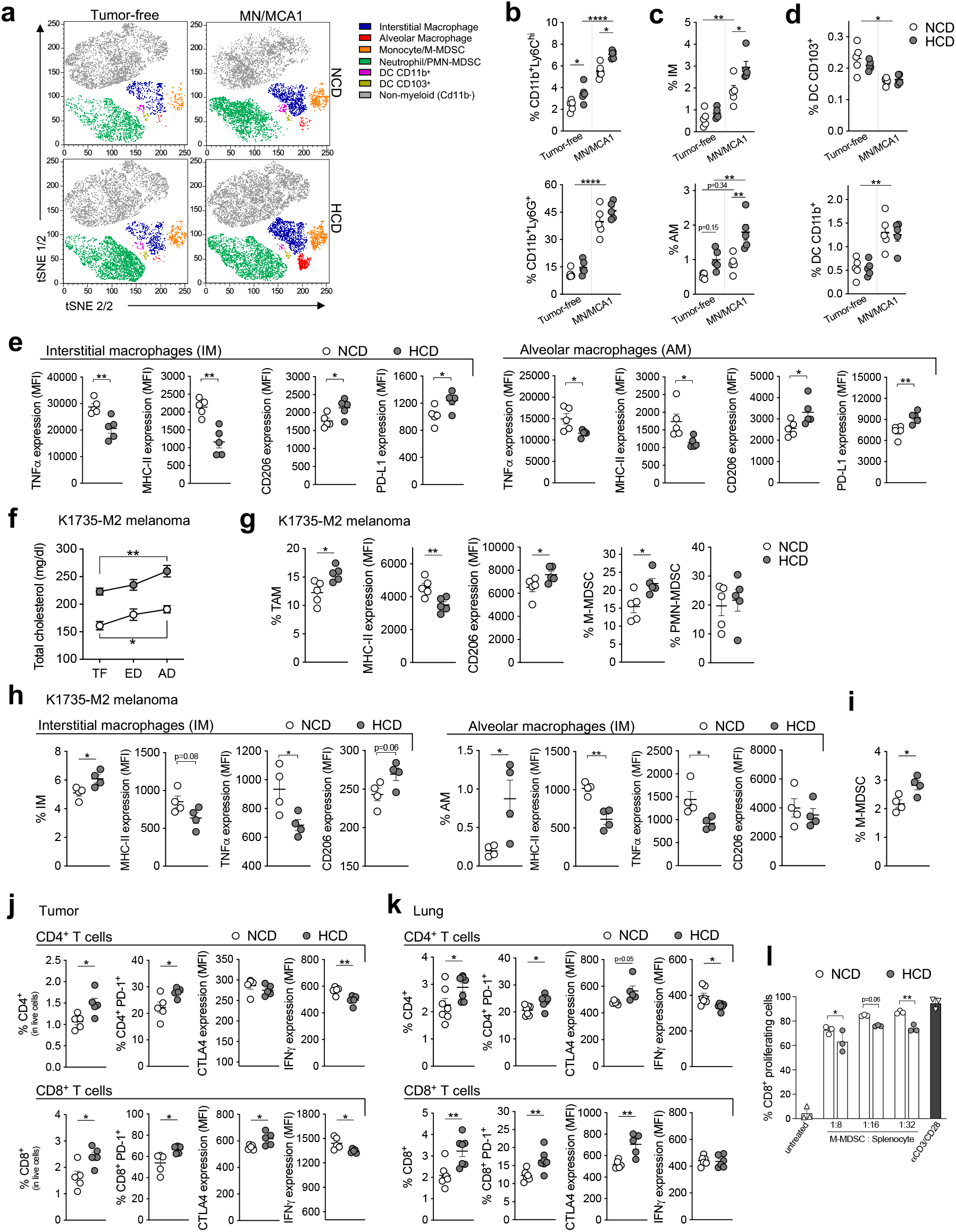
Hypercholesterolemia shapes immunosuppressive myeloid cells in metastatic tissue from both MN/MCA1- and K1735-M2-bearing mice. **a,** t-distributed stochastic neighborhood embedding (t-SNE) of CD45^+^ cells and myeloid subsets in the lungs. **b-d,** Frequencies of lung monocytic and granulocytic cells (**b**), interstitial (IM), and alveolar macrophages (AM) (**c**), and CD103^+^ and CD11b^+^ dendritic cell subsets (**d**) (n=5). **e,** Expression levels of M1 (TNFα, MHC-II) and M2 (CD206, PD-L1) polarization markers in IMs (left) and AMs (right). **f,** Total blood cholesterol levels in NCD- or HCD-fed mice prior to injection of K1735-M2 tumor cells (TF), at either the time of primary tumor removal (early disease, ED) or lung metastasis evaluation (advanced disease, AD) (n=6). **g,** FACS quantification of TAMs and relative MHC-II and CD206 expression levels (MFI), as well as M-MDSC and PMN-MDSC frequencies, in tumor tissues from NCD- or HCD-fed K1735-M2 mice fed (n=5). **h,** FACS quantification of IM (left) and AM (right) frequencies and relative MHC-II, TNFα and CD206 expression (MFI). **i,** FACS quantification of M-MDSCs in metastatic lungs from NCD- or HCD-fed K1735-M2 mice (n=4). **j, k,** Frequencies of CD4^+^ (top) and CD8^+^ (bottom) T cells and relative PD-1^+^ subsets and expression levels of CTLA4 and IFNγ in tumors (n=5) (**j**) and lungs (n=6) (**k**). **l,** Proliferation of CFSE-labeled CD8^+^ T cells activated with anti-CD3 and anti-CD28, alone or in coculture with FACS-sorted M-MDSCs from NCD- or HCD-fed MN/MCA1 mice (n=3) at different ratios. Data are representative of at least three (**b-e**), or two (**j-l**) independent experiments; **f-i,** one experiment was performed. Data are presented as mean ± SEM. \**P* < 0.05, \*\**P* < 0.01, \*\*\**P* < 0.001, \*\*\*\**P* < 0.0001 between selected relevant comparisons. **b-e,** One-way ANOVA with Tukey’s multiple comparisons test; **f-k,** Unpaired two-tailed *t*-test; **l,** One-way ANOVA with Šidák’s multiple comparisons test.

**Extended Data Fig. 6.**
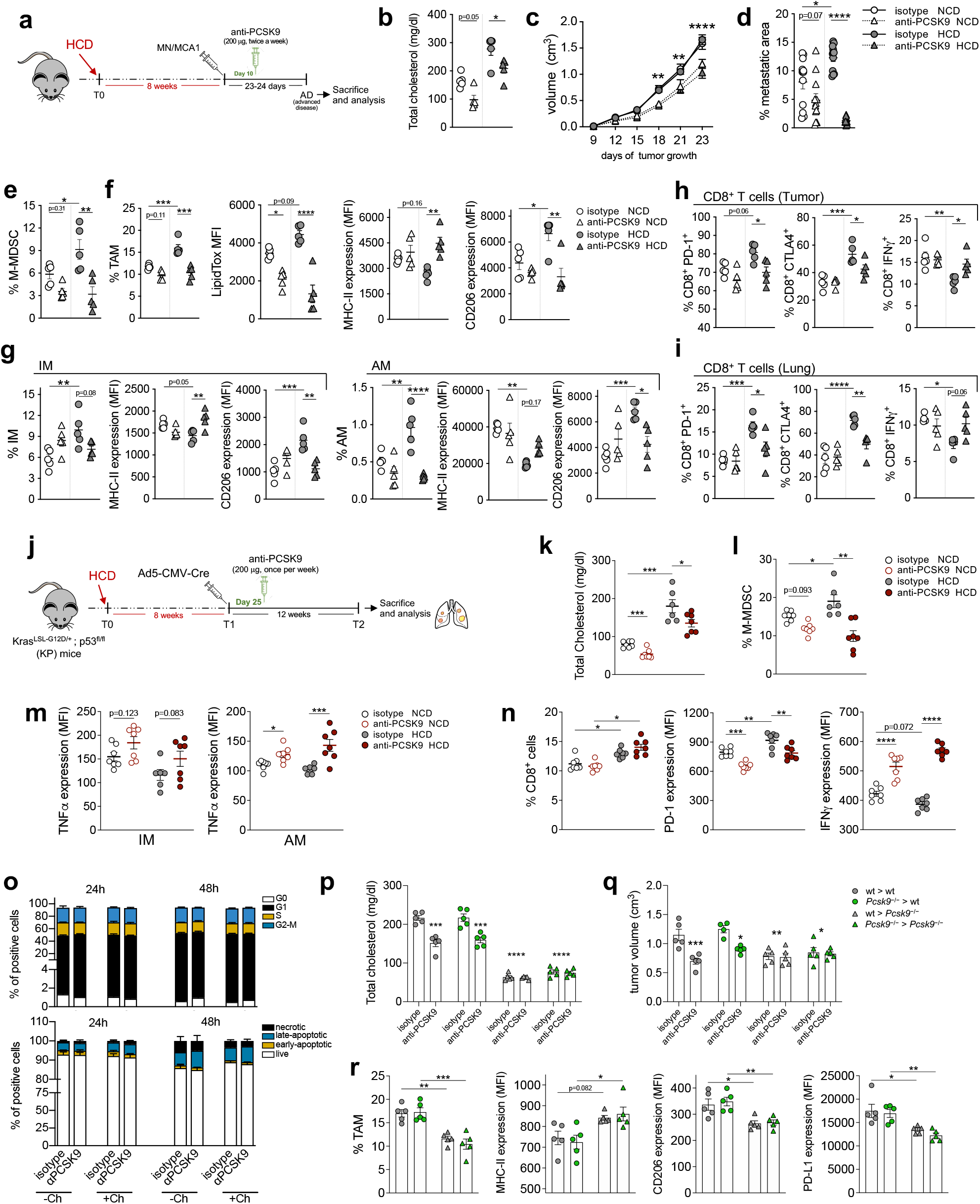
Lowering cholesterol in cancer bearers mitigates immunosuppressive myelopoiesis. **a,** Experimental design: wt mice were fed HCD for 8 weeks before being injected with MN/MCA1 tumor cells. When tumors were palpable, an anti-PCSK9 blocking antibody was administered. After 23 days of tumor growth, mice were sacrificed for samples collection and analysis. **b-i,** NCD- or HCD-fed MN/MCA1-bearing wt mice treated with anti-PCSK9 or isotype control antibody (n=5): **b,** total blood cholesterol levels; **c,** tumor volumes; **d,** lung metastatic areas; **e**, frequencies of intratumoral M-MDSCs; **f**, FACS quantification of TAM frequencies and their relative LipidTOX, MHC-II, and CD206 expression levels (MFI); **g,** frequencies of pulmonary IMs (left) and AMs (right), and relative MHC-II and CD206 expression levels (MFI); **h, i,** Frequency of PD-1^+^, CTLA4^+^, and IFNγ^+^ in CD8 T cells from tumors (**h**) or metastatic lungs (**i**). **j,** Experimental design: KP mice at 6 weeks of age were conditioned with HCD or NCD for 8 weeks before intranasal inoculation of Ad5-CMV-Cre. After further 25 days, an anti-PCSK9 blocking antibody was administered once a week. After 12 weeks, mice were sacrificed for samples collection and analysis. **k-n,** NCD- or HCD-fed KP mice treated with anti-PCSK9 or isotype control antibody (n=6): **k,** total blood cholesterol levels; **l,** blood M-MDSC frequency; **m,** TNFα expression (MFI) in IM (left) and AM (right) cells; **n,** frequency of pulmonary CD8^+^ T cells and relative PD-1 and IFNγ expression levels (MFI). **o,** FACS evaluation of cell cycle (top) and viability (bottom) of MN/MCA1 tumor cells treated *in vitro* with anti-PCSK9 antibody or isotype control, supplemented or not with cholesterol (Ch), for 24 or 48 h (n=3). **p-r,** Wt or *Pcsk9*^-/-^ mice on HCD, transplanted with either wt (wt>wt; wt>*Pcsk9*^-/-^) or *Pcsk9*^-/-^ (*Pcsk9*^-/-^>wt; *Pcsk9*^-/-^>*Pcsk9*^-/-^) BM cells, engrafted with MN/MCA1 cells and treated with anti-PCSK9 antibody or isotype control: **p,** total blood cholesterol levels; **q,** primary tumor volume. **r**, FACS analysis of TAM frequency (left) and relative MHC-II, CD206, and PD-L1 expression levels (left to right) (MFI) in isotype-treated mice (n=5). **b-i, n, p-r,** One experiment was performed; **k-m, o,** data are representative of two independent experiments. Data are presented as mean ± SEM. \**P* < 0.05, \*\**P* < 0.01, \*\*\**P* < 0.001, \*\*\*\**P* < 0.0001 between selected relevant comparisons. **b, d-o, r,** One-way ANOVA with Tukey’s multiple comparisons test; **c, p, q,** Two-way ANOVA with Tukey’s multiple comparisons test.

**Extended Data Figure 7.**
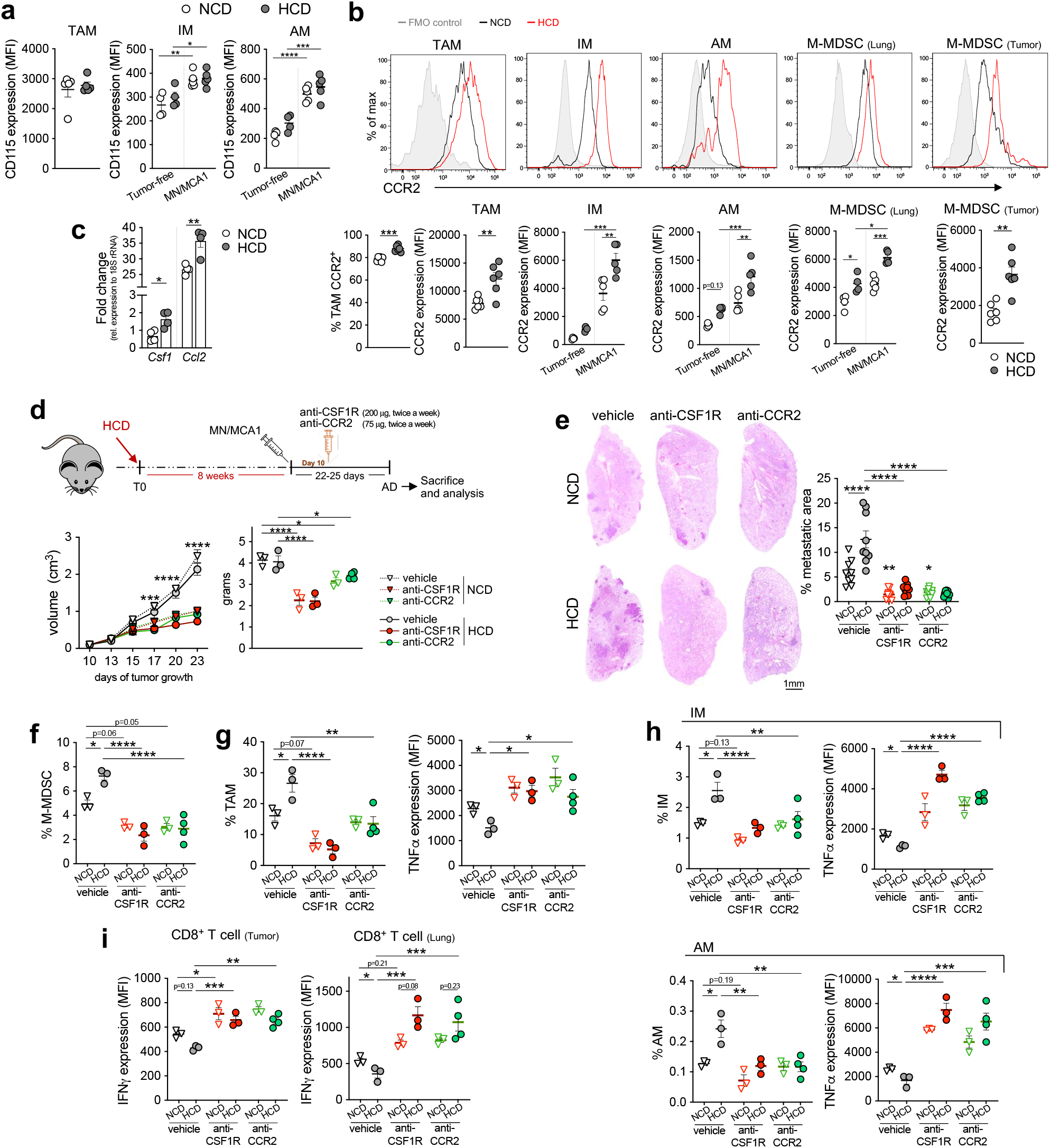
CSF-1/CSF-1R and CCL2/CCR2 axes in hypercholesterolemia-induced myelopoiesis. **a,** FACS quantification of CD115 expression (MFI) in TAMs, IMs, and AMs from NCD- or HCD-fed MN/MCA1-bearing (n=5) or tumor-free (n=4) mice. **b,** Flow cytometry histograms (top) and quantification (bottom) of CCR2 expression (MFI) in TAMs, IMs, AMs, and M-MDSCs from NCD- or HCD-fed MN/MCA1-bearing (n=5), or tumor-free (n=4) mice. **c,** mRNA expression of *Csf1* and *Ccl2* genes in MN/MCA1 tumors from NCD- or HCD-fed mice (n=4). **d-h,** NCD or HCD-fed MN/MCA1 mice were treated with anti-CSF1R, anti-CCR2, or vehicle control (vehicle, anti-CSF1R in NCD or HCD, n=3; anti-CCR2 in NCD, n=3; anti-CCR2 in HCD, n=4). **d** (top), Experimental design: adult mice were kept in HCD for 8 weeks prior to injection of MN/MCA1 tumor cells. When tumors were palpable, an anti-CSF1R or anti-CCR2 antibody was administered. Mice were sacrificed after 24 days of tumor growth. **d** (bottom), Primary tumor growth expressed as volume and weight; **e**, Lung metastatic area; **f**, intratumoral M-MDSCs; **g**, intratumoral TAM frequency and relative TNFα expression (MFI); **h**, pulmonary IM (top) and AM (bottom) frequencies and respective TNFα expression; **i,** IFNγ expression (MFI) in CD8^+^ T cells of tumors and metastatic lungs. **a, b**, Data are representative of at least three independent experiments; **c-i,** one experiment was performed. Data are presented as mean ± SEM. \**P* < 0.05, \*\**P* < 0.01, \*\*\**P* < 0.001, \*\*\*\**P* < 0.0001 between selected relevant comparisons. **a, b, d** (right)**, e-i,** One-way ANOVA with Tukey’s multiple comparisons test; **c**, Unpaired two-tailed *t*-test; **d** (left), Two-way ANOVA with Tukey’s multiple comparisons test.

**Extended Data Figure 8.**
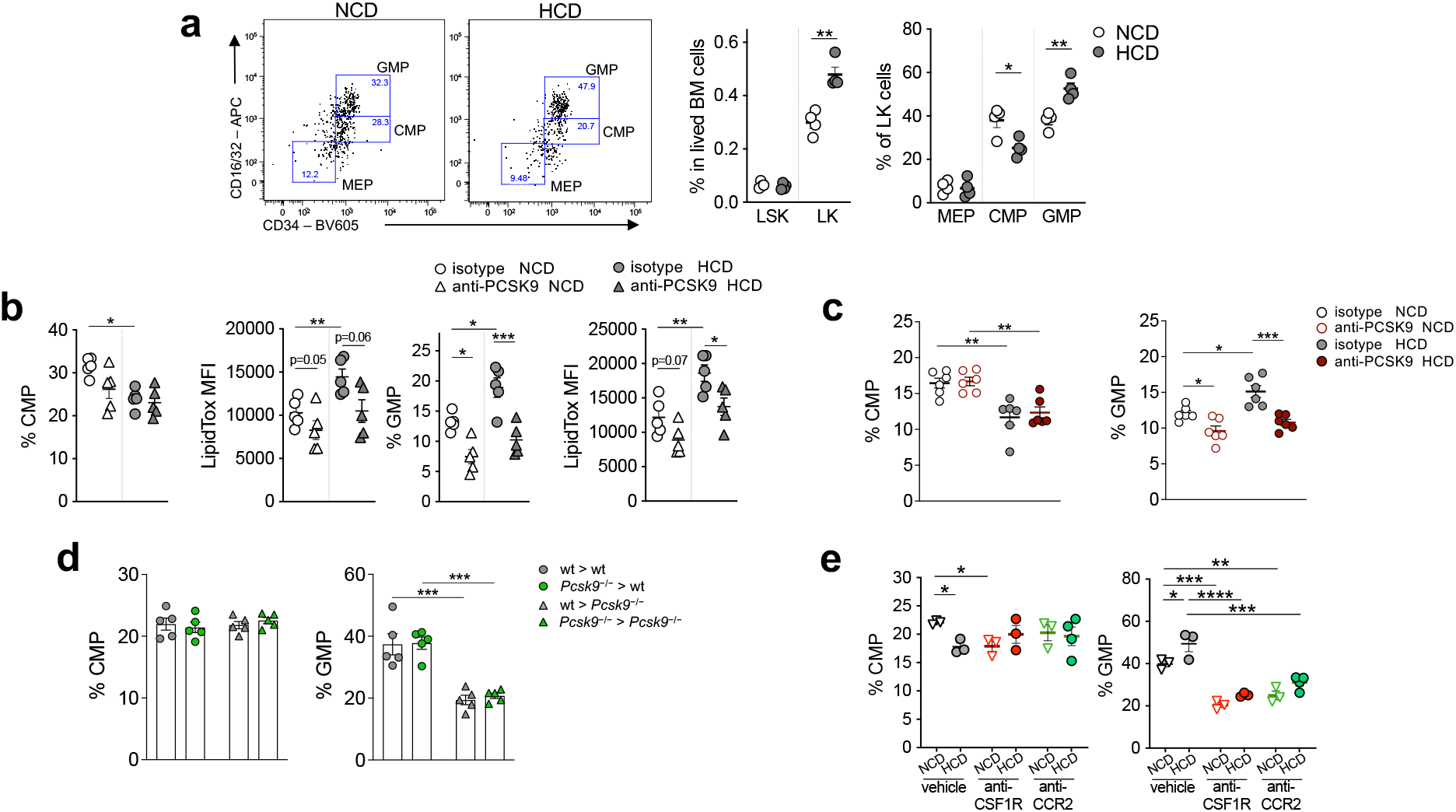
Hypercholesterolemia boosts CMPs to GMPs transition. **a,** FACS quantification of LSK, LK, MEPs, CMPs, and GMPs in BM of NCD- or HCD-fed MN/MCA1 mice (n=4). **b,** Frequency of CMPs and GMPs with relative LipidTOX quantification (MFI) in BM of NCD- or HCD-fed MN/MCA1 mice treated with anti-PCSK9 or isotype control antibody (n=5). **c,** FACS quantification of CMPs and GMPs in BM of NCD- or HCD-fed KP mice treated with anti-PCSK9 or isotype control antibody (n=6). **d,** Frequency of CMPs and GMPs in BM of HCD-fed MN/MCA1 wt or *Pcsk9*^-/-^ mice transplanted with either wt (wt>wt; wt>*Pcsk9*^-/-^) or *Pcsk9*^-/-^ (*Pcsk9*^-/-^>wt; *Pcsk9*^-/-^>*Pcsk9*^-/-^) BM cells (n=5). **e,** Frequency of CMPs and GMPs in BM from NCD- or HCD-fed MN/MCA1 mice treated with anti-CSF1R, anti-CCR2, or vehicle control (vehicle, anti-CSF1R in NCD or HCD, n=3; anti-CCR2 in NCD, n=3; anti-CCR2 in HCD, n=4). Data are representative of at least three (**a**), or two (**c**) independent experiments. **b, d, e,** one experiment was performed. Data are expressed as mean ± SEM. \**P* < 0.05, \*\**P* < 0.01, \*\*\**P* < 0.001, \*\*\*\**P* < 0.0001 between selected relevant comparisons. **a,** Unpaired two-tailed *t*-test. **b-e,** One-way ANOVA with Tukey’s multiple comparisons test.

**Extended Data Fig. 9.**
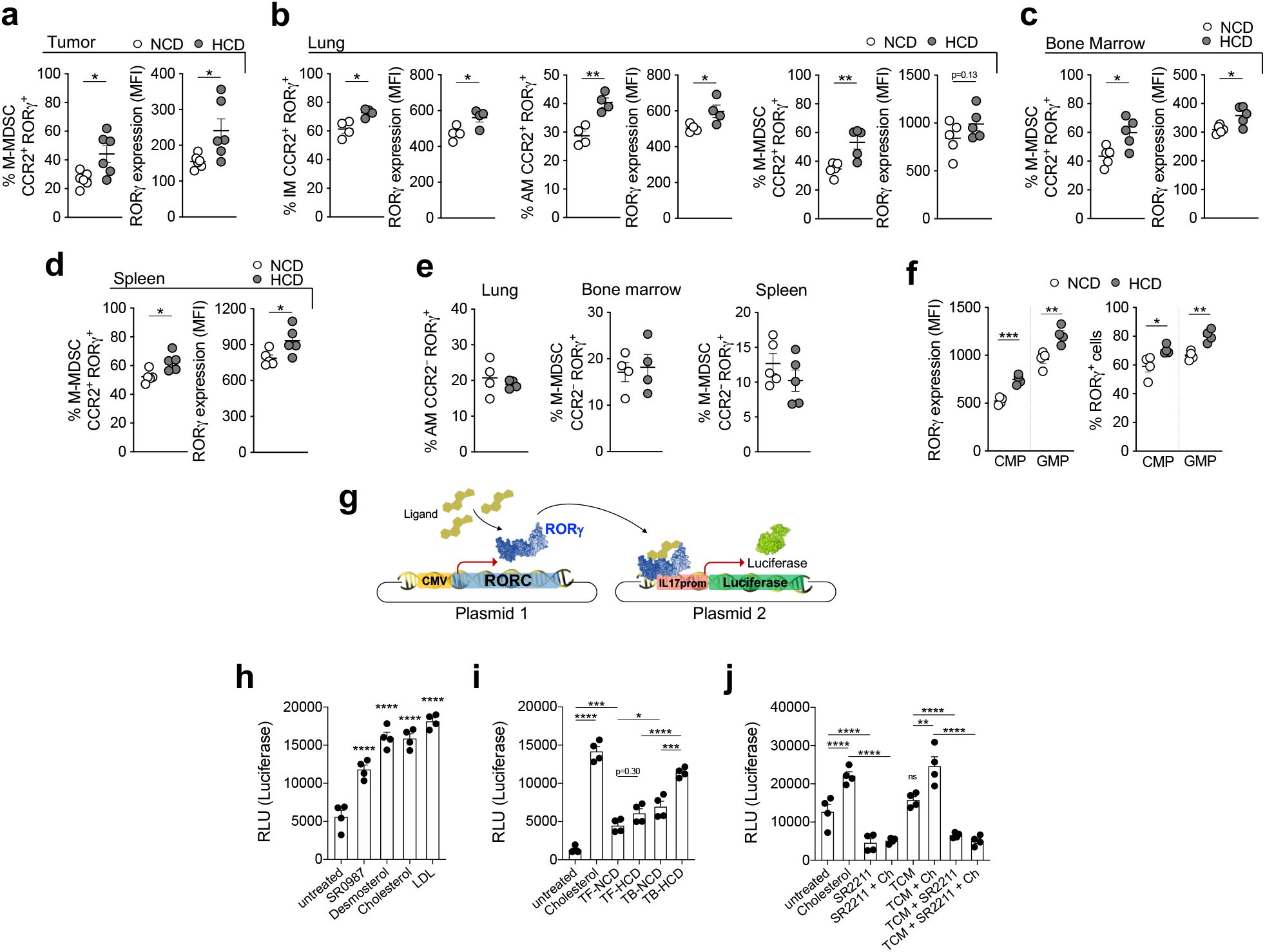
Cholesterol enhances myeloid expression of RORγ and its transcriptional activity. MN/MCA1 mice were fed NCD or HCD. **a-d,** Frequency of RORγ^+^CCR2^+^ cells and relative RORγ expression levels in: **a,** M-MDSCs from tumors (n=6); **b,** lung IMs (n=4), AMs (n=4), and M-MDSCs (n=5); **c,** BM M-MDSCs (n=5); **d,** Spleen M-MDSCs (n=5). **e,** Frequency of RORγ^+^CCR2^-^ cells and relative RORγ expression levels in lung AMs (n=4), BM or spleen M-MDSCs (n=5). **f,** Frequency of RORγ^+^ CMPs and GMPs cells and relative RORγ expression levels (n=4). **g,** HEK293 cells were co-transfected with CMV-RORγ and IL17A_prom-Luc plasmids. Plasmid 1: RORγ gene is constitutively expressed under the control of CMV promoter. Plasmid 2: Luciferase reporter gene is under the control of the minimal promoter of *Il17a* gene, whose transcription is driven by RORγ. **h-j,** HEK293 cells, co-transfected with plasmids 1 and 2, were stimulated with SR0987, desmosterol, cholesterol, or LDL (**h**), or with sera from NCD- or HCD-fed MN/MCA1 TB or TF mice (n=4) (**i**), or exposed to single or combinations of cholesterol, SR2211 and TCM (**j**). Luciferase activity was quantified as relative light units (RLU). Data are representative of five (**a, b, d**), two (**c, e, h-j**), or three (**f**) independent experiments. Data are expressed as mean ± SEM. \**P* < 0.05, \*\**P* < 0.01, \*\*\**P* < 0.001, \*\*\*\**P* < 0.0001 between selected relevant comparisons. **a-f,** Unpaired two-tailed *t*-test; **h-j,** One-way ANOVA with Tukey’s multiple comparisons test.

**Extended Data Fig. 10.**
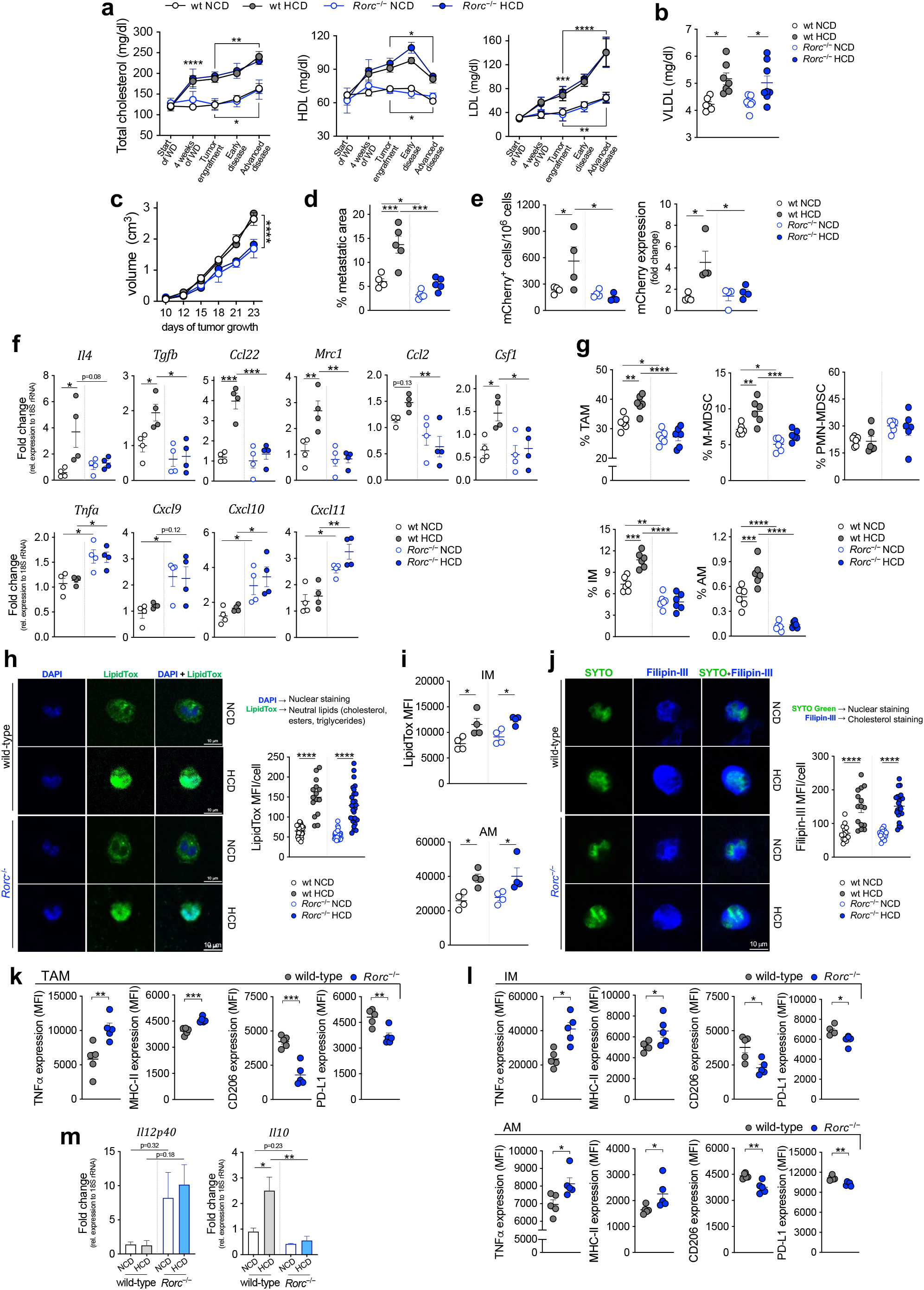
RORγ couples hypercholesterolemia to protumoral myelopoiesis. **a-j,**Wt or *Rorc*^-/-^ MN/MCA1 mice fed NCD or HCD. **a,** Total, HDL, and LDL blood cholesterol levels (n=7); **b,** blood VLDL levels at AD (n=7); **c,** tumor growth (volume) (n=5); **d,** lung metastatic area (n=5); **e,** quantification of blood circulating mCherry^+^ MN/MCA1 tumor cells (FACS) and mCherry mRNA expression (RT-PCR) (n=4); **f,** intratumoral mRNA expression of M2 and M1 genes (n=4); **g,** FACS quantification of intratumoral TAMs, M-MDSCs, PMN-MDSCs, and pulmonary IMs and AMs; **h,** immunofluorescence analysis of LipidTOX neutral lipid staining of FACS-sorted TAMs. Images are representative of each experimental group, where at least 15 cells were randomly counted. Scale bar, 10 μm; **i,** FACS quantification of LipidTOX mean fluorescence intensity (MFI) in IM and AM subsets (n=4); **j,** immunofluorescence analysis of Filipin-III cholesterol staining in FACS-sorted TAMs. Images are representative of each experimental group, where at least 15 cells were randomly counted. Scale bar: 10 μm. **k, l,** FACS quantification of M1 (TNFα, MHC-II) and M2 (CD206, PD- L1) polarization markers in TAMs (**k**), pulmonary IM (**l**, top) and AM (**l**, bottom) subsets, from HCD- fed wt and *Rorc^-/-^*mice (n=5). **m,** mRNA expression of M1-like *Il12p40* and M2-like *Il10* genes in FACS-sorted TAMs (n=2). Data are representative of five (**c, d**), two (**a, e, i**), or at least two (**g, k, l**) independent experiments; **b, f, h, j, m.** One experiment was performed. Data are expressed as mean ± SEM. \**P* < 0.05, \*\**P* < 0.01, \*\*\**P* < 0.001, \*\*\*\**P* < 0.0001 between selected relevant comparisons. **a, c,** Two-way ANOVA with Tukey’s multiple comparisons test; **b, d-i, m**, One-way ANOVA with Tukey’s multiple comparisons test; **k, l**, Unpaired two-tailed *t*-test.

**Extended Data Figure 11.**
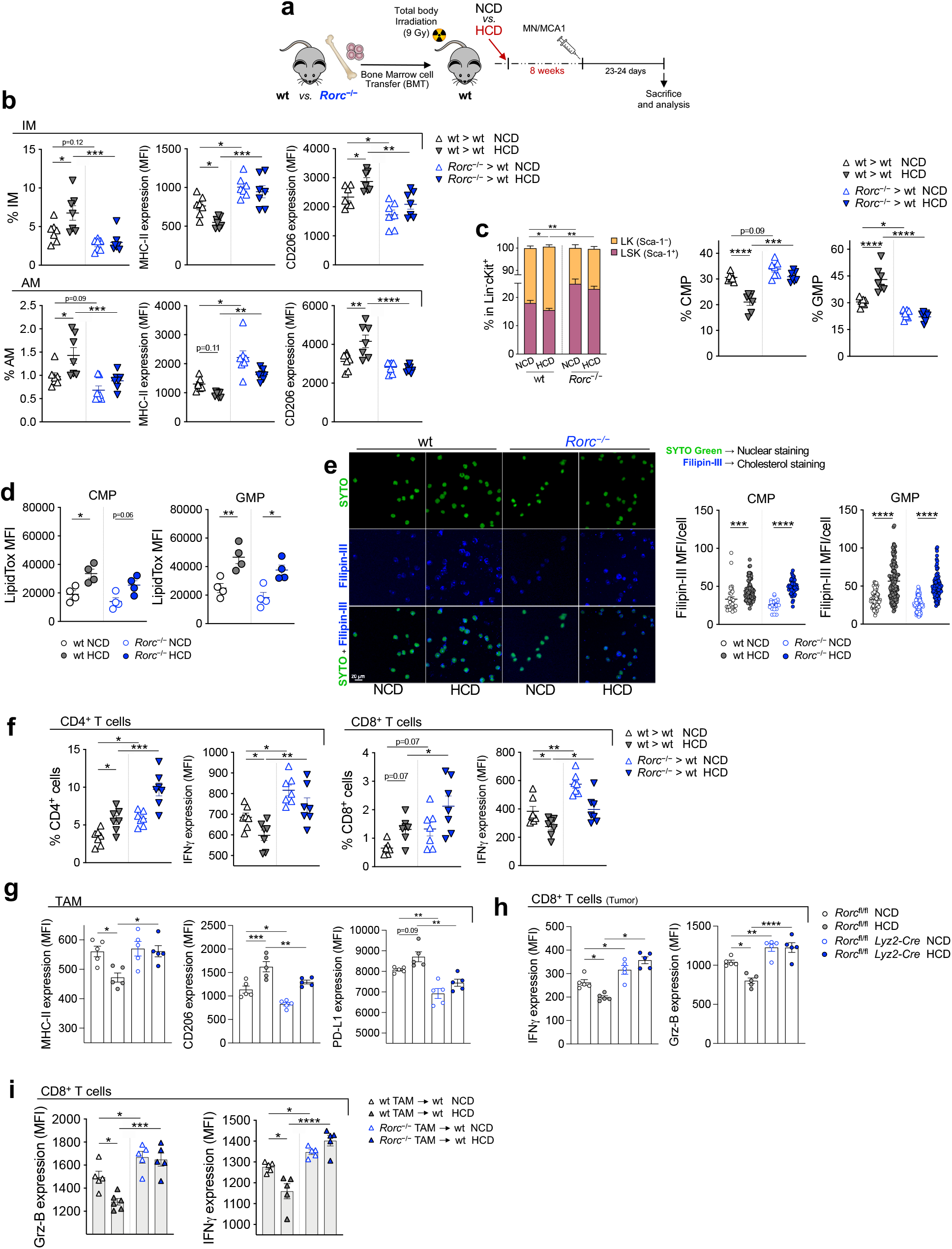
RORγ deficiency limits the effects of cholesterol on myeloid cell maturation. **a,** Experimental design: wt mice were lethally irradiated (9 gy) and transplanted with wt or *Rorc*^-/-^ BM cells. After 4 weeks, required for complete hematopoietic reconstitution, mice were conditioned for an additional 8 weeks with HCD or NCD, and then engrafted with MN/MCA1 cells. After 25 days of tumor growth, mice were sacrificed for sample collection and analysis. **b, c,** NCD- or HCD-fed MN/MCA1 wt mice transplanted with either wt (wt>wt) or *Rorc*^-/-^ (*Rorc*^-/-^>wt) BM cells (n=7). **b**, FACS quantification of IMs (top) and AMs (bottom) and their relative MHC-II and CD206 expression levels; **c**, FACS quantification of LK (Sca-1^-^) and LSK (Sca-1^+^) BM cells, CMP, and GMP frequency. **d**, FACS quantification of LipidTOX MFI in CMPs and GMPs from wt or *Rorc*^-/-^ MN/MCA1 mice on NCD or HCD (n=4). **e,** Immunofluorescence analysis of Filipin-III cholesterol staining in FACS-sorted CMPs and GMPs. Images are representative of GMP subset from each group. At least 20 cells were randomly counted. Scale bar, 20 μm. **f,** FACS quantification of intratumoral CD4^+^ and CD8^+^ cells and their IFNγ expression (MFI) in NCD- or HCD-fed MN/MCA1 wt mice transplanted with either wt (wt>wt) or *Rorc*^-/-^ (*Rorc*^-/-^>wt) BM cells (n=7). **g, h,** FACS quantification in *Rorc*^fl/fl^ and *Rorc*^fl/fl^ *Lyz2-Cre* MN/MCA1 mice on NCD or HCD (n=5) of MHC-II, CD206 and PD-L1 expression (MFI) in TAMs (**g**), and IFNγ and Grz-B expression (MFI) of intratumoral CD8^+^ T cells (**h**). **i,** FACS quantification of IFNγ and Grz-B expression (MFI) of intratumoral CD8^+^ T cells from MN/MCA1 wt mice on NCD or HCD adoptively transferred with either wt or *Rorc*^-/-^ TAMs (n=5). **a-c, e-i,** one experiment was performed; **d,** data are representative of two independent experiments. Data are expressed as mean ± SEM. \**P* < 0.05, \*\**P* < 0.01, \*\*\**P* < 0.001, \*\*\*\**P* < 0.0001 between selected relevant comparisons. One-way ANOVA with Tukey’s multiple comparisons test.

**Extended Data Figure 12.**
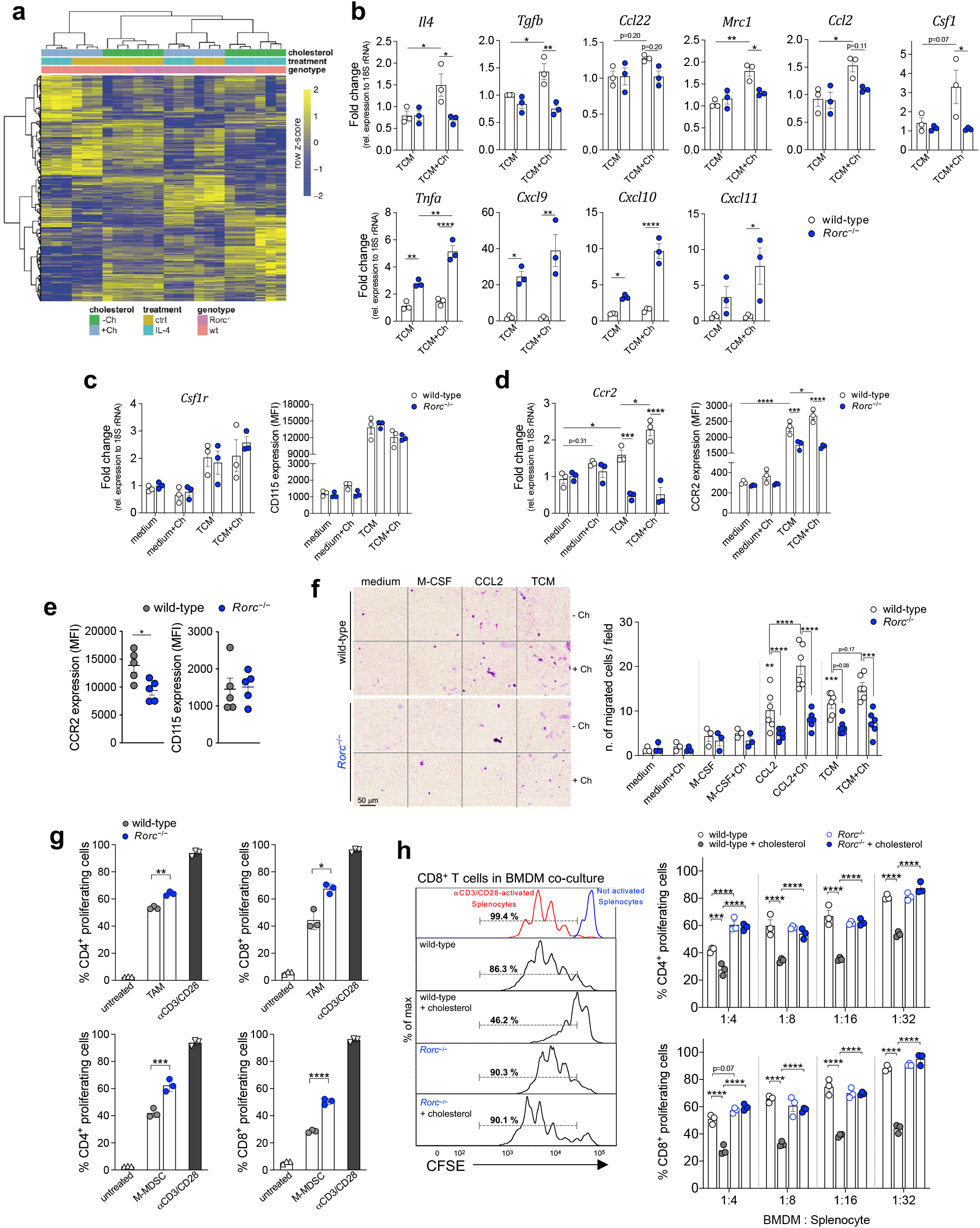
RORγ acts as a cholesterol sensor and modulator of myeloid cell immunosuppressive functions. **a,** RNA-seq analysis on wt or *Rorc*^-/-^ BMDMs differentiated with M-CSF with or without cholesterol (Ch) supplementation, and then treated or not with IL-4 (n=3). Heat map of unsupervised hierarchical clustering of 8382 differentially expressed genes (DEGs; FDR < 0.05 & |log2FC| > 0.5) found in at least one group supplemented with cholesterol (*vs.* the non-Ch- supplemented control) across four different biological conditions: wt untreated ctrl; wt IL-4; *Rorc*^-/-^ untreated ctrl, *Rorc*^-/-^ IL-4. **b,** mRNA expression of representative M2 (top) and M1 (bottom) polarization genes (n=3) in wt and *Rorc*^-/-^ peritoneal macrophages (PECs) treated with MN/MCA1- conditioned medium (TCM), supplemented or not with cholesterol (Ch). **c, d,** mRNA (left) and surface expression (MFI) (right) of *Csf1r*/CD115 (**c**) and *Ccr2*/CCR2 (**d**) in wt or *Rorc*^-/-^ PECs, treated with TCM plus Ch (n=3). **e,** FACS quantification of CCR2 and CD115 (MFI) in TAMs from HCD-fed wt or *Rorc*^-/-^ MN/MCA1 mice (n=5). **f,** Transwell migration assay of wt or *Rorc*^-/-^ PECs, pre-conditioned or not with cholesterol (Ch), in response to M-CSF (n=1), CCL2 (n=2), TCM (n=2) or medium (n=1). At least three fields per sample were randomly counted. Scale bar, 50 μm. **g, h,** Proliferation of CFSE-labeled CD4^+^ and CD8^+^ T cells activated with anti-CD3 and anti-CD28, in the presence or absence of (**g**) FACS-sorted TAMs (top) or M-MDSCs (bottom) (1:8) from either wt or *Rorc*^-/-^ HCD-fed MN/MCA1 mice (n=3), or (**h**) TCM-conditioned BMDMs from wt or *Rorc*^-/-^ mice supplemented or not with Ch. Data are representative of three (**e**) or two (**f**) independent experiments; **a-d, g-h,** One experiment was performed. Data are expressed as mean ± SEM. \**P* < 0.05, \*\**P* < 0.01, \*\*\**P* < 0.001, \*\*\*\**P* < 0.0001 between selected relevant comparisons. **a-d,** One-way ANOVA with Tukey’s multiple comparisons test; **e-g,** Unpaired two-tailed *t*-test; **f, h,** Two-way ANOVA with Tukey’s multiple comparisons test.

**Extended Data Fig. 13.**
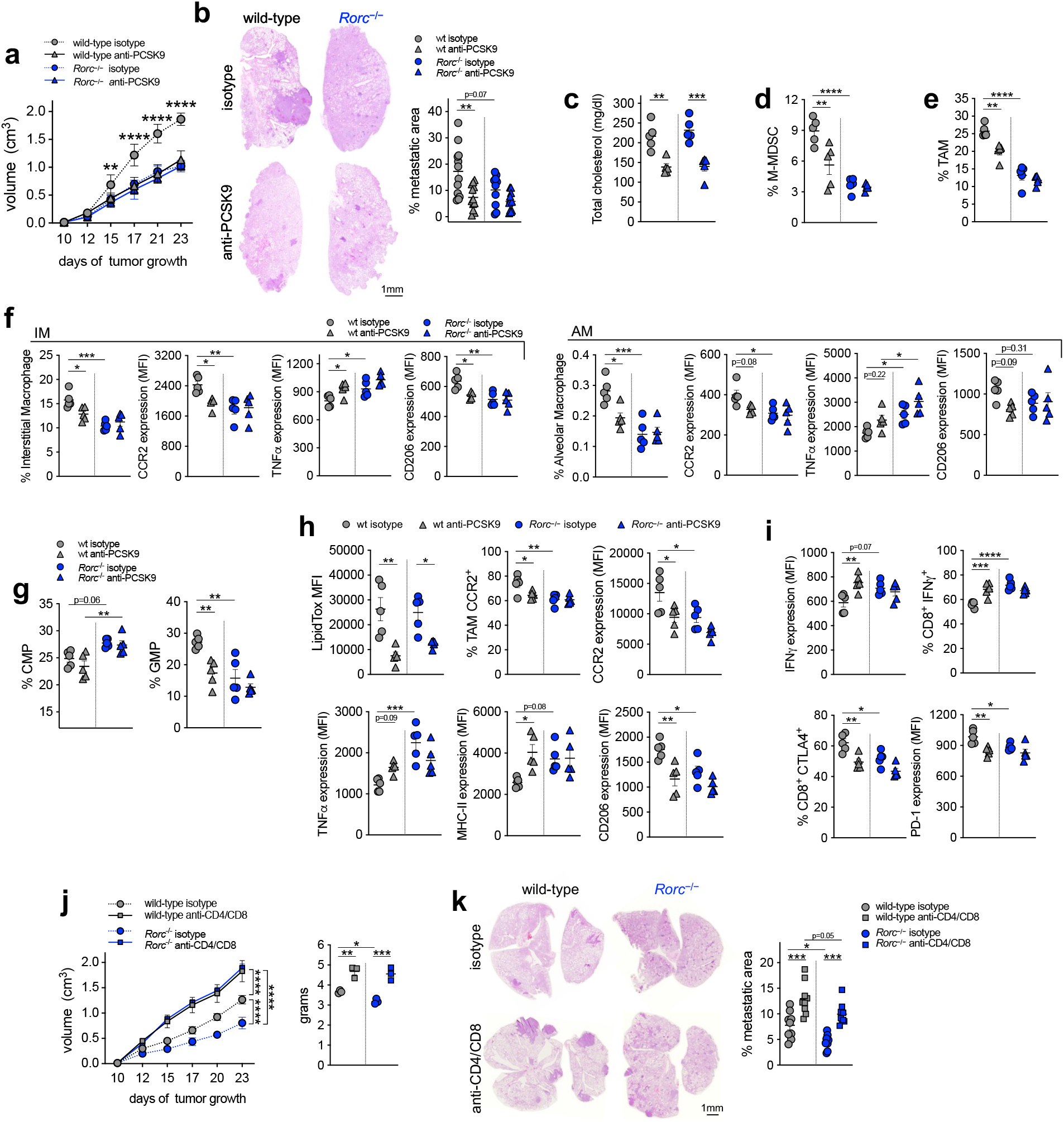
Effects of anti-PCSK9 on RORγ-dependent myelopoiesis and T cell activation. **a-i,**HCD-fed wt or *Rorc*^-/-^ MN/MCA1 mice treated with anti-PCSK9 antibody or isotype control (n=5). **a,** Tumor growth (volume); **b,** lung metastasis formation. Representative images are shown. Scale bar, 1 mm; **c,** total blood cholesterol; **d, e,** FACS quantification of intratumoral M-MDSCs (**d**) and TAMs (**e**); **f,** frequency of lung IMs (left) and AMs (right) and relative CCR2, TNFα, and CD206 expression (MFI); **g**, FACS quantification of CMPs and GMPs in BM; **h,** FACS quantification of LipidTOX mean fluorescence intensity (MFI), frequency of CCR2^+^ TAMs, and CCR2, TNFα, MHC-II, CD206 expression (MFI) in TAMs; **i,** Frequency of intratumoral CD8^+^IFNγ^+^ and CD8^+^CTLA4^+^ T cells, as well as their relative IFNγ and PD-1 expression levels (MFI). **j, k,** Tumor growth (volume and weight) (**j**) and lung metastatic area (**k**) in HCD-fed MN/MCA1 wt or *Rorc*^-/-^ mice treated with anti-CD4/anti-CD8 depleting antibodies or control isotype (n=3). One experiment was performed. Data are expressed as mean ± SEM. \**P* < 0.05, \*\**P* < 0.01, \*\*\**P* < 0.001, \*\*\*\**P* < 0.0001 between selected relevant comparisons. **a, j** (left), Two-way ANOVA with Tukey’s multiple comparisons test. **b-i, j** (right)**, k,** One-way ANOVA with Tukey’s multiple comparisons test.

**Extended Data Fig. 14.**
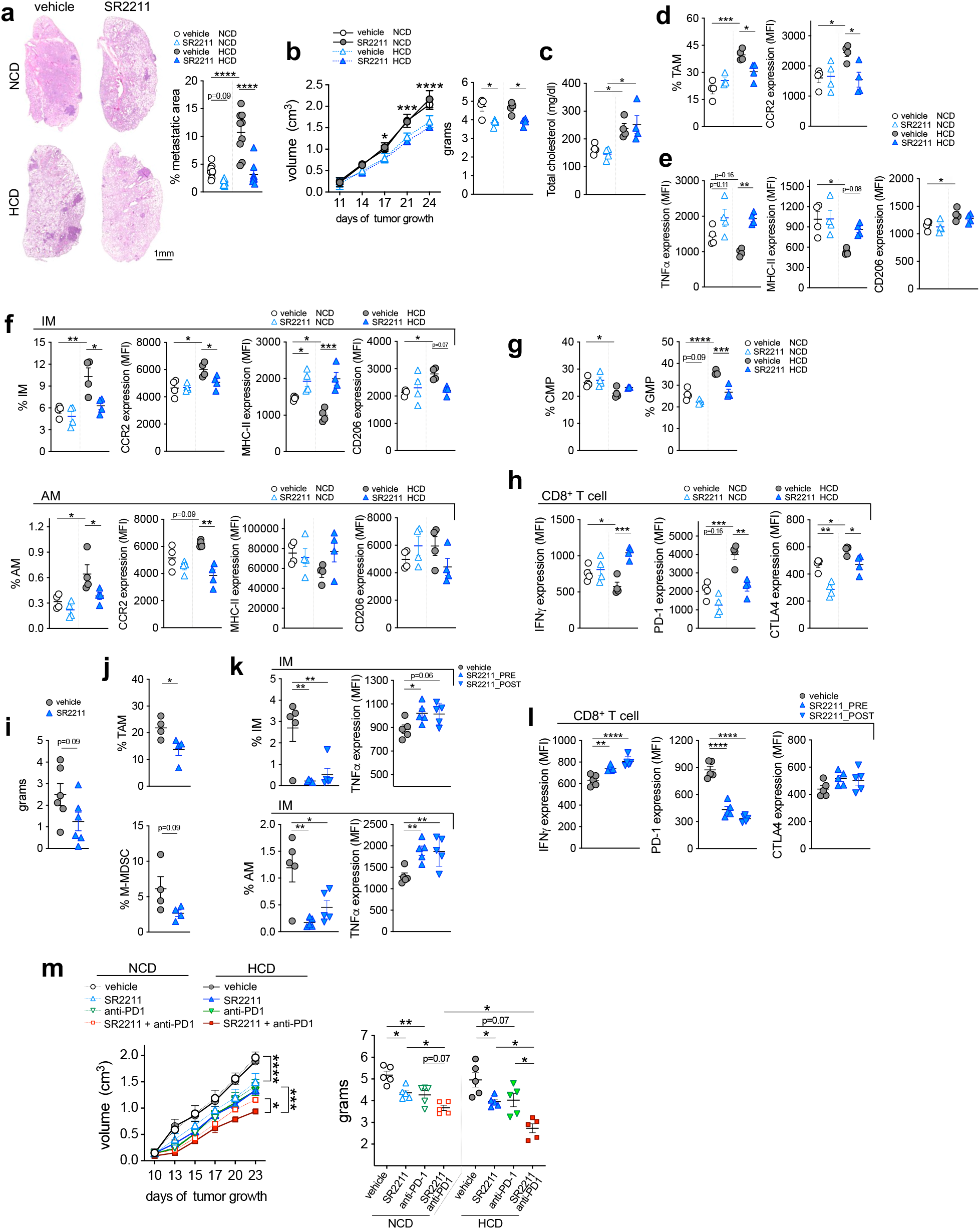
Targeting RORγ improves immunotherapy response. **a-h**, NCD- or HCD-fed MN/MCA1 wt mice treated with the RORγ inhibitor SR2211 or vehicle control (n=4). **a,** Lung metastatic area; representative images are shown, scale bar, 1 mm; **b,** tumor growth (volume and weight); **c,** total blood cholesterol; **d,** FACS quantification of TAMs and relative CCR2 expression; **e**, TNFα, MHC-II, and CD206 expression by TAMs (MFI); **f,** frequency of IM (top) and AM (bottom) subsets and relative expression (MFI) of CCR2, MHC-II, and CD206; **g,** CMP and GMP frequencies in BM; **h**, IFNγ, PD-1 and CTLA4 expression (MFI) in intratumoral CD8^+^ T cells. **i-l,** HCD-fed K1735-M2-bearing wt mice treated with SR2211 or vehicle control. **i,** Tumor weight (n=6); **j,** FACS quantification of TAMs and tumor-infiltrating M-MDSCs (n=4); **k,** IM and AM frequencies and relative TNFα expression (MFI) (n=5); **l,** IFNγ, PD1, and CTLA4 expression (MFI) by lung CD8^+^ T cells (n=5). **m,** Tumor growth (volume and weight) in NCD- or HCD-fed wt mice treated with vehicle, SR2211, anti-PD1, or SR2211 plus anti-PD1 (n=5). One experiment was performed. Data are expressed as mean ± SEM. \**P* < 0.05, \*\**P* < 0.01, \*\*\**P* < 0.001, \*\*\*\**P* < 0.0001 between selected relevant comparisons. **a, b** (right)**, c-h, k, l, m** (right), One-way ANOVA with Tukey’s multiple comparisons test. **i, j,** Unpaired two-tailed *t*-test. **b** (left)**, m** (left), Two-way ANOVA with Tukey’s multiple comparisons test

**Supplementary Fig. 1.**
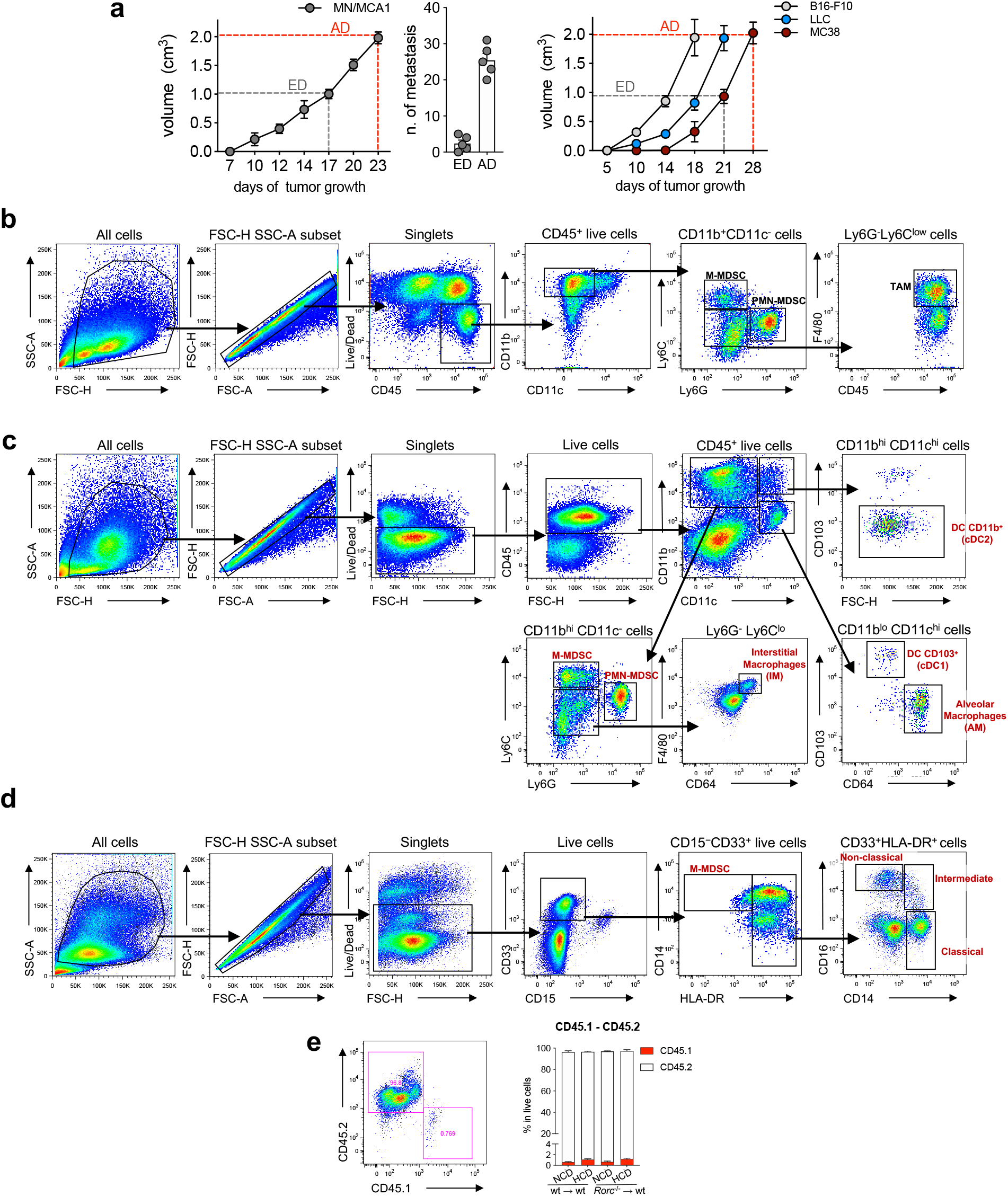
**a,** Left, representation of tumor progression in the early (ED) and advanced (AD) stages of the disease and (center) respective number of metastasis, in MN/MCA1-transplanted mice. Right, primary tumor growth in mice transplanted with B16-F10, LLC and MC38 cells. **b,** Gating strategy of myeloid subsets (M-MDSCs, PMN-MDSCs and TAMs) in murine cancers. **c,** Gating strategy of myeloid subsets (M- MDSCs, PMN-MDSCs, IMs, AMs, dendritic cells/DCs) in murine lungs. **d,** Gating strategy of human monocytic cell subsets (classical, intermediate, non-classical monocytes and M-MDSCs) in PBMCs. **e**, Check of BM reconstitution analyzing the percentage of blood CD45.2^+^ cells from donor mice *vs.* CD45.1^+^ cells of bone marrow depleted (lethally irradiated) recipent mice.

## References

1 Strauss, L., Guarneri, V., Gennari, A. & Sica, A. Implications of metabolism-driven myeloid dysfunctions in cancer therapy. Cell Mol Immunol 18, 829–841, doi:10.1038/s41423-020-00556-w (2021).

2 Jetten, A. M., Takeda, Y., Slominski, A. & Kang, H. S. Retinoic acid-related Orphan Receptor gamma (RORgamma): connecting sterol metabolism to regulation of the immune system and autoimmune disease. Curr Opin Toxicol 8, 66–80, doi:10.1016/j.cotox.2018.01.005 (2018).

3 Strauss, L. et al. RORC1 Regulates Tumor-Promoting “Emergency” Granulo-Monocytopoiesis. Cancer Cell 28, 253–269, doi:10.1016/j.ccell.2015.07.006 (2015).

4 Tall, A. R. & Yvan-Charvet, L. Cholesterol, inflammation and innate immunity. Nat Rev Immunol 15, 104–116, doi:10.1038/nri3793 (2015).

5 Swirski, F. K. et al. Ly-6Chi monocytes dominate hypercholesterolemia-associated monocytosis and give rise to macrophages in atheromata. J Clin Invest 117, 195–205, doi:10.1172/JCI29950 (2007).

6 Mantovani, A., Allavena, P., Sica, A. & Balkwill, F. Cancer-related inflammation. Nature 454, 436–444, doi:10.1038/nature07205 (2008).

7 Hegde, S., Leader, A. M. & Merad, M. MDSC: Markers, development, states, and unaddressed complexity. Immunity 54, 875–884, doi:10.1016/j.immuni.2021.04.004 (2021).

8 Guc, E. & Pollard, J. W. Redefining macrophage and neutrophil biology in the metastatic cascade. Immunity 54, 885–902, doi:10.1016/j.immuni.2021.03.022 (2021).

9 Condamine, T., Ramachandran, I., Youn, J. I. & Gabrilovich, D. I. Regulation of tumor metastasis by myeloid-derived suppressor cells. Annu Rev Med 66, 97–110, doi:10.1146/annurev-med-051013-052304 (2015).

10 Condamine, T., Mastio, J. & Gabrilovich, D. I. Transcriptional regulation of myeloid-derived suppressor cells. J Leukoc Biol 98, 913–922, doi:10.1189/jlb.4RI0515-204R (2015).

11 Porta, C. et al. Tumor-Derived Prostaglandin E2 Promotes p50 NF-kappaB-Dependent Differentiation of Monocytic MDSCs. Cancer Res 80, 2874–2888, doi:10.1158/0008-5472.CAN-19-2843 (2020).

12 Bleve, A., Durante, B., Sica, A. & Consonni, F. M. Lipid Metabolism and Cancer Immunotherapy: Immunosuppressive Myeloid Cells at the Crossroad. Int J Mol Sci 21, doi:10.3390/ijms21165845 (2020).

13 Hotamisligil, G. S. Inflammation, metaflammation and immunometabolic disorders. Nature 542, 177–185, doi:10.1038/nature21363 (2017).

14 Murray, P. J. Obesity corrupts myelopoiesis. Cell Metab 19, 735–736, doi:10.1016/j.cmet.2014.04.010 (2014).

15 Quail, D. F. et al. Obesity alters the lung myeloid cell landscape to enhance breast cancer metastasis through IL5 and GM-CSF. Nat Cell Biol 19, 974–987, doi:10.1038/ncb3578 (2017).

16 Iyengar, N. M., Hudis, C. A. & Dannenberg, A. J. Obesity and cancer: local and systemic mechanisms. Annu Rev Med 66, 297–309, doi:10.1146/annurev-med-050913-022228 (2015).

17 Kuzu, O. F., Noory, M. A. & Robertson, G. P. The Role of Cholesterol in Cancer. Cancer Res 76, 2063–2070, doi:10.1158/0008-5472.CAN-15-2613 (2016).

18 Avgerinos, K. I., Spyrou, N., Mantzoros, C. S. & Dalamaga, M. Obesity and cancer risk: Emerging biological mechanisms and perspectives. Metabolism 92, 121–135, doi:10.1016/j.metabol.2018.11.001 (2019).

19 Petrelli, F. et al. Association of Obesity With Survival Outcomes in Patients With Cancer: A Systematic Review and Meta-analysis. JAMA Netw Open 4, e213520, doi:10.1001/jamanetworkopen.2021.3520 (2021).

20 Bovenga, F., Sabba, C. & Moschetta, A. Uncoupling nuclear receptor LXR and cholesterol metabolism in cancer. Cell Metab 21, 517–526, doi:10.1016/j.cmet.2015.03.002 (2015).

21 Condamine, T. et al. Lectin-type oxidized LDL receptor-1 distinguishes population of human polymorphonuclear myeloid-derived suppressor cells in cancer patients. Sci Immunol 1, doi:10.1126/sciimmunol.aaf8943 (2016).

22 Goossens, P. et al. Membrane Cholesterol Efflux Drives Tumor-Associated Macrophage Reprogramming and Tumor Progression. Cell Metab 29, 1376–1389 e1374, doi:10.1016/j.cmet.2019.02.016 (2019).

23 Tavazoie, M. F. et al. LXR/ApoE Activation Restricts Innate Immune Suppression in Cancer. Cell 172, 825–840 e818, doi:10.1016/j.cell.2017.12.026 (2018).

24 Raccosta, L. et al. The oxysterol-CXCR2 axis plays a key role in the recruitment of tumor-promoting neutrophils. J Exp Med 210, 1711–1728, doi:10.1084/jem.20130440 (2013).

25 Di Conza, G. et al. Tumor-induced reshuffling of lipid composition on the endoplasmic reticulum membrane sustains macrophage survival and pro-tumorigenic activity. Nat Immunol 22, 1403–1415, doi:10.1038/s41590-021-01047-4 (2021).

26 Huang, B., Song, B. L. & Xu, C. Cholesterol metabolism in cancer: mechanisms and therapeutic opportunities. Nat Metab 2, 132–141, doi:10.1038/s42255-020-0174-0 (2020).

27 Luo, J., Yang, H. & Song, B. L. Mechanisms and regulation of cholesterol homeostasis. Nat Rev Mol Cell Biol 21, 225–245, doi:10.1038/s41580-019-0190-7 (2020).

28 Liu, X. et al. Inhibition of PCSK9 potentiates immune checkpoint therapy for cancer. Nature 588, 693–698, doi:10.1038/s41586-020-2911-7 (2020).

29 Groen, A. K., Bloks, V. W., Verkade, H. & Kuipers, F. Cross-talk between liver and intestine in control of cholesterol and energy homeostasis. Mol Aspects Med 37, 77–88, doi:10.1016/j.mam.2014.02.001 (2014).

30 Di Ciaula, A. et al. Bile Acid Physiology. Ann Hepatol 16 Suppl 1, S4–S14, doi:10.5604/01.3001.0010.5493 (2017).

31 Wang, D. Q. Regulation of intestinal cholesterol absorption. Annu Rev Physiol 69, 221–248, doi:10.1146/annurev.physiol.69.031905.160725 (2007).

32 DuPage, M., Dooley, A. L. & Jacks, T. Conditional mouse lung cancer models using adenoviral or lentiviral delivery of Cre recombinase. Nat Protoc 4, 1064–1072, doi:10.1038/nprot.2009.95 (2009).

33 Miyazawa, H. et al. Effect of Porphyromonas gingivalis infection on post-transcriptional regulation of the low-density lipoprotein receptor in mice. Lipids Health Dis 11, 121, doi:10.1186/1476-511X-11-121 (2012).

34 Lo, J. C. et al. Lymphotoxin beta receptor-dependent control of lipid homeostasis. Science 316, 285–288, doi:10.1126/science.1137221 (2007).

35 Frostegard, J., Ahmed, S., Hafstrom, I., Ajeganova, S. & Rahman, M. Low levels of PCSK9 are associated with remission in patients with rheumatoid arthritis treated with anti-TNF-alpha: potential underlying mechanisms. Arthritis Res Ther 23, 32, doi:10.1186/s13075-020-02386-7 (2021).

36 Tang, Z. H. et al. PCSK9: A novel inflammation modulator in atherosclerosis? J Cell Physiol 234, 2345–2355, doi:10.1002/jcp.27254 (2019).

37 Briukhovetska, D. et al. Interleukins in cancer: from biology to therapy. Nat Rev Cancer 21, 481–499, doi:10.1038/s41568-021-00363-z (2021).

38 Ferraz-Amaro, I. et al. Effect of IL-6 Receptor Blockade on Proprotein Convertase Subtilisin/Kexin Type-9 and Cholesterol Efflux Capacity in Rheumatoid Arthritis Patients. Horm Metab Res 51, 200–209, doi:10.1055/a-0833-4627 (2019).

39 Chang, C. S. et al. The relationship between pulse pressure with plasma PCSK9 and interleukin-6 among patients with acute ischemic stroke and dyslipidemia. Brain Res 1795, 148080, doi:10.1016/j.brainres.2022.148080 (2022).

40 Ding, Z. et al. NLRP3 inflammasome via IL-1beta regulates PCSK9 secretion. Theranostics 10, 7100–7110, doi:10.7150/thno.45939 (2020).

41 Pingili, A. K. et al. Immune checkpoint blockade reprograms systemic immune landscape and tumor microenvironment in obesity-associated breast cancer. Cell Rep 35, 109285, doi:10.1016/j.celrep.2021.109285 (2021).

42 Murray, P. J. et al. Macrophage activation and polarization: nomenclature and experimental guidelines. Immunity 41, 14–20, doi:10.1016/j.immuni.2014.06.008 (2014).

43 Consonni, F. M. et al. Heme catabolism by tumor-associated macrophages controls metastasis formation. Nat Immunol 22, 595–606, doi:10.1038/s41590-021-00921-5 (2021).

44 Baek, A. E. et al. The cholesterol metabolite 27 hydroxycholesterol facilitates breast cancer metastasis through its actions on immune cells. Nat Commun 8, 864, doi:10.1038/s41467-017-00910-z (2017).

45 Michelet, X. et al. Metabolic reprogramming of natural killer cells in obesity limits antitumor responses. Nat Immunol 19, 1330–1340, doi:10.1038/s41590-018-0251-7 (2018).

46 Pascual, G. et al. Targeting metastasis-initiating cells through the fatty acid receptor CD36. Nature 541, 41–45, doi:10.1038/nature20791 (2017).

47 Loyher, P. L. et al. Macrophages of distinct origins contribute to tumor development in the lung. J Exp Med 215, 2536–2553, doi:10.1084/jem.20180534 (2018).

48 Casanova-Acebes, M. et al. Tissue-resident macrophages provide a pro-tumorigenic niche to early NSCLC cells. Nature 595, 578–584, doi:10.1038/s41586-021-03651-8 (2021).

49 Wang, Z. et al. Paradoxical effects of obesity on T cell function during tumor progression and PD-1 checkpoint blockade. Nat Med 25, 141–151, doi:10.1038/s41591-018-0221-5 (2019).

50 Chan, J. C. et al. A proprotein convertase subtilisin/kexin type 9 neutralizing antibody reduces serum cholesterol in mice and nonhuman primates. Proc Natl Acad Sci U S A 106, 9820–9825, doi:10.1073/pnas.0903849106 (2009).

51 Jenkins, S. J. & Hume, D. A. Homeostasis in the mononuclear phagocyte system. Trends Immunol 35, 358–367, doi:10.1016/j.it.2014.06.006 (2014).

52 Bernelot Moens, S. J., et al. PCSK9 monoclonal antibodies reverse the pro-inflammatory profile of monocytes in familial hypercholesterolaemia. Eur Heart J 38, 1584–1593, doi:10.1093/eurheartj/ehx002 (2017).

53 Chen, Y. et al. The mouse CCR2 gene is regulated by two promoters that are responsive to plasma cholesterol and peroxisome proliferator-activated receptor gamma ligands. Biochem Biophys Res Commun 332, 188–193, doi:10.1016/j.bbrc.2005.04.110 (2005).

54 Kloudova, A., Guengerich, F. P. & Soucek, P. The Role of Oxysterols in Human Cancer. Trends Endocrinol Metab 28, 485–496, doi:10.1016/j.tem.2017.03.002 (2017).

55 Guillemot-Legris, O., Mutemberezi, V., Cani, P. D. & Muccioli, G. G. Obesity is associated with changes in oxysterol metabolism and levels in mice liver, hypothalamus, adipose tissue and plasma. Sci Rep 6, 19694, doi:10.1038/srep19694 (2016).

56 Tremblay-Franco, M. et al. Effect of obesity and metabolic syndrome on plasma oxysterols and fatty acids in human. Steroids 99, 287–292, doi:10.1016/j.steroids.2015.03.019 (2015).

57 Singh, A. K. et al. SUMOylation of ROR-gammat inhibits IL-17 expression and inflammation via HDAC2. Nat Commun 9, 4515, doi:10.1038/s41467-018-06924-5 (2018).

58 Kumar, N. et al. Identification of SR2211: a potent synthetic RORgamma-selective modulator. ACS Chem Biol 7, 672–677, doi:10.1021/cb200496y (2012).

59 Xi, Y. et al. Mechanisms of induction of tumors by cholesterol and potential therapeutic prospects. Biomed Pharmacother 144, 112277, doi:10.1016/j.biopha.2021.112277 (2021).

60 Bielecka-Dabrowa, A., Hannam, S., Rysz, J. & Banach, M. Malignancy-associated dyslipidemia. Open Cardiovasc Med J 5, 35–40, doi:10.2174/1874192401105010035 (2011).

61 Agustsson, T. et al. Mechanism of increased lipolysis in cancer cachexia. Cancer Res 67, 5531–5537, doi:10.1158/0008-5472.CAN-06-4585 (2007).

62 Doroshow, D. B. et al. PD-L1 as a biomarker of response to immune-checkpoint inhibitors. Nat Rev Clin Oncol 18, 345–362, doi:10.1038/s41571-021-00473-5 (2021).

63 McQuade, J. L. et al. Association of body-mass index and outcomes in patients with metastatic melanoma treated with targeted therapy, immunotherapy, or chemotherapy: a retrospective, multicohort analysis. Lancet Oncol 19, 310–322, doi:10.1016/S1470-2045(18)30078-0 (2018).

## Supplementary References

64 Kraus, D., Yang, Q. & Kahn, B. B. Lipid Extraction from Mouse Feces. Bio Protoc 5, doi:10.21769/bioprotoc.1375 (2015).

65 Charni-Natan, M. & Goldstein, I. Protocol for Primary Mouse Hepatocyte Isolation. STAR Protoc 1, 100086, doi:10.1016/j.xpro.2020.100086 (2020).

66 Honda, A. et al. Highly sensitive assay of HMG-CoA reductase activity by LC-ESI-MS/MS. J Lipid Res 48, 1212–1220, doi:10.1194/jlr.D600049-JLR200 (2007).

67 Masini, M. A. et al. Prolonged exposure to simulated microgravity promotes stemness impairing morphological, metabolic and migratory profile of pancreatic cancer cells: a comprehensive proteomic, lipidomic and transcriptomic analysis. Cell Mol Life Sci 79, 226, doi:10.1007/s00018-022-04243-z (2022).

68 Tishchenko, I., Milioli, H. H., Riveros, C. & Moscato, P. Extensive Transcriptomic and Genomic Analysis Provides New Insights about Luminal Breast Cancers. PLoS One 11, e0158259, doi:10.1371/journal.pone.0158259 (2016).

69 Dobin, A. et al. STAR: ultrafast universal RNA-seq aligner. Bioinformatics 29, 15–21, doi:10.1093/bioinformatics/bts635 (2013).

70 Robinson, M. D., McCarthy, D. J. & Smyth, G. K. edgeR: a Bioconductor package for differential expression analysis of digital gene expression data. Bioinformatics 26, 139–140, doi:10.1093/bioinformatics/btp616 (2010).

71 Ritchie, M. E. et al. limma powers differential expression analyses for RNA-sequencing and microarray studies. Nucleic Acids Res 43, e47, doi:10.1093/nar/gkv007 (2015).

72 Kolde, R. Pheatmap: pretty heatmaps. R package version 1, 726 (2012).

73 de Tayrac, M., Le, S., Aubry, M., Mosser, J. & Husson, F. Simultaneous analysis of distinct Omics data sets with integration of biological knowledge: Multiple Factor Analysis approach. BMC Genomics 10, 32, doi:10.1186/1471-2164-10-32 (2009).

74 Conway, J. R., Lex, A. & Gehlenborg, N. UpSetR: an R package for the visualization of intersecting sets and their properties. Bioinformatics 33, 2938–2940, doi:10.1093/bioinformatics/btx364 (2017).

75 Kuleshov, M. V. et al. Enrichr: a comprehensive gene set enrichment analysis web server 2016 update. Nucleic Acids Res 44, W90–97, doi:10.1093/nar/gkw377 (2016).

