## Supplementary Figure 1 for "RORγ bridges cancer-driven lipid dysmetabolism and myeloid immunosuppression"

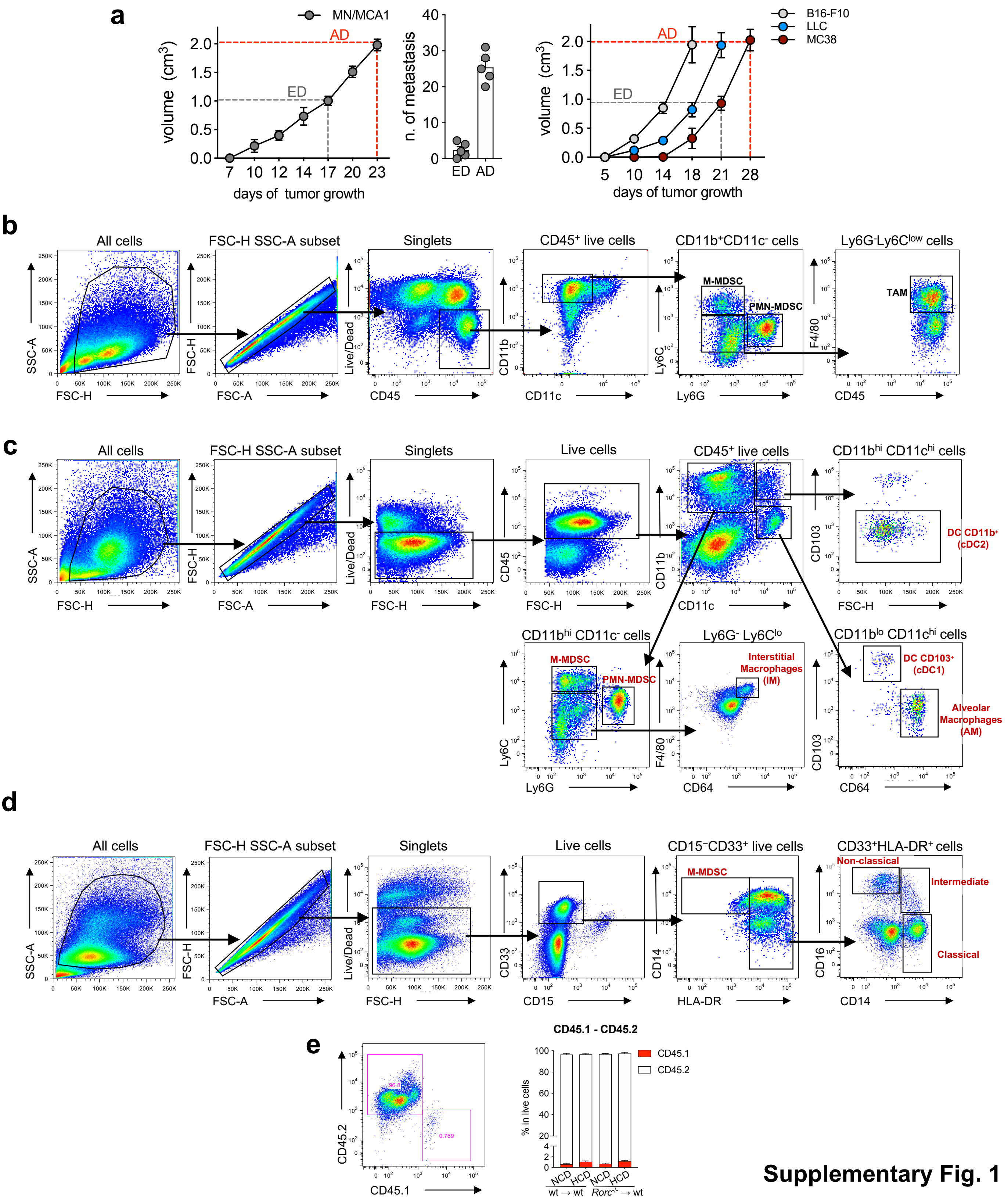


**Supplementary Fig. 1**

**Supplementary Fig. 1. a,** Left, representation of tumor progression in the early (ED) and advanced (AD) stages of the disease and (center) respective number of metastasis, in MN/MCA1-transplanted mice. Right, primary tumor growth in mice transplanted with B16-F10, LLC and MC38 cells. **b,** Gating strategy of myeloid subsets (M-MDSCs, PMN-MDSCs and TAMs) in murine cancers. **c,** Gating strategy of myeloid subsets (M-MDSCs, PMN-MDSCs, IMs, AMs, dendritic cells/DCs) in murine lungs. **d,** Gating strategy of human monocytic cell subsets (classical, intermediate, non-classical monocytes and M-MDSCs) in PBMCs. **e**, Check of BM reconstitution analyzing the percentage of blood CD45.2$^{+}$ cells from donor mice *vs.* CD45.1$^{+}$ cells of bone marrow depleted (lethally irradiated) recipent mice.
