## Supplementary Table 2 for "RORγ bridges cancer-driven lipid dysmetabolism and myeloid immunosuppression"

| Group | Monocytic cell subset | Total Cholesterol | | HDL | | LDL | |
|---|---|---|---|---|---|---|---|
| | | Pearson *r* | p value | Pearson *r* | p value | Pearson *r* | p value |
| **Healthy Donors (HD, n=18)** | Classical | 0.02419 | 0.9118 | -0.6403 | 0.0042 | 0.2339 | 0.3502 |
| | Classical RORγ$^+$ | 0.3546 | 0.1488 | 0.1850 | 0.4623 | 0.3315 | 0.1790 |
| | Intermediate | -0.4096 | 0.0914 | 0.2849 | 0.2518 | -0.4833 | 0.0422 |
| | Intermediate RORγ$^+$ | 0.6901 | 0.0015 | -0.4644 | 0.0522 | 0.7977 | <0.0001 |
| | Non-classical | 0.02261 | 0.9290 | -0.3008 | 0.2252 | 0.1023 | 0.6864 |
| | Non-classical RORγ$^+$ | 0.3661 | 0.1351 | -0.5747 | 0.0126 | 0.5329 | 0.0228 |
| | MDSC | 0.2330 | 0.3522 | -0.2511 | 0.3148 | 0.3010 | 0.2249 |
| | MDSC RORγ$^+$ | 0.4068 | 0.0939 | -0.3128 | 0.2063 | 0.4890 | 0.0394 |
| **NSCLC (CA, n=23)** | Classical | -0.3808 | 0.0730 | -0.1998 | 0.3607 | -0.4841 | 0.0192 |
| | Classical RORγ$^+$ | 0.009384 | 0.9661 | 0.1658 | 0.4496 | -0.02231 | 0.9195 |
| | Intermediate | 0.08649 | 0.6948 | -0.08488 | 0.7002 | -0.1479 | 0.5006 |
| | Intermediate RORγ$^+$ | -0.09342 | 0.6716 | 0.2050 | 0.3482 | -0.006684 | 0.9759 |
| | Non-classical | 0.2518 | 0.2465 | 0.03308 | 0.8809 | 0.001560 | 0.9944 |
| | Non-classical RORγ$^+$ | 0.007072 | 0.9745 | -0.4516 | 0.4912 | -0.06257 | 0.7767 |
| | MDSC | 0.5606 | 0.0054 | 0.0305 | 0.0305 | 0.4340 | 0.0385 |
| | MDSC RORγ$^+$ | 0.4206 | 0.0457 | 0.2863 | 0.2863 | 0.4291 | 0.0411 |

**Supplementary Table 3**. Pearson correlation coefficient (*r*) and statistical analysis (p value) between the frequency of monocytic and RORγ$^+$ cell subsets (classical, HLA-DR$^+$CD14$^+$CD16$^-$; intermediate, HLA-DR$^+$CD14$^+$CD16$^+$; non-classical, HLA-DR$^+$CD14$^-$CD16$^+$; myeloid-derived suppressor cells, MDSC, HLA-DR$^-$CD14$^+$) and blood cholesterol levels.
