## Supplementary Tables 4 and 5 for "RORγ bridges cancer-driven lipid dysmetabolism and myeloid immunosuppression"

Supplementary Table 4. Murine antibodies for flow cytometry analysis

| **Marker** | **Fluorophore** | **Clone** | **Titer** | **Company** | **n. catalog** |
|---|---|---|---|---|---|
| CD45 | FITC | 30-F11 | 1:400 | Biolegend | 103108 |
| | PerCP | 30-F11 | 1:200 | Biolegend | 103130 |
| | BUV563 | 30-F11 | 1:400 | BD Horizon | 565710 |
| CD45.1 | APC | A20 | 1:200 | Biolegend | 110720 |
| CD45.2 | PerCP-Cy5.5 | 104 | 1:200 | Biolegend | 109827 |
| CD11b | PerCP-Cy5.5 | M1/70 | 1:200 | eBioscience | 45-0112-82 |
| | PE | M1/70 | 1:400 | Biolegend | 101208 |
| | FITC | M1/70 | 1:200 | Biolegend | 101206 |
| | BV711 | M1/70 | 1:600 | Biolegend | 101242 |
| F4/80 | APC | BM8 | 1:100 | Biolegend | 123116 |
| | PE | BM8 | 1:200 | Biolegend | 123110 |
| | PE-Cy7 | BM8 | 1:200 | Biolegend | 123114 |
| | BV421 | BM8 | 1:100 | Biolegend | 123137 |
| Ly6G | PE | 1A8 | 1:200 | Biolegend | 127608 |
| | BV570 | 1A8 | 1:100 | Biolegend | 127629 |
| Ly6C | APC-Cy7 | HK1.4 | 1:600 | eBioscience | 47-5932-82 |
| | APC | HK1.4 | 1:600 | Biolegend | 128016 |
| TNFα | Alexa Fluor 647 | MP6-XT22 | 1:100 | BD Pharmingen | 557730 |
| | PE-Cy7 | MP6-XT22 | 1:100 | BD Pharmingen | 561041 |
| MHCII | PE-Cy7 | M5/114.15.2 | 1:200 | eBioscience | 78-5321-82 |
| | PE | M5/114.15.2 | 1:400 | Biolegend | 107607 |
| | BV480 | M5/114.15.2 | 1:600 | BD Horizon | 566088 |
| iNOS | PE-eFluor 610 | CXNFT | 1:100 | eBioscience | 61-592082 |
| CCR7 | PE | 4B12 | 1:200 | eBioscience | 12-1971-82 |
| CD206 | APC | C068C2 | 1:100 | Biolegend | 141708 |
| | PE/Dazzle-594 | C068C2 | 1:200 | Biolegend | 141731 |
| PD-L1 | PE | 10F.9G2 | 1:100 | Biolegend | 124308 |
| CD204 | APC-Vio770 | REA148 | 1:200 | Miltenyi | 130-105-575 |
| | PE-Vio770 | REA148 | 1:200 | Miltenyi | 130-105-574 |
| IDO1 | PE | mIDO-48 | 1:100 | eBioscience | 121-9473-80 |
| CD11c | APC | N418 | 1:100 | Biolegend | 117310 |
| | BV605 | N418 | 1:100 | eBioscience | 63-0114-82 |
| CD64 | APC | X54-5/7.1 | 1:100 | eBioscience | 17-0641-82 |
| CD103 | PE-CF594 | M290 | 1:100 | BD Horizon | 565849 |
| CD8 | APC | 53-6.7 | 1:100 | Biolegend | 100712 |
| | BUV805 | 53-6.7 | 1:200 | BD Horizon | 564920 |
| | PE | 53-6.7 | 1:200 | Biolegend | 100708 |
| | PE-Cy7 | 53-6.7 | 1:200 | Biolegend | 100721 |
| | PE-Cy5 | 53-6.7 | 1:200 | eBioscience | 15-0081-82 |
| CD4 | PE | GK1.5 | 1:200 | Biolegend | 100408 |
| | PE-Cy7 | GK1.5 | 1:200 | Biolegend | 100422 |
| | BUV496 | GK1.5 | 1:100 | BD Horizon | 564667 |
| | APC | GK1.5 | 1:200 | Biolegend | 100412 |
| CD3 | PerCP | 145-2C11 | 1:100 | Biolegend | 100326 |
| | BV650 | 145-2C11 | 1:100 | Biolegend | 100229 |
| PD-1 | PE-Cy7 | 29F.1A12 | 1:100 | Biolegend | 135215 |
| | PE | 29F.1A12 | 1:100 | Biolegend | 135205 |
| CTLA4 | APC | UC10-4B9 | 1:100 | Biolegend | 106309 |
| | BV421 | UC10-4B9 | 1:100 | Biolegend | 106311 |

| | | | | | |
|---|---|---|---|---|---|
| IFNγ | FITC | XMG1.2 | 1:100 | Biolegend | 505806 |
| | PE | XMG1.2 | 1:200 | BD Pharmingen | 554412 |
| | Pacific Blue | XMG1.2 | 1:100 | Biolegend | 505818 |
| CD31 | PE | MEC 13.3 | 1:100 | BD Pharmingen | 553373 |
| CCR2 | PE-Cy7 | SA203G11 | 1:200 | Biolegend | 150612 |
| CD115 | PE | AFS98 | 1:100 | Biolegend | 135505 |
| | APC | AFS98 | 1:100 | Biolegend | 135510 |
| Hematopoietic Lineage | eFluor 450 | NA | 1:400 | eBioscience | 88-7772-72 |
| c-Kit | PE-Cy7 | 2B8 | 1:200 | Biolegend | 105814 |
| CD16/32 | APC | 93 | 1:100 | eBioscience | 17-0161-82 |
| | PE | 2.4G2 | 1:100 | BD Pharmingen | 553145 |
| | APC-Cy7 | 93 | 1:200 | Biolegend | 101327 |
| CD34 | FITC | RAM34 | 1:100 | BD Pharmingen | 560238 |
| | BV605 | RAM34 | 1:200 | BD Optibuild | 750918 |
| RORγ | APC | AFKJS | 1:100 | eBioscience | 17-6988-80 |
| | PE | AFKJS | 1:100 | eBioscience | 17-6988-82 |
| | BV421 | Q31-378 | 1:100 | BD Horizon | 562894 |
| PCSK9 | unconjugated | 2F1 | 1:100 | Invitrogen | MA5-32843 |
| LDLR | unconjugated | SJ0197 | 1:100 | Invitrogen | MA5-32075 |

### Supplementary Table 5. Human antibodies for flow cytometry analysis

| **Marker** | **Fluorophore** | **Clone** | **Titer** | **Company** | **n. catalog** |
|---|---|---|---|---|---|
| CD33 | PerCP-Cy5.5 | WM53 | 1:80 | Biolegend | 303414 |
| CD15 | FITC | SSEA-1 | 1:100 | Biolegend | 394706 |
| HLA-DR | Pacific Blue | L243 | 1:80 | Biolegend | 307624 |
| CD14 | PE-Cy7 | M5E2 | 1:150 | Biolegend | 301814 |
| CD16 | BV605 | 3G8 | 1/150 | Biolegend | 302040 |
| CCR2 | APC | 48607 | 1:100 | R&D systems | FAB151A |
| RORγ | PE | AFKJS | 1:100 | eBioscience | 17-6988-82 |
